# Molecular mechanism of naturally-encoded signaling-bias at the complement anaphylatoxin receptors

**DOI:** 10.1101/2025.11.01.685996

**Authors:** Divyanshu Tiwari, Kazuhiro Sawada, Annu Dalal, Sudha Mishra, Xaria X. Li, Joshua C. Dent, Kiae Kim, Manish K. Yadav, Nabarun Roy, Manisankar Ganguly, Nilanjana Banerjee, Tomasz Maciej Stepniewski, Donghoon Ahn, Kohei Yamaguchi, Hidetaka S. Oshima, Kana Hashimoto, Jenny N. Fung, Titaya Lerskiatiphanich, Cedric S. Cui, John D. Lee, Jana Selent, Asuka Inoue, Richard Clark, Ka Young Chung, Ramanuj Banerjee, Fumiya K. Sano, Trent M. Woodruff, Osamu Nureki, Arun K. Shukla

**Affiliations:** Department of Biological Sciences and Bioengineering, Indian Institute of Technology, Kanpur 208016, India; Department of Biological Sciences, Graduate School of Science, The University of Tokyo, Tokyo, Japan; School of Biomedical Sciences, Faculty of Health, Medicine, and Behavioural Sciences, The University of Queensland, Brisbane, QLD 4072, Australia; School of Pharmacy, Sungkyunkwan University, Suwon 16419, Republic of Korea; Research Programme on Biomedical Informatics (GRIB), Hospital del Mar Medical Research Institute & Pompeu Fabra University, Barcelona, Spain; Graduate School of Pharmaceutical Sciences, Tohoku University, Sendai, Miyagi, 980-8578, Japan; Graduate School of Pharmaceutical Sciences, Kyoto University, Kyoto 606-8501, Japan

**Keywords:** GPCRs, ACRs, signaling-bias, β-Arrestins, anaphylatoxin receptors, complement cascade, drug discovery

## Abstract

The conceptual framework of biased signaling has revolutionized our understanding of GPCR signaling and regulatory paradigms, and greatly impacted the efforts focused on the discovery of GPCR-targeted therapeutics. However, the mechanistic basis of biased signaling remains primarily defined based on synthetic ligands and receptor mutants with relatively limited progress in understanding naturally-encoded signaling-bias. Here, we present fundamental molecular and structural insights into naturally-encoded signaling-bias at the complement anaphylatoxin C5a receptors namely, C5aR1 and C5aR2. We first discover that C5a^-d-Arg^, the naturally-occurring version of C5a lacking the terminal arginine, exhibits robust G-protein signaling-bias at C5aR1, characterised by attenuated βarr recruitment. This signaling-bias manifests in both cytokine release from primary human immune cells, and *in vivo,* during neutrophil mobilization. We combine the cryo-EM structures of C5a/C5a^-d-Arg^-C5aR1 complexes with MD simulation, site-directed mutagenesis, and cellular experiments to elucidate that the G-protein-bias exhibited by C5a^-d-Arg^ results from a distinct orientation of TM7 and helix 8 in C5aR1 leading to inefficient GRK recruitment and receptor phosphorylation. Next, we determine the first cryo-EM structures of C5aR2, a naturally-encoded β-arrestin-biased receptor, in an apo state, complexed with the natural agonists C5a and C5a^-d-Arg^, and three peptide agonists including a first-in-class, newly discovered C5aR2-selective agonist, R8Y. These structural snapshots reveal key differences between the binding of C5a and C5a^-d-Arg^ to C5aR1 and C5aR2, and provide a molecular basis of functional specialization at these two receptors. Moreover, the structural insights also allow us to decipher the molecular basis of naturally-encoded signaling-bias at C5aR2 originating from a shallower cytoplasmic interface with hydrophobic interior pocket that is not permissive to efficient G-protein-coupling and activation. Finally, we also engineer and characterize loss-of-function and gain-of-function variants of C5aR1 and C5aR2, which in turn corroborate and validate the structural observations presented here. Collectively, our findings offer crucial insights into previously lacking molecular mechanisms of the naturally-encoded signaling-bias at GPCRs, which have broad implications not only for the general framework of biased-signaling, but also for novel therapeutic design.

## Introduction

G protein-coupled receptors (GPCRs) constitute a large superfamily of seven transmembrane receptors (7TMRs) with a direct involvement in a broad array of physiological and pathophysiological processes, which also makes them highly sought-after drug targets^1,2^. Upon agonist-stimulation, GPCRs typically couple to, and signal through, heterotrimeric G-proteins and β-arrestin (βarrs), and it has been possible to design synthetic ligands capable of preferentially activating one of these transducers^3,4^. These ligands are known as biased-agonists and the phenomenon of preferential activation of a specific transducer as biased agonism, and their therapeutic promise has brought about a paradigm change in GPCR-targeted novel drug discovery^5–7^. The conceptual framework of biased-signaling is based primarily on synthetic ligands and receptor mutants for the most commonly studied systems, and the examples of naturally-encoded biased ligands are rather limited^4,5,7–10^. This represents a key knowledge gap and poses an important caveat in fully appreciating the physiological implications of this therapeutically important paradigm. Interestingly, the complement anaphylatoxins and their cognate receptors encoded in the complement cascade constitute an intriguing system to probe the naturally-encoded ligand-induced and receptor-mediated signaling-bias^11–14^.

The complement cascade plays a critical role in the innate immune response mechanisms, especially in the complex landscape of host-pathogen interactions, as a protective mechanism^15,16^. The potent anaphylatoxins referred to as C3a and C5a are generated in the final steps by the proteolytic cleavage of complement proteins C3 and C5, and they activate three different 7TMRs known as C3aR, C5aR1 and C5aR2 to exert the functional outcomes such as chemotaxis, degranulation, and cytokine production^17–19^. These receptors are expressed by a variety of immune cells such as macrophages and neutrophils, and upon activation by the corresponding anaphylatoxins, mediate a wide array of functional responses^14,20–24^. Their excessive and sustained activation is often associated with multiple pathophysiological conditions including sepsis, autoimmune disorders, rheumatoid arthritis, and multiple sclerosis, making them important drug targets for novel therapeutics^14,23,25–30^. Interestingly, the terminal arginine residue in C3a and C5a are cleaved by carboxypeptidases to generate C3a^-d-Arg^ and C5a^-d-Arg^, respectively, and it is commonly believed to be a mechanism to dampen the inflammatory response as the terminal arginine is critical for the binding of C3a/C5a to the corresponding receptors^31–34^. However, there are indications in the literature that C5a^-d-Arg^ may still bind to C5aR1 and C5aR2 and exert functional responses^23^. Therefore, a systematic and comprehensive exploration of C3a^-d-Arg^/C5a^-d-Arg^ interaction with their cognate receptors is essential to resolve uncertainties regarding their functional and physiological relevance. Moreover, a molecular understanding of these interactions guide the design of receptor subtype-selective ligands, especially for C5aR2 which currently lacks potent tool compounds, and help segregate overlapping and distinct functions of C5aR1 and C5aR2^24^.

There are a set of 7TMRs that lack functional G-protein-coupling despite having an overall architecture similar to GPCRs, and these are classified as Atypical Chemokine Receptors (ACKRs) as they recognize chemokines as their natural agonists^35–39^. While four of these receptors namely ACKR2-5 couple to βarrs, one of these, ACKR1, also known as the Duffy antigen receptor for chemokines (DARC), lacks a measurable coupling to βarrs as well^40^. The sub-family of these so called non-canonical GPCRs is further expanded by C5aR2, which also lacks functional G-protein-coupling but maintains robust βarr recruitment despite being activated by C5a and C5a^-d-Arg12^. Taken together, these five receptors, i.e., ACKR2-5 and C5aR2 constitute a sub-family referred to as Arrestin-Coupled Receptors (ACRs), and present an excellent system to study the intricacies of naturally-encoded signaling-bias at the receptor level. In particular, most of these ACRs share a natural agonist with a prototypical GPCR, thereby presenting a GPCR-ACR pair to directly compare the commonalities and differences in their ligand binding, conformational changes, and transducer-coupling to elucidate the key principles of biased signaling^41^.

Here, we demonstrate using a combination of cellular, biochemical, and pharmacological approaches that C5a^-d-Arg^ acts as a robust G-protein-biased agonist at C5aR1, and the transducer-coupling-bias is linked to distinct cellular and functional outcomes. We present the cryo-EM structures of C5a^-d-Arg^-bound C5aR1 and combine the structural insights with biochemical experiments to uncover the molecular mechanism driving the signaling-bias. Moreover, we also determine the first cryo-EM structures of the C5aR2 in apo-state, in complex with C5a, and a set of peptide agonists including a first-in-class, C5aR2-selective agonist. These structural snapshots elucidate the molecular basis of intrinsic signaling-bias encoded at C5aR2, and also uncover the design principles that allow us to engineer sub-type selective and signaling-biased C5a variants at C5aR1 and C5aR2.

## Results

### C5a^-d-Arg^ is a naturally-encoded biased agonist at C5aR1

As reported previously^42^ and reproduced here (**Figure 1A-C**, **S1A-H**), we observed that contrary to the broadly presented notion in the literature^33,43,44^, C5a^-d-Arg^ activates G-proteins with potency and efficacy nearly indistinguishable from C5a while it is substantially attenuated in βarr recruitment. The calculation of bias-factor further corroborates these observations, and establishes C5a^-d-Arg^ as a G-protein-biased agonist at C5aR1. Similar to the data observed in HEK-293T cells, we observed an equivalent Ca^2+^ response for both C5a and C5a^-d-Arg^ in primary human monocyte derived macrophages (HMDMs) and mouse bone marrow derived macrophages (BMDMs) (**Figure 1D-E)**. However, ERK1/2 phosphorylation in HEK-293, HMDMs, and BMDMs, and RhoA activation in HEK-293 cells was significantly attenuated for C5a^-d-Arg^ compared to C5a (**Figure 1F-I, S1I-J, S1K-L**). Interestingly, C5a and C5a^-d-Arg^ elicited a similar response in terms of IL-8 release from human macrophages (**Figure 1J**) while the bell-shaped dose-response typically observed for C5a in the PMN (polymorphonuclear leukocytes) migration assay^45^ was not observed for C5a^-d-Arg^ (**Figure 1K**). Moreover, plasma-purified C5a^-d-Arg^ recapitulated a similar pattern as that of the recombinant C5a^-d-Arg^ in terms of Ca^2+^ influx in HMDMs (**Figure 1L**), βarr2 recruitment in HEK-293 cells measured by BRET-assay (**Figure 1M**), pERK1/2 phosphorylation in HMDM **(Figure 1N)** and BMDMs (**Figure 1O**), IL-8 release (**Figure 1P**), and PMN migration (**Figure 1Q).** In order to further link these *in vitro* observations with an *in vivo* readout, we administered C5a and C5a^-d-Arg^ in mice to induce bone marrow neutrophil mobilization into the blood. We observed a significant reduction in neutrophil blood mobilization for C5a^-d-Arg^ compared to C5a (**Figure 1R**), which aligns with a functional-bias encoded by C5a^-d-Arg^. Interestingly, pre-dosing mice with the C5aR1-selctive antagonist PMX205^46^, followed by the administration of C5a and C5a^-d-Arg^, reduced neutrophil blood mobilization as expected (due to blockade of C5aR1), but more importantly, there was no apparent difference between C5a vs. C5a^-d-Arg^ (**Figure 1R**). These data indicate that the attenuation of βarr recruitment exhibited by C5a^-d-Arg^ translates into a functional response (i.e., neutrophil mobilization) *in vivo*, and that the residual response after C5aR1 blockade likely arises from the second C5a receptor, C5aR2, which is discussed in a subsequent section. Taken together, these data corroborate an intrinsic functional-bias encoded by C5a^-d-Arg^ at the human and mouse C5aR1, and also establish it as one of the very few naturally-encoded biased agonists identified till date.

**Figure 1:**
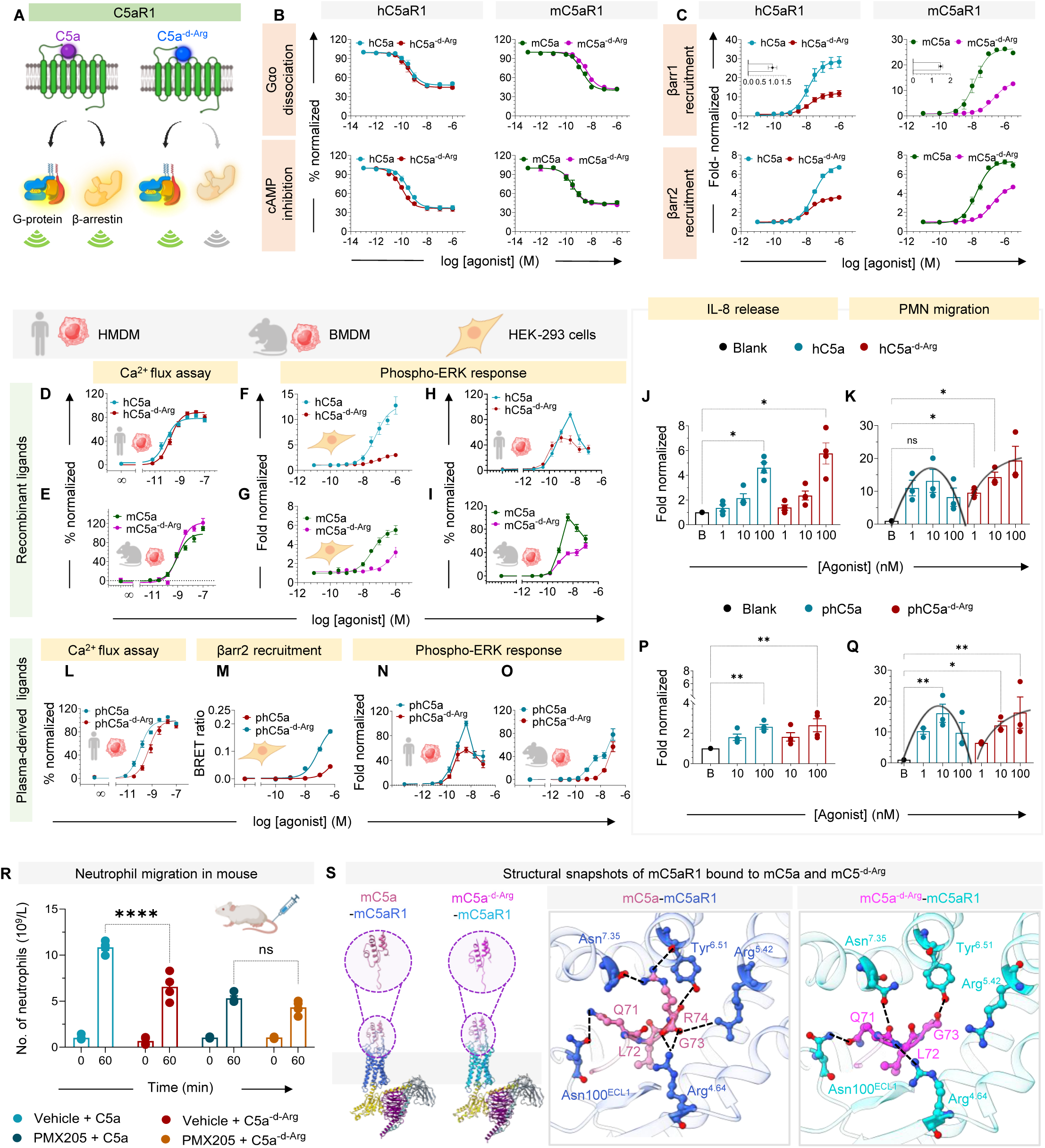
Naturally encoded ligand bias at complement anaphylatoxin receptor for C5a (C5aR1) **(A)** Schematic illustration representing C5a^-d-Arg^ mediated ligand bias at C5aR1. Schematic was prepared in <u>BioRender</u>. **(B)** Heterotrimeric GoA dissociation (upper panel) and cAMP inhibition (lower panel) was measured as a functional readout of human (h) and mouse (m) C5aR1 activation by (h/m) C5a and (h/m) C5a^-d-Arg^, employing NanoBiT-based enzyme complementation assay and GloSensor assay, respectively. Data (mean ± SEM) represent n=3-4 independent experiments. **(C)** âarr1/2 recruitment downstream to (h/m) C5aR1 in response to (h/m) C5a and (h/m) C5a^-d-Arg^ measured by NanoBiT assay. Data (mean ± SEM) represent n=3 independent experiments, fold normalized with respect to response recorded for lowest ligand dose. The inset within the upper panel displays the bias plot representing the G-protein bias encoded by C5a^-d-Arg^. C5aR1 mediated signaling in response to recombinant **(D-K)** and human plasma-derived **(L-Q)** C5a and C5a^-d-Arg^ showing C5a^-d-Arg^ preferentially mediates G-protein signaling upon activation. **(D-E)** Ligand induced intracellular calcium mobilization in human monocyte derived macrophages (HMDMs) (upper panel, in response to hC5a and hC5a^-d-Arg^) and mouse bone marrow derived macrophage (BMDM) (lower panel, in response to mC5a and mC5a^-d-Arg^) was monitored using the calcium dye Fluo-4 NW for 100 seconds, with ligand added at 16 seconds. Changes in fluorescence were normalized to the maximum ligand-induced response. Data represent mean ± SEM (n=3), independent donors. **(F-I)** ERK phosphorylation using SRE-based reporter assay in HEK-293 cells **(F-G)** and AlphaLISA Surefire-Ultra p-ERK1/2 kit in HMDM **(H)** /BMDM **(I)**. Data was fold-normalized to the lowest ligand concentration for SRF-RE reporter assay. Data represent mean ± SEM (n≥4), independent experiments. **(J)** IL-8 release in HMDMs in response to hC5a and hC5a^-d-Arg^. Cells were incubated with respective ligands for 24 h prior to supernatant collection. Data was analysed using repeated measures one-way ANOVA followed by Dunnette’s post-hoc test, comparing each condition to the unstimulated control (Blank). Significant IL-8 release were observed at 100 nM for both hC5a (adjusted p=0.0106) and hC5a^-d-Arg^ (adjusted p= 0.0352). Data represent mean ± SEM (n≥4), independent experiments. **(K)** Ligand-induced human polymorphonuclear neutrophils (PMN) chemotaxis was assessed using the Corning® FluoroBlok system, with migration quantified at 20 minutes post-ligand addition and normalized to the medium-only treated cells (fold-baseline, n = 3 independent donors) (ANOVA, p < 0.05*, p < 0.01**, p < 0.001***). **(L)** Ligand induced Calcium mobilization in HMDM in response to plasma-derived hC5a and hC5a^-d-Arg^. **(M)** βarr2 recruitment in response to increasing ligand concentration downstream to C5aR1 measured by BRET-based assay in HEK-293 cells. The ligand-induced BRET ratio following ligand stimulation at 40 minutes, normalized with respect to that of lowest ligand dose. **(N-O)** ERK-phosphorylation assay (similar to panel H-I) in HMDM **(N)** and BMDM **(O)** in response to varying ligand doses. Data represent mean ± SEM (n ≥ 4), independent experiments. **(P-Q)** IL-8 release **(P)** and PMN migration **(Q)** in response to plasma-derived hC5a and hC5a^-d-Arg^ (undertaken similar to panel J-K, respectively). Data represent fold-baseline, n = 3 independent donors (ANOVA, p < 0.05*, p < 0.01**, p < 0.001***) **(R)** Neutrophil migration in mouse in response to C5a and C5a^-d-Arg^ in the absence and presence of C5aR1-inhibitor, PMX205, measured post 60 minutes ligand injection to mouse. Data represent n=4 independent experiments, and analysis was carried out by using two-way ANOVA, Tukey’s multiple comparisons (p<0.0001**** and ns = not significant) **(S)** Structural snapshots of mC5a and mC5a^-d-Arg^ bound mC5aR1-Gαoβγ-ScFv16 complexes determined by cryo-EM at a global resolution of 3.15 Å and 3.13 Å, respectively (Gαo, yellow, Gβ, magenta Gγ, turquoise, ScFv16, gray). The right panel showing the interaction of carboxyl-terminal residues of mC5a and mC5a^-d-Arg^ with the residues present in orthosteric binding cavity of mC5aR1.

### Molecular mechanism of signaling-bias of C5a^-d-Arg^

In order to understand the molecular basis of signaling-bias exhibited by C5a^-d-Arg^, we determined the cryo-EM structure of mC5a-mC5aR1-G-protein and mC5a^-d-Arg^-mC5aR1-G-protein complexes (overall snapshot presented here in **Figure 1S**)^42^. Similar to the hC5a^-d-Arg^-hC5aR1-G-protein complex reported earlier^11^, we observed that the last three amino acids in mC5a^-d-Arg^ namely Q71, L72, and G73 slide into a binding pocket that is occupied by the terminal arginine in mC5a to compensate for critical interactions in the pocket (**Figure 1S**). Therefore, it is likely that the differences observed at the functional level are encoded by differential dynamics of these interactions. To probe this hypothesis, we employed molecular dynamics (MD) simulations^47,48^ using the structural templates of C5a and C5a^-d-Arg^ complexes.

We observed that R74 in C5a forms extensive polar interactions in the orthosteric binding site with Arg175^4.64^, Arg206^5.42^, and Asp282^7.35^ while G73 in C5a^-d-Arg^ can only partly recapitulate these interactions, for example, with Arg175^4.64^ and Arg206^5.42^, but not with Asp282^7.35^ (**Figure 2A and S2A**). (To avoid any ambiguity, one-letter amino acid code for ligand and three-letter amino acid code for receptor residues have been used). Interestingly, the loss of C5a^-d-Arg^ interaction with Asp282^7.35^ substantially alters the contact network in the orthosteric binding pocket of the receptor. In particular, we observed that Arg175^4.64^ and Arg206^5.42^ maintained some interaction with C5a^-d-Arg^ during the course of simulation, while Asp282^7.35^ did not engage with the ligand **(Figure 2A**). In contrast, all three residues maintained a stable interaction with C5a. Consistent with the MD simulation, site-directed mutagenesis of Arg175^4.64^ and Arg206^5.42^ to alanine resulted in a significant loss of G-protein-coupling for C5a^-d-Arg^ while the effects were rather modest for C5a **(Figure 2B, S2B and S2F**). Notably, all three mutations dramatically attenuated βarr recruitment for both C5a and C5a^-d-Arg^ (**Figure 2C, S2C and S2F**). These observations suggest that a loss of either one of these residues can be tolerated for C5a-induced G-protein activation due to sustained interaction with the remaining two residues, while the loss of even one of these interactions has a more drastic effect on C5a^-d-Arg^-induced G-protein-coupling. Conversely, engaging all three residues is critical for βarr recruitment by either ligand.

**Figure 2:**
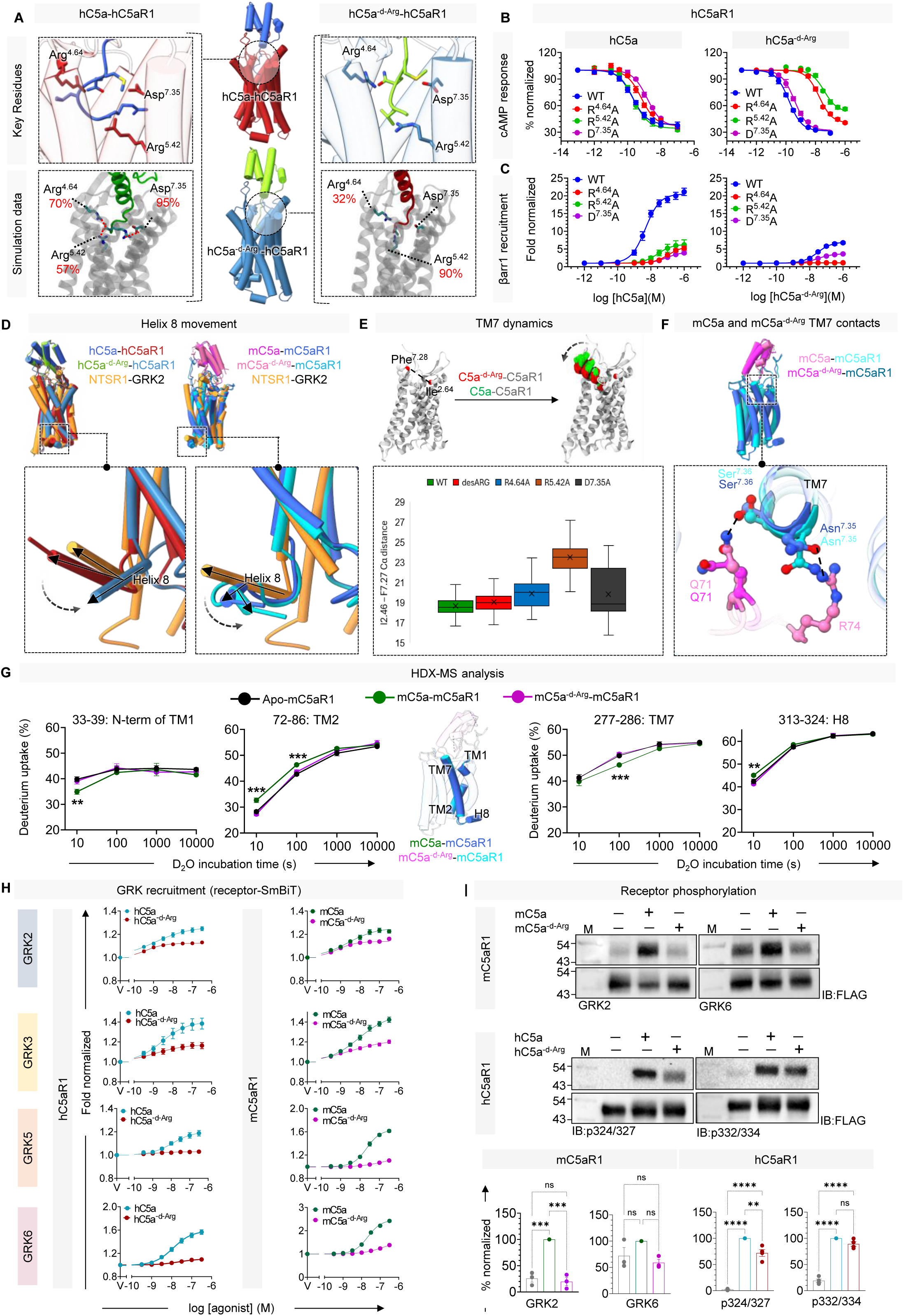
Mechanistic insights into G-protein bias driven by C5a^-d-Arg^. **(A)** Structural snapshots illustrating key interactions of hC5a-hC5aR1 (PDB: 8IA2) and hC5a^-d-Arg^-hC5aR1 (PDB: 8JZZ). Models are represented in tube helices with hC5a in royal blue, hC5a^-d-Arg^ in lime green, hC5a-bound hC5aR1 in brick-red and hC5a^-d-Arg^-bound hC5aR1 in dodger-blue. Molecular dynamic simulation (MDS) (lower panel) highlighting the percentage involvement of orthosteric binding residues with hC5a and hC5a^-d-Arg^. **(B-C)** cAMP inhibition (**B**) and βarr1 recruitment (**C**) measured downstream of hC5aR1 mutants (R^4.64^A, R^5.42^A, D^7.35^A), rationally designed based on MDS studies. Data represents mean ± SEM (n=3), independent experiments. **(D)** Cartoon representation comparing helix 8 movement in hC5a-hC5aR1 (8IA2) Vs. hC5a^-d-Arg^-hC5aR1 (8JZZ) and mC5a-mC5aR1 Vs. mC5a^-d-Arg^-mC5aR1 structures with respect to NTSR1-GRK2 (PDB: 8JPF) structure, revealing distinct positioning of helix 8 in C5a^-d-Arg^-bound C5aR1, potentially occluding GRK docking site in the receptor. **(E)** Structure-based simulation studies comparing C5a and C5a^-d-Arg^ activation of C5aR1 highlighting a key difference in TM7 conformation, measured by the distance between Ile^2.64^ and Phe^7.28^ (indicated by black dots) in C5a^-d-Arg^ and C5a-activated mutants. **(F)** Structural superposition of mC5a and mC5a^-d-Arg^-bound mC5aR1 highlighting the loss of hydrogen bonds with Ser^7.36^ and Asn^7.32^ of TM7 when bound to C5a^-d-Arg^. **(G)** Structural superposition of mC5a-mC5aR1 and mC5a^-d-Arg^-mC5aR1 (left-panel) highlighting the transmembrane regions (TM) undergoing reduced HDX. HDX-MS analysis (right) of mC5aR1 highlighted regions (left) undergoing significant decrease in deuterium uptake upon incubation either with ligand (mC5a/mC5a^-d-Arg^) or without ligand (Apo-mC5aR1). Statistical analysis was performed by applying one-way ANOVA followed by Tukey’s multiple comparisons test. (p<0.001***, p<0.01**). **(H)** NanoBiT-assay showing GRK recruitment downstream to (h/m) C5aR1 in response to (h/m) C5a and C5a^-d-Arg^.Data represents mean ± SEM (n=3), independent experiments, normalized with lowest ligand concentration considered as 1. **(I)** Phosphorylation detection via pIMAGO kit (upper panel) and phosphorylation site-specific antibodies (lower panel) to assess the receptor phosphorylation following ligand stimulation. Representative blots and densitometric analysis are shown below (green-mC5a, pink – mC5a^-d-Arg^, blue – hC5a and maroon – hC5a^-d-Arg^. Densitometric plots represent mean ± SEM (n = 3-4), independent experiments, with stimulated condition normalized with respect to unstimulated condition which is considered as 100% (p<0.0001****, p=0.0001***, p=0.0013**).

Structural comparison of C5a-C5aR1 and C5a^-d-Arg^-C5aR1 revealed an overall similar structure with comparable spatial positioning of the C5a^-d-Arg^ core domains on the receptor with a small linear shift, and the carboxy terminus residues of C5a^-d-Arg^ adopting a similar hook-like conformation as observed in C5a-C5aR1 (**Figure S2D-E**). Interestingly, helix 8 of C5aR1 in the C5a^-d-Arg^-bound structure undergoes a rotation of ∼120° and a linear shift of ∼5 Å (as measured from the Cα of Ser314^7.36^) towards the cytoplasmic portion of TM1 compared to that in C5a-C5aR1 structure (**Figure 2D**). Our MD simulation study also suggests that disrupting the contact pattern in the orthosteric binding pocket impacts the conformation space explored by TM7, wherein the loss of ligand interaction with TM7 (Asp282^7.35^) leads to a significant shift of TM7 (**Figure 2E**). These differences between the C5a vs. C5a^-d-Arg^-bound structures can be quantified in terms of the distance between TM2 and TM7 (Ile96^2.64^ and Phe275^7.28^), which is markedly larger in the C5a^-d-Arg^-bound C5aR1 (**Figure 2E**). In our mC5a/mC5a^-d-Arg^ structures, we also observe Q71 of mC5a establishes hydrogen bonds with Ser284 (S^7.36^) of mC5aR1, which is lost in mC5a^-d-Arg^-bound mC5aR1 (**Figure 2F**). Interestingly, HDX-MS experiments also show significant decrease in deuterium exchange in the N-terminus of TM1 and TM7, in mC5a-bound mC5aR1, compared to mC5a^-d-Arg^-mC5aR1, with the decrease being more profound in TM7. This observation aligns with the structural interpretation that TM7 is stabilized by the interactions established by mC5a with the receptor, and the loss of this interaction results in flexibility in TM7 of mC5a^-d-Arg^-bound mC5aR1 (**Figure 2G and S2G**). It is likely that this conformational change in TM7 propagates down to the intracellular side of the receptor and impacts the arrangement of helix 8.

A previously reported structure of NTSR1 in complex with GRK2 demonstrates a direct engagement of helix 8 in the receptor with the N-terminus of GRK2, thereby holding GRK2 in a spatial position to facilitate efficient receptor phosphorylation^49^. Using this as a reference, it is plausible that the spatial rearrangement of helix 8 in C5a^-d-Arg^-bound C5aR1 may impact efficient GRK engagement with the receptor. Indeed, a direct measurement of C5aR1 interaction with GRKs using a NanoBiT assay revealed a significant attenuation of receptor-GRK engagement upon stimulation with C5a^-d-Arg^ compared to C5a (**Figure 2H**). This further translates into inefficient phosphorylation of the C5aR1 as reflected in bulk phosphorylation measured using the pIMAGO assay, and site-specific phosphorylation measured using phospho-site-specific antibodies (**Figure 2I**). Taken together, these data provide a molecular explanation for attenuated βarr recruitment as observed in the cellular context upon stimulation of the receptor with C5a^-d-Arg^ compared to C5a, and ensuing signaling-bias.

### Structural basis of C5a-binding to C5aR2

Next, we focused our attention on C5aR2 in order to understand the molecular basis of naturally-encoded βarr-bias of this receptor despite binding the same natural agonists as C5aR1, i.e., C5a and C5a^-d-Arg^ (**Figure 3A**). Interestingly, we observed that unlike C5aR1, both C5a and C5a^-d-Arg^ display nearly identical potency and efficacy at C5aR2 in the βarr recruitment assays (**Figure 3B and S3A**). These observations in transfected cells align well with the *in vivo* data where the residual neutrophil mobilization upon C5a and C5a^-d-Arg^ stimulation with C5aR1 blockade, likely mediated by C5aR2, are nearly identical (**Figure 1R**). The structural analysis of C5aR2 using cryo-EM poses a challenge due to lack of G-protein-coupling, and previous efforts to isolate C5aR2-βarr complexes stabilized by Fab30 yielded only miniscule amounts of ternary complexes suitable for high-resolution analysis^12^. While purifying recombinant C5a and C5a^-d-Arg^, we use an N-terminal fusion of Thioredoxin (TrxA), which is a small (16 kDa), soluble, and thermostable protein^50,51^, and we reasoned that a complex of Trx-C5a/C5a^-d-Arg^-C5aR2 may help structural analysis using cryo-EM. Therefore, we first measured the pharmacology of Trx-C5a/C5a^-d-Arg^ vis-à-vis C5a/C5a^-d-Arg^, and observed that they were equally potent and efficacious in βarr recruitment assay (**Figure S3B**). Next, we reconstituted Trx-C5a-C5aR2 and Trx-C5a^-d-Arg^-C5aR2 complexes and subjected them to cryo-EM analysis. While both complexes exhibited 2D class averages with clear density for the ligand and the receptor, Trx-C5a^-d-Arg^-C5aR2 complex (**Figure S3C)** yielded a low-resolution 3D reconstruction while C5a-C5aR2 complex yielded a structure at an overall resolution of 3.8 Å (**Figure 3C and S4A**).

**Figure 3:**
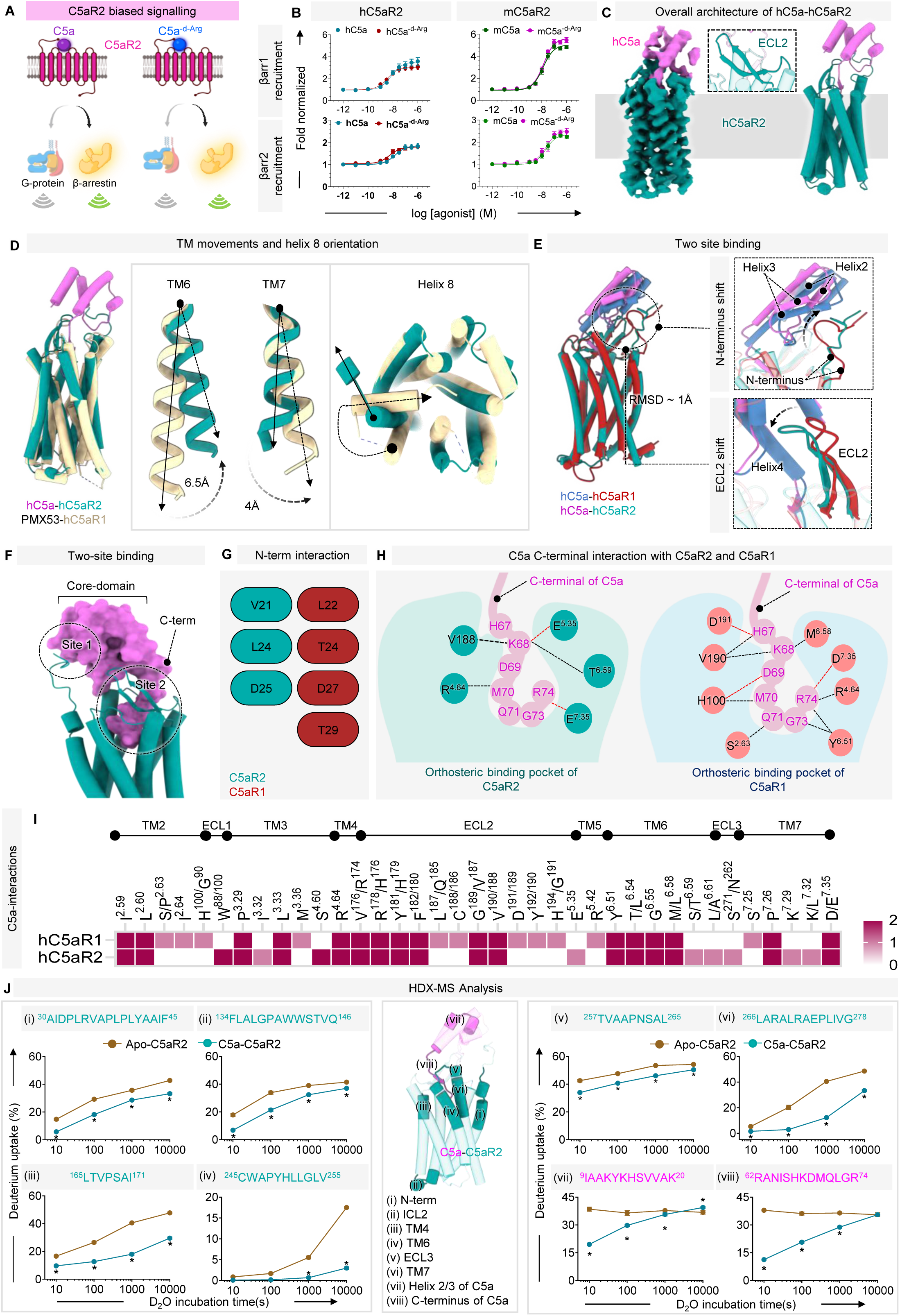
Functional and structural insights into C5a binding to C5aR2. **(A)** C5aR2 selectively signals through β-arrestin upon ligand binding, remains inactive in terms of G-protein signaling. Schematic was prepared in <u>BioRender</u>. **(B)** âarr1/2 recruitment downstream to (h/m) C5aR2 in response to (h/m) C5a and C5a^-d-Arg^ measured by NanoBiT-assay in HEK-293 cells. The data represents mean ± SEM (n=6), independent experiments where fold normalization is carried out by normalizing the luminescence values for each ligand dose with respect to the smallest ligand dose taken as 1. **(C)** Cryo-EM density map of Trx-C5a-bound C5aR2 at 3.8 Å and structural snapshot representing overall seven-transmembrane architecture of C5a-bound C5aR2 (show in tube helices) with ECL2 shown in ribbon representation. **(D)** Superposition of PMX53-bound C5aR1 (PDB:6C1R) and C5a-bound C5aR2, comparing transmembrane (TM6 and TM7) and helix 8 orientation in C5aR2 upon activation. **(E)** Superposition of C5a-bound C5aR1 (PDB:8IA2) and C5a-bound C5aR2, comparing differential interactions of C5a with C5aR1 and C5aR2. Upper-right panel showing an upward shift in the N-terminus of C5aR2 to establish contact with helix 2 of C5a. Lower-right panel depicting shift in ECL2 of C5aR2 towards cytoplasmic cavity with respect to C5aR1 ECL2. **(F)** Two-site binding of C5a (shown in surface) and C5aR2 (shown in tube helices). **(G-H)** Schematic representation of C5a core helices interaction with N-terminus **(G)** and carboxyl-terminal terminal interaction with C5aR2 and C5aR1 orthosteric pocket residues **(H)**. Black and red dotted lines represent hydrogen bonds and salt-bridge interactions, respectively. **(I)** Heat map displaying global interaction map of C5a-bound to C5aR1 and C5aR2. Each column represents an analogous residue position in C5aR1/2, where an interaction with C5a may be mapped. TM-helix residues have been numbered according to the Ballesteros-Weinstein (BW) numbering scheme and those in the N-term or loops have been numbered according to their order of occurrence in the receptor. To determine conserved residue positions, transmembrane helix residues were looked up from the GPCRdb (GPCRdb.org). If the residues at a particular BW-position are different across the receptors, they have been indicated. For example, at the BW 2.63 position, because C5aR1 has S and C5aR2 has P, it has been indicated as S/P^2.63^. For loops, where the GPCRdb did not report corresponding residues, they were inferred from the structural superposition of hC5a-bound hC5aR1 (PDB 8IA2) and hC5a-bound C5aR2. Similar to TM-residue numbering, any difference in the residue or its number at a corresponding position have been indicated. Heat-map scale shows whether a particular residue of C5aR1 or C5aR2 is involved in the interaction with C5a. If a particular residue from both the receptors is involved in interaction with C5a, it is given a score of 2, if it exists in only one receptor, it has given a score of 1 and the corresponding interaction for the other receptor is given a score of 0. Scores are encoded in colors and visualized as heat-maps. **(J)** Cartoon representation of C5a-bound C5aR2 highlighting the regions undergoing significant reduction in deuterium uptake. (i-viii) Deuterium uptake plots of selected peptides of C5aR2 and C5a (brown: Apo-C5aR2, turquoise: C5a-C5aR2). Results were derived from three independent experiments. The statistical significance of the differences was determined using Student’s t-test (*p< 0.05). Data are presented as mean ± standard error of the mean. * Indicates statistically significant difference between alone and complex, respectively.

The C5a-C5aR2 structure exhibits the canonical 7TM architecture with the 2^nd^ extracellular loop (ECL2) adopting an anti-parallel β-hairpin conformation as previously observed in the structures of C5aR1^11^ (**Figure 3C**). Despite moderate resolution, the cryo-EM map allowed unambiguous modelling of the transmembrane region, ECLs and ICLs, and C5a (**Figure S6A**). In addition, clear density for the distal N-terminus of the receptor from Pro20 and helix 8 are also observed in the structure (**Figure 3C and S8**). Although a structure of C5aR2 in a prototypical inactive state is not available, the comparison with antagonist-bound C5aR1 structure published previously^52^ (PDB: 6C1R), reveals significant conformational changes reminiscent of an active receptor conformation. For example, TM6 of C5aR2 shifts outward by ∼6.5 Å (as determined with respect to the Cα positions of Leu327^6.34^ in C5aR1 and Arg232^6.34^ in C5aR2), while TM7 moves inward by ∼4 Å (measured from the Cα atoms of Gly304^7.57^ in C5aR1 and Gly295^7.57^ in C5aR2) (**Figure 3D**). Moreover, the comparison of C5a-C5aR2 structure with that of C5a-C5aR1 shows an overall similar conformation with a main-chain RMSD of ∼1 Å (**Figure 3E**). In the inactive C5aR1 structure, helix 8 assumes an inverted orientation, and reorients itself significantly upon receptor activation^11,52^. The position of helix 8 in C5a-C5aR2 structure is also reminiscent of the active state C5aR1 (**Figure 3D**).

The core domain of C5a interacts with the N-terminus of C5aR2 while the carboxyl-terminus engages with the orthosteric binding pocket representing a two-site binding mode, similar to that of C5aR1 (**Figure 3F**). The extensive interaction between C5aR2 N-terminus and C5a core domain, a defining characteristic of the C5a-C5aR2 structure, involves several non-bonded contacts including the interaction of K20, D24, and I41 of C5a with Val21, Leu24, and Asp25 of C5aR2, respectively (**Figure S3D**). Interestingly however, there are also notable differences between C5a-binding to C5aR1 and C5aR2 (**Figure 3G-I and S3D-F**). For example, the N-terminus of C5aR1 is oriented to engage with the loop between helix 2 and helix 3 (H2-H3 loop) of C5a, while the N-terminus of C5aR2 is shifted such that it interacts with H2 as well as H3-H4 loop of C5a (**Figure 3E and S3D-E**). While ECL2 of C5aR1 interacts with the H1 residues of C5a, in C5aR2, ECL2 is displaced towards the extracellular opening of the orthosteric pocket and positions near to the H2-H3 loop in the C5a-C5aR2 structure (**Figure 3E and S3F**). These observations are further corroborated by site-directed mutagenesis data on C5aR1 as discussed previously^42^, and C5aR2 here (**Figure S3G-H**). For example, Arg^5.42^ is conserved in both C5aR1 and C5aR2 but makes contact with C5a only in C5aR1. Accordingly, its mutation to alanine leads to a dramatic loss of βarr1 recruitment for C5aR1 (**Figure 2C**) while it remains unchanged for C5aR2 (**Figure S3G**). Interestingly, the pattern of βarr recruitment upon C5a and C5a^-d-Arg^ stimulation of a set of C5aR2 mutants reflect a near-identical response, suggesting a conserved mode of binding of these two ligands to C5aR2 leading to a similar potency and efficacy as mentioned earlier (**Figure 3B and S3G**).

In order to validate the structural observations further, we performed HDX time-kinetics analyses on apo-C5aR2 and C5a-C5aR2 complexes. We observed a robust sequence coverage of C5a and C5aR2 in these experiments as outlined together with the technical details in **Figure S9A-B**. Upon C5a binding, the HDX levels of several regions in C5aR2 showed a decrease including the N-terminus, ECL3, TM4, TM6, TM7, and ICL2 (**Figure 3J, S9C and Supplementary Dataset 1B, 1C, 1E and 1F**). This pattern is in sync with the cryo-EM structure such as a direct interaction of C5a with the N-terminus and ECL2 region. In addition, the observed reduction in HDX levels in the TM regions suggest possible activation-dependent conformational changes through allosteric mechanism, which is also corroborated by the structural comparison of C5a-C5aR2 structure with the antagonist PMX53-bound inactive state of C5aR1^52^ (PDB: 6C1R). Furthermore, the decrease in HDX levels at the ICL2 and the distal end of TM7 also likely reflect agonist-induced conformational propagation resulting in a short helix formation in ICL2 and reorientation of helix 8 upon activation. Finally, the comparison of HDX level of C5a in the free and receptor-bound state reveals a significant decrease in both, the carboxyl-terminal and amino-terminal regions, which aligns well with the two-site binding mode of C5a on C5aR2 (**Figure 3J and Figure S9D)**.

### Structural basis of C5a^-d-Arg^-recognition by C5aR2

Our functional characterisation of mouse C5aR2 for βarr1/2 recruitment in HEK-293 cells shows near identical response for mC5a and mC5a^-d-Arg^ as observed for hC5a and hC5a^-d-Arg^ on hC5aR2 **(Figure 3B)**. We reasoned that mC5a^-d-Arg^-mC5aR2 may yield a complex better amenable to structural analysis compared to the human receptor, and thereby, help us in decipher the binding mode of C5a^-d-Arg^ on C5aR2. Interestingly, in our attempt to purify mC5aR2, we observed that unlike hC5aR2, it exhibits a distinct and substantial dimeric population (**Figure 4A**), and it allowed us to determine the structures of mC5a^-d-Arg^-bound mC5aR2 complex as a dimer using cryo-EM. As anticipated, structure determination revealed dimeric assembly of mC5aR2 wherein both the protomers lie adjacent to each other in a single detergent micelle bound to mC5a^-d-Arg^ (**Figure 4B, Figure S4B, S6B and S8**). The mC5a^-d-Arg^-mC5aR2 exhibits the canonical 7TM architecture with the 2^nd^ extracellular loop (ECL2) adopting an anti-parallel β-hairpin conformation as previously observed in the structures of C5aR1 (**Figure 4B**). The structural superposition of hC5aR2 and one of the protomer of mC5aR2 showed a comparable RMSD values for TM regions while TM1 and cytoplasmic loops and helix 8 displayed significant deviation (**Figure 4C**). Moreover, our structural analysis uncovered a previously unrecognised dimeric assembly in mouse C5aR2, which is stabilised by the two inter-protomer disulfide bonds between Cys306 and Cys313 in helix 8 of opposing protomers. This is further held together by tight hydrophobic packing between Phe^1.43^ and Leu^1.44^ of TM1 (**Figure 4D**). To our knowledge, this represents the first report of a dual disulfide-mediated covalent linkage stabilising a class A GPCR dimer. The presence of this covalent interface exclusively in mouse C5aR2 suggests a species-specific structural adaptation that may have significant functional consequences compared to human receptor. This finding broadens the current understanding of GPCR dimerization and also underscores the significance of species-specific receptor functions. Similar to C5a, C5a^-d-Arg^ also displays a two-site binding mode on C5aR2, acquires an overall similar positioning as C5a with slight shift in the core-domain (**Figure 4E**). This is also reflected in identical positioning of terminal residues i.e., R74 and G76 of C5a and C5a^-d-Arg^, respectively (**Figure 4F**). A comparative analysis of ligand-receptor interactions reveals structural features that may contribute to the retained potency of C5a^-d-Arg^ at C5aR2. The binding of C5a^-d-Arg^ to mC5aR2 is characterized by an extensive interaction, wherein G76 aligns in the position of R74 in C5a and forms hydrogen bonds with Arg179^4.64^ and the backbone nitrogen of Arg210^5.42^ (**Figure 4F and 4G**). The C-terminal region engages in multiple additional interactions, for example, K71 forms salt bridge with Glu^5.35^ and hydrogen bond Ile^6.58^, H70 forms salt-bridge with Gly193^ECL2^, Q74 and L75 forms hydrogen bonds with Glu^7.35^ and Arg^6.55^. This results in a network comprising ten hydrogen bonds and one salt-bridge distributes across C-terminal residues (**Figure 4G**). Taken together with the structural analysis of C5a^-d-Arg^-C5aR1, these observations suggest that while C5aR1 may rely more critically on R74-mediated contacts for conformational stabilisation linked to βarr signalling, C5aR2 can accommodate R74 loss by redistributing interactions to alternative residues in the C-terminal region, thereby maintaining receptor activation to an efficacy similar to that of C5a.

**Figure 4.**
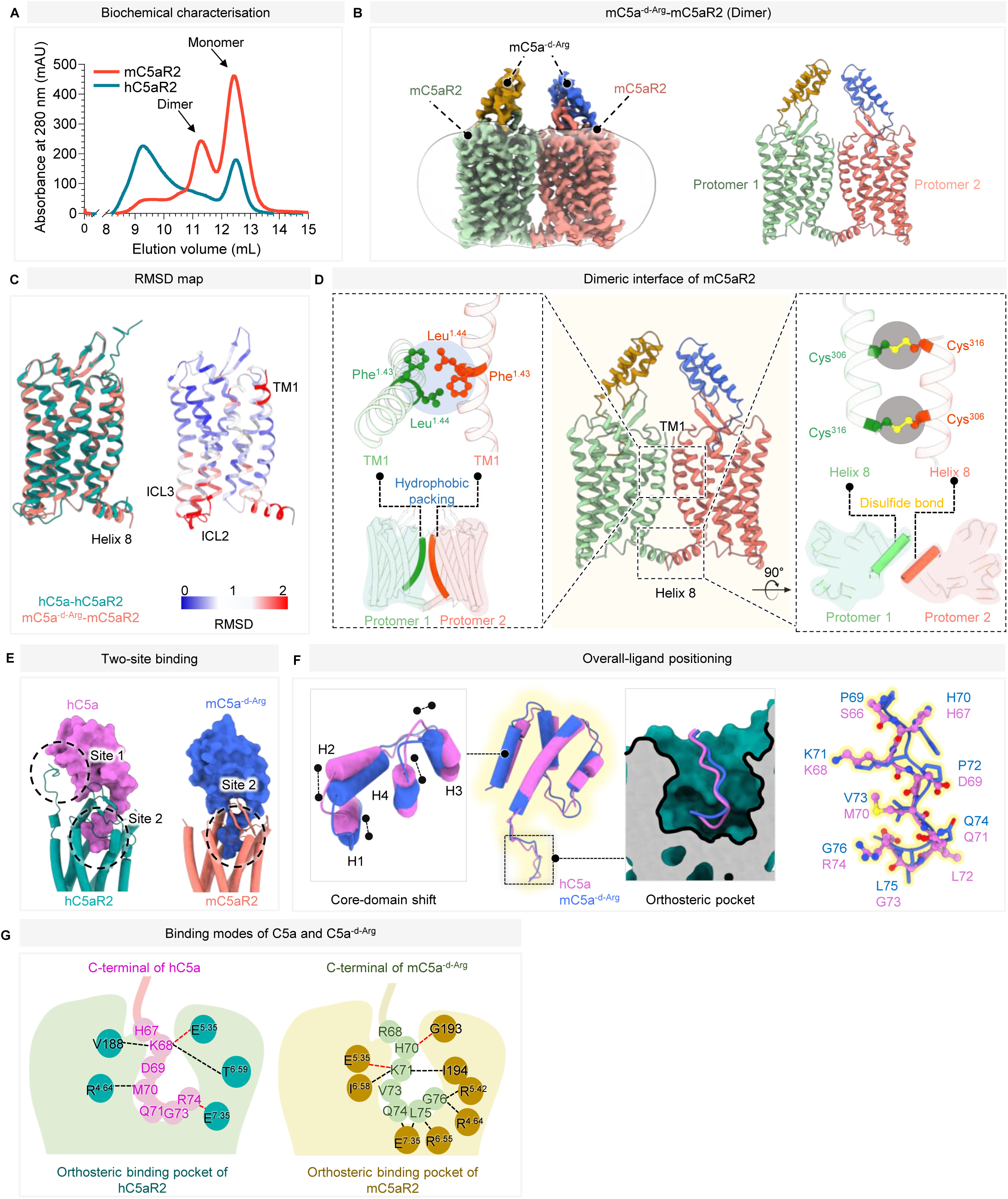
**Molecular insights into C5a and C5a^-d-Arg^ binding to C5aR2.** (A) Size-exclusion chromatography profile of the purified human and mouse C5aR2, revealing a substantial dimeric population for the mouse receptor. (B) Structural snapshots of cryo-EM densities and atomic models of mC5a^-d-Arg^-mC5aR2 complex. (C) Structural comparison of the human C5aR2 and mouse C5aR2 using superimposition and RMSD across the entire receptor mapped onto the mouse C5aR2. (D) Dimeric interface of the mC5aR2 indicating TM1-TM1 interface with the involvement of key residues, and H8-H8 interface mediated by two disulfide bridges between the indicated residues. (E) Structural snapshots revealing two-site binding mechanism of C5a-C5aR2 and mC5a^-d-Arg^-mC5aR2 (F) Superposition of hC5a and mC5a^-d-Arg^ obtained from the hC5a-hC5aR2 and mC5a^-d-Arg^-mC5aR2 structures, showing overall ligand positioning, core-domain shift, C-terminus positioning in C5aR2 orthosteric pocket and terminal G76 of mC5a^-d-Arg^ occupying similar positioning as that of R74 in hC5a. (G) Schematic representation of C-terminal residues of hC5a and mC5a^-d-Arg^ interacting with different residues lining orthosteric pocket of hC5aR2 and mC5aR2 highlighting extensive interaction network of C5a and C5a^-d-Arg^ (black dashed lines indicate hydrogen bonds, red-dashed lines indicate salt-bridges.

### Discovery of a C5aR2-selective agonist

Considering the similarities between the interaction of C5a with C5aR1 and C5aR2 in the orthosteric pocket, we envisioned that peptides derived from the carboxyl-terminus of C5a may also serve as C5aR2 agonists (**Figure 5A**). Accordingly, we screened a broad set of C5a- and C3a-derived peptides on C5aR2 in βarr recruitment assay (**Figure 5B and S10A**) and observed that C5a^pep^ and EP54 exhibited a significant response (**Figure 5B and S10E-F**). However, these peptides are also known to activate C3aR and C5aR1^11^, and previously described C5aR2-selective peptides, i.e., P32 and P59, have rather low efficacy (**Figure 5B**). Therefore, we revisited our previous data focused on the identification of selective peptide agonists for C5aR1^53^, and we identified two peptides namely, BM2020-7 and BM2020-8 that had improved efficacy and potency at C5aR2 compared to P32 and P59^54^ (**Table S1**). However, neither peptide was selective for C5aR2 with BM2020-7 having moderate activity at C3aR and BM2020-8 being a potent agonist of both C3aR and C5aR1. Considering that C5a and C5a^-d-Arg^ have similar efficacy at C5aR2, we hypothesised that we could modify the C-terminal arginine of BM2020-7 and BM2020-8 to introduce selectivity for C5aR2. This hypothesis is also substantiated by our previous observation that the terminal arginine in these peptides is crucial for their potency at C3aR and C5aR1^11^.

**Figure 5:**
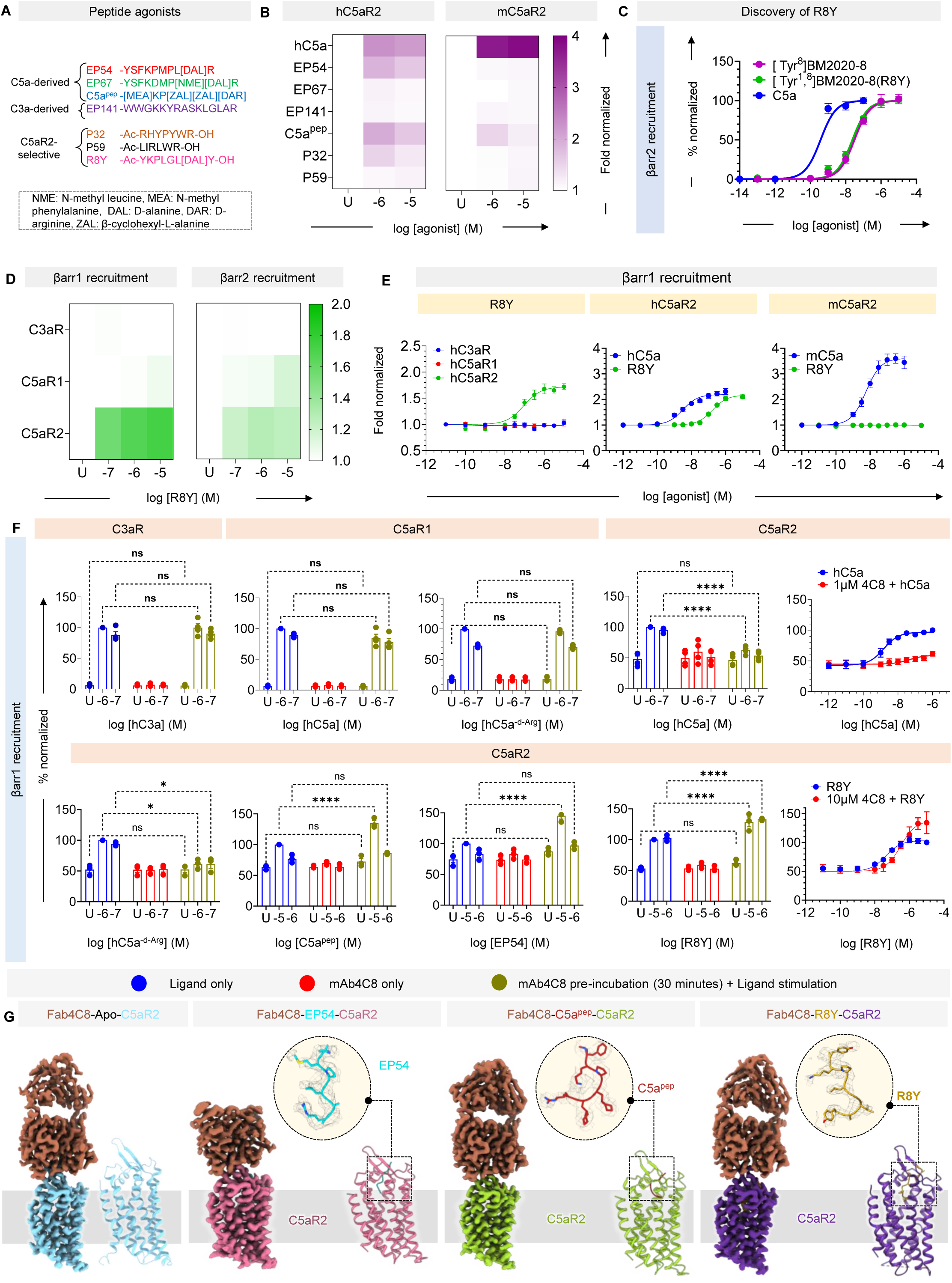
Comprehensive characterization of C5aR2-specific peptide ligands and previously discovered C5aR2-blocking antibody mAb4C8. **(A)** Sequence of small peptide ligands known to activate C3aR and C5aR1 along with C5aR2-selective peptides, P32 and P59 and novel decapeptide, R8Y. **(B)** Screening of previously published peptides to assess agonistic activity on human and mouse C5aR2 measured by βarr1 recruitment using NanoBiT-based assay. The data represents mean values for two independent experiments. The heat map shows fold normalized values for two different ligand doses. The raw counts for ligand stimulated conditions were normalized with respect to unstimulated conditions taken as 1. **(C)** âarr2 recruitment downstream to C5aR2 in response to the novel peptide, R8Y, rationally deigned by modifying BM2020-8 peptide backbone. For measuring βarr2 recruitment, BRET-based luminescence assay was employed and % normalization was carried out by normalizing response for varying ligand doses with respect to that of the highest ligand dose taken as 100. **(D)** âarr1/2 recruitment to investigate selectivity profile of R8Y in activating C3aR, C5aR1 and C5aR2 at two ligand doses. NanoBiT-based assay was employed in HEK-293 cells transiently transfected with either C3aR or C5aR1 or C5aR2. Heat map showing mean values for three independent experiments, normalized with respect to unstimulated condition considered as 1. **(E)** Dose-response curve showing R8Y-induced βarr1 recruitment downstream to hC3aR/hC5aR1/hC5aR2/mC5aR2 highlighting its species-specific and C5aR2-selective nature. **(F)** Characterization of mAb4C8 by measuring its agonistic and antagonistic properties on C3aR, C5aR1 and C5aR2. NanoBiT-based βarr1 recruitment was performed under three conditions, viz. native ligand stimulated (in blue), mAb4C8 stimulated (to assess agonistic properties; in red) and pre-incubation of mAb4C8 for 30 minutes prior to induction with ligand doses (to assess antagonistic properties; in olive-green). The assay displays antagonistic activity of mAb4C8, selectively for C5aR2, marked by severe reduction in C5a and C5a^-d-Arg^-mediated βarr1 recruitment in the presence of mAb4C8. The right-most panel displays hC5a- or R8Y-induced dose-response curve for βarr1 recruitment downstream to hC5aR2, in the presence or absence of mAb4C8, at indicated molar concentrations. Data represents mean ± SEM for three-four independent experiments. Statistical significance of the data was carried out by applying two-way ANOVA followed by Tukey’s multiple comparisons test (p<0.0001****). **(G)** Structural snapshots of cryo-EM density map and models of C5aR2 bound to EP54, C5a^pep^ and R8Y along with Fab4C8 or without any ligand, i.e., Apo-C5aR2. To left, EM-density map and to right, respective model map presented in cartoon style.

Therefore, we synthesised a set of BM2020-7 analogues where we systematically modified the C-terminal residue to asparagine, aspartic acid, glutamine, glutamic acid, histidine, leucine, lysine, phenylalanine, tryptophan, tyrosine, or serine. We then assessed the ability of these peptides to activate C5aR2-mediated βarr2 recruitment at a concentration of 100 µM. Of these peptides, the tyrosine analogue demonstrated full agonist activity (relative to C5a) at C5aR2 (**Figure S10B)** and subsequent concentration-response experiment revealed that it had an EC50 of ∼0.9 μM (**Figure S10C**). As predicted, this tyrosine analogue did not activate C3aR or C5aR1-mediated ERK phosphorylation up to 100 µM (**Table S1**). As BM2020-8 was a more potent, albeit non-selective, agonist of C5aR2 compared to BM2020-7, we replaced the arginine in BM2020-8 with tyrosine to produce ^[Tyr8]^BM2020-8, which had an EC50 of 35 nM at C5aR2 (**Figure 5C**) but was a partial agonist at C5aR1 (EC50 = ∼120 nM, 41% efficacy relative to C5a). Finally, we replaced the phenylalanine at position 1 in ^[Tyr8]^BM2020-8 with tyrosine to generate ^[Tyr1, Tyr8]^BM2020-8, referred to as R8Y hereon, which remained a full agonist of C5aR2 (EC50 = ∼13 nM) (**Figure 5C**), and it did not activate C3aR or C5aR1 up to 10 µM (**Figure S10D and Table S1**). Finally, we confirmed the subtype selectivity of R8Y in HEK-293 cells expressing comparable levels of C3aR, C5aR1, and C5aR2 (**Figure 5D and S10G**).

### Molecular basis of C5aR2 activation by peptide agonists

In order to understand the molecular basis of subtype-selectivity and activation of C5aR2 by the peptide agonists, we focused our efforts on determining the structures of C5aR2 in complex with EP54 (C3aR/C5aR1/C5aR2 cross-reactive), C5a^pep^ (C5aR1/C5aR2 cross-reactive), and R8Y (C5aR2 selective). As the strategy used for C5a-C5aR2 is not feasible here, we tested a previously described anti-C5aR2 monoclonal antibody, referred to as 4C8^55^, as a fiducial marker for cryo-EM analysis (**Figure S11A**). We first confirmed the ability of this antibody to recognize C5aR2 and observed that it effectively blocks C5a-induced βarr recruitment at C5aR2 but not at C5aR1 or C3a-induced βarr recruitment at C3aR (**Figure 5F and S10L**). We also did not observe any agonistic effect of 4C8 by itself, and therefore, taken together, these data confirm that 4C8 acts as a competitive and selective inhibitor of C5a at C5aR2. Interestingly however, 4C8 pre-incubation did not impact βarr recruitment at C5aR2 in response to any of the peptide agonists suggesting that binding of 4C8 is permissive for peptide interaction and presumably receptor activation (**Figure 5F and S10L**).

Next, we generated the Fab version of 4C8 using papain digestion and subsequent purification, and observed that it formed a stable complex with C5aR2 either in the apo-state or in presence of peptide agonists (**Figure S11B-E**). We successfully determined the cryo-EM structures of Fab4C8-stabilized apo-C5aR2, EP54-hC5aR2, C5a^pep^-C5aR2, and R8Y-C5aR2 complexes at 3.0 Å -3.2 Å resolution range (**Figure S4C and S5A-C**), and the cryo-EM maps allowed unambiguous modelling of the receptor regions and the ligands (**Figure 5G and S7A-D**). Structural alignment of these structures reveals an overall similar conformation, with an RMSD of <1 Å across the main-chain atoms (**Figure 6A**). A complete description of the residues that are resolved is presented in **Figure S8**. Expectedly, unlike the C5a-C5aR2 structure, the N-terminus of the receptor was not resolved well in these structures owing to a lack of interaction with the peptide agonists, which are likely engaged only with the orthosteric binding pocket. In addition, the carboxyl-terminus of the receptor including helix 8, and some of the ICL2 and ICL3 loops were also not resolved well, indicating their structural flexibility (**Figure S8**).

**Figure 6:**
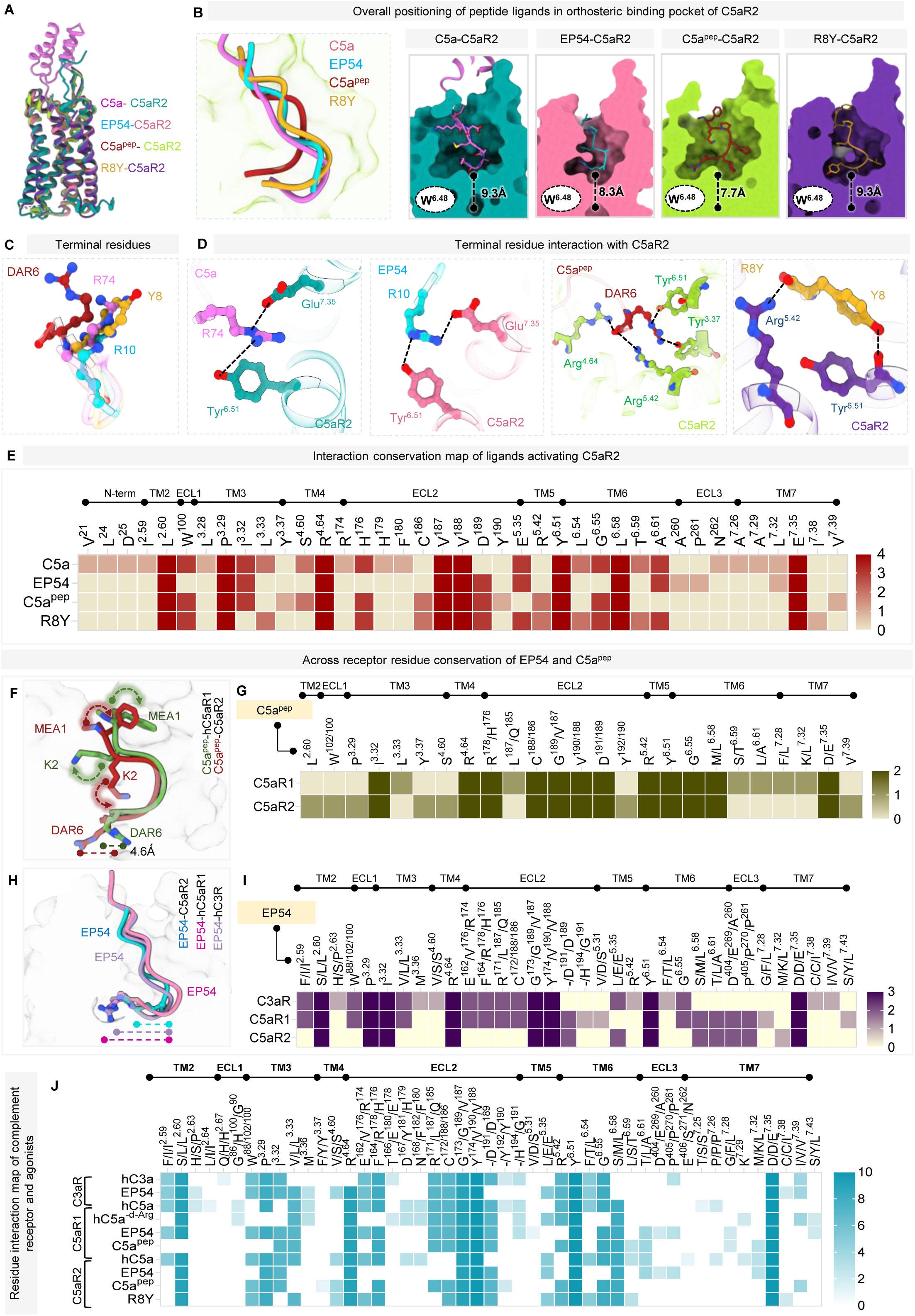
Molecular insights into diverse ligand recognition by C5aR2. **(A)** Overall structural superposition of C5aR2 bound to C5a, EP54, C5a^pep^, and R8Y. **(B)** Superposition of the C-terminal part of C5aR2-bound ligands, in the C5aR2 orthosteric pocket in the C5a^pep^-C5aR2 structure (green background). The hook-shaped C-termini of all ligands (shown in ribbon and atom representation) fit into a similar binding cavity in the orthosteric pocket of C5aR2, shown as surface slices, with the depth of penetration of each ligand from the conserved W^6.48^ residue of C5aR2. **(C-D)** Superposition of the C-terminal residues (R74 in C5a, R10 in EP54, DAR6 in C5a^pep^ and Y8 in R8Y) of C5aR2-bound ligands showing their similar or distinct orientations in the C5aR2 orthosteric pocket **(C)**, which leads to them engaging distinct sets of residues for interaction **(D)**. **(E)** Heatmap summarizing interactions between the ligands C5a, R8Y, EP54 and C5a^pep^ with C5aR2, as inferred from the respective structural snapshots, where each row represents a ligand, and each column represents a C5aR2 residue where an interaction with a ligand may be mapped. TM-helix residues have been numbered according to the Ballesteros-Weinstein (BW) numbering scheme and those in the N-term or loops have been numbered according to their order of occurrence in the receptor. **(F)** Superposition of C5a^pep^ bound to C5aR1 and C5aR2 depicting slight differences in the orientation or positioning of some residues (shown by dashed straight or curved lines), although the overall conformation of the ligand backbone is maintained. **(G)** Heatmap comparing interactions of C5a^pep^ with C5aR1 (PDB: 9UMR) or C5aR2. The residue numbering and analogous residue determination scheme is same as in Figure 3I. **(H)** Superposition of EP54 bound to either hC3aR, hC5aR1 or C5aR2, showing an overall conserved hook-like conformation, with slight shifts in the C-termini, indicated by dashed lines **(I)** Heatmap comparing EP54 interactions with analogous receptor residues across EP54-bound C3aR (PDB 8I95), EP54-C5aR1 (PDB: 9UMX) and EP54-C5aR2. The residue numbering and analogous residue determination scheme is same as in Figure 3I, with the inclusion of analogous positions for hC3aR as well, using the hC3a-hC3aR as a reference (PDB: 8I9L). **(J)** Heatmap comparing ligand-receptor interactions at analogous positions across complement receptors, viz., hC3a-hC3aR (PDB: 8I9L), EP54-hC3aR (PDB: 8I95), hC5a-hC5aR1 (PDB 8IA2), hC5a^-d-Arg^ –hC5aR1 (PDB: 8JZZ), EP54-hC5aR1 (PDB: 9UMX), C5a^pep^-C5aR1 (PDB: 9UMR), hC5a-C5aR2, EP54-C5aR2, C5a^pep^-C5aR2 and R8Y-C5aR2. The residue numbering scheme is same as in Figure 3I.

In addition to the N-terminus, the apo-C5aR2 structure exhibits notable flexibility in other regions as well when compared to the C5a-C5aR2 structure (**Figure S12A**). Specifically, the loop connecting β-strand 1 and β-strand 2 of ECL2 (G178–R183) shifts away from the orthosteric binding pocket, likely due to the absence of stabilizing interactions with the core domain of C5a (**Figure S12B**). Likewise, the extracellular end of TM6 and TM7, and the ECL3, exhibit an outward movement in the apo-C5aR2 structure compared to C5a-C5aR2 (**Figure S12A-B**). This structural rearrangement is supported by an increase in the HDX level in this region in apo- vs. C5a-bound states of C5aR2 (**Figure 3J**). The intracellular side of the receptor shows an overall smaller cavity in the apo-C5aR2 structure with an approximate volume of ∼3500 Å^3^ compared to that of C5a-C5aR2 with a volume of ∼4200 Å^3^. This difference likely originates from distinct positioning of the TM helices on the intracellular side in the apo-C5aR2, which emulates an inactive-like conformation of receptor with reference to the antagonist-bound C5aR1 structure^52^ (PDB: 6C1R) (**Figure S12B**).

Reminiscent of the binding mode of C5a, the peptide agonists EP54, C5a^pep^ and R8Y also adopt a hook-like conformation and penetrate deep into the orthosteric pocket of the receptor at a vertical distance of ∼8-9 Å from the conserved Trp246^6.48^, and make extensive interactions with the residues from TM2-7, ECL1 and ECL2 (**Figure 6B**). In accordance with the likely dispensable role of the terminal arginine (R74) of C5a in C5aR2 activation as reflected by C5a^-d-Arg^ pharmacology, the terminal residues of C5a^pep^ (i.e., d-Arginine) and R8Y (tyrosine) engage via either hydrogen bonds or polar networks with Glu273^7.35^, Tyr249^6.51^and Arg173^4.64^, similar to that observed for C5a and EP54 (**Figure 6C-D**). A comparison of overall interactions for the peptide agonists and C5a revealed a set of residues common to all the agonists, located either within the orthosteric pocket, or in the ECL2, suggesting a convergent mode of ligand recognition (**Figure 6E**). On the other hand, C5a engages with a few additional unique residues, particularly on ECL3 and the N-terminus, which likely imparts a higher potency for the receptors, analogous to that observed for C5aR1 as well.

Structural analysis of C5a^pep^-C5aR2 structure reveals that C5a^pep^ forms seven hydrogen bonds along with several non-bonded contacts within the orthosteric pocket of C5aR2. The terminal DAR6 in C5a^pep^ forms ionic bond and cation-π interaction with Tyr249^6.51^ and hydrogen bonds with Arg173^4.64^ and Arg204^5.42^, while it engages with Tyr119^3.37^ through ionic interaction (**Figure 6D**). In addition, other non-bonded contacts that maintain the conformation of DAR6 in C5a^pep^ include contacts with Ser169^4.60^ and Gly253^6.55^ of C5aR2 (**Figure S12C**). K2 in C5a^pep^ engages with Val188^ECL2^ and Glu273^7.35^ through a hydrogen bond and with Val187^ECL2^ through a non-bonded contact while MEA1 in C5a^pep^ makes a π-π interaction with His176^ECL2^ and engages with Val187^ECL2^ and Asp189^ECL2^ through non-bonded contacts to further facilitate the stabilization of the ligand within the orthosteric pocket. (**Figure S12C**).

Further, we compared the binding modes of C5a^pep^ on C5aR1 and C5aR2 using the structures presented here with those reported recently^42^. We observed that, although the overall backbone conformation of C5a^pep^ in the C5a^pep^-C5aR2 structure closely resembles with that observed in C5a^pep^-C5aR1 complex, there is a notable difference in the orientation of the terminal DAR6 residue between the two receptors (**Figure 6F**). In the C5a^pep^-C5aR2 structure, the guanidinium group of the DAR6 residue undergoes a linear horizontal shift of ∼5 Å towards TM5 within the orthosteric pocket compared to C5a^pep^-C5aR1. Furthermore, the orientation of the MEA1 and K6 in C5a^pep^ between the C5aR1 and C5aR2 are also distinct as they are positioned opposite to each other. Despite these differences, comparing the overall interaction of C5a^pep^ with C5aR1 and C5aR2 reveals a similar interaction pattern, as expected from its ability to activate both C5aR1 and C5aR2 (**Figure 6G)**.

Even though C5a^pep^, EP54 and R8Y exhibit a similar binding mode on C5aR2, there are clear differences while comparing the intricate details of the binding of these three peptide agonists at C5aR2, as well as when comparing EP54 binding to C5aR1 vs. C5aR2, and R8Y vs. C5a binding to C5aR2. For example, compared to the positioning of the guanidino group of the terminal arginine of EP54 in the EP54-hC5aR1 structure^42^, the terminal Arg10 of EP54 in the EP54-C5aR2 structure appears to be slightly constrained, and docks into an alternative sub-pocket within the orthosteric site (**Figure 6H**). Moreover, in C5aR2, EP54 is stabilized through the formation of hydrogen bonds with Glu273^7.35^ and Val188^ECL2^, and an ionic interaction with the hydroxyl group of Tyr249^6.51^(**Figure 6D**). Similar to C5a^pep^, several other residues of EP54 establish non-bonded contacts within the orthosteric binding pocket of C5aR2 (**Figure S12D**). Despite these subtle differences, a global interaction analysis of EP54 on C3aR, C5aR1 and C5aR2 revealed a conserved binding mechanism by the involvement of similar residues lining orthosteric binding pocket and ECL2, perhaps attributing to the agonistic property of EP54 across complement receptors (**Figure 6I**).

In the R8Y-C5aR2 structure, Y8 of R8Y undergoes a linear transition downwards by ∼5.5 Å relative to the spatial positioning of DAR6 in C5a^pep^-C5aR2, and it engages with Tyr249^6.51^ through a π-π interaction and with Arg204^5.42^ through a salt bridge (**Figure 6D**). Additionally, K2 of R8Y establishes hydrogen bonds with Val188^ECL2^ and Leu256^6.58^, while the backbone oxygen of G5 in R8Y forms hydrogen bonds with Arg173^4.64^ (**Figure S12E**). However, R74 of C5a in C5a-C5aR2 occupies the same position as Y8 of R8Y and establishes a hydrogen bond with Glu273^7.35^, while K68 of C5a forms three hydrogen bonds, one each with Val188^ECL2^, Glu197^5.35^ and Thr257^6.59^ (**Figure S12F**). These additional interactions by C5a compared to R8Y, together with the involvement of N-terminus of the receptor, possibly explains the relatively weaker potency of R8Y at C5aR2 compared to C5a. Finally, a structural alignment of ligand-receptor interactions for natural and synthetic-peptide agonist-bound structures of C3aR, C5aR1, and C5aR2 reveal that C5aR2 also employs a “Five-Point-Switch” to recognize diverse ligands (**Figure 6J**), as identified for C3aR and C5aR1 in the recent study^42^.

### Species-specific pharmacology and transducer-coupling of C5aR2

Inspired by the dramatic species-specific pharmacology observed at the C3aR and C5aR1^42^, we also probed the activity of the C5aR2-selective agonist discovered here i.e. R8Y. Strikingly, our functional assays revealed remarkable selectivity of R8Y for human C5aR2, and it does not seem to activate mouse C5aR2 as assessed using βarr recruitment assay (**Figure 5E and S10I and S10K**). In addition, the sequence comparison of the human and mouse C5aR2 shows a different phosphorylation signature in the carboxyl-terminus wherein the mouse receptor contains the P-X-P-P motif critical for driving βarr recruitment and activation while the human receptor does not (**Figure S13A**). This is further reflected in the pattern of reactivity of Ib30, an intrabody sensor designed to report an active conformation of βarr1 in cellular context^56^, wherein we observe a robust signal of Ib30 reactivity for the mouse receptor but not for the human C5aR2 (**Figure S13B-C**). These data further corroborate the species-specific specialization of transducer-coupling at C5aR2, which converges to a similar observation for C5aR1 in terms of βarr interaction and activation as reported recently^42^.

### Molecular insights into βarr-bias encoded at C5aR2

Comparison of the C5a-C5aR2 structure with C5a-C5aR1 complex (PDB: 8IA2) reveals a cytoplasmic pocket dimension of ∼30 Å (as measured from the Cα atoms of Trp141^ICL2^ to Phe291^7.53^) in C5aR2 with a pocket volume of 4,250 Å^3^ while in C5aR1, the pocket spans ∼22 Å (as measured from the Cα atoms of Cys144^ICL2^ to Tyr300^7.53^) with a pocket volume of ∼2,900 Å^3^. Thus, the cytoplasmic pocket of C5aR2 appears to be relatively wider, and in addition, it is also less charged and more hydrophobic compared to that of C5aR1. These differences may prevent a stable docking, and conformational changes required for an efficient coupling and activation of G-proteins to C5aR2 (**Figure 7A**). The polar and charged amino acids within the cytoplasmic pocket in the C5aR1 structure enable strong electrostatic and hydrogen-bond interactions with G-proteins, thereby stabilizing the complex and facilitating activation (**Figure 7A**). However, a significant variation in these key polar and charged residues can be observed in C5aR2, which instead has a more hydrophobic environment. These altered pocket properties are ill-suited to efficiently interact with the largely hydrophilic and charged surfaces of G-proteins, resulting in an absence of G-protein-coupling (**Figure S14A**).

**Figure 7:**
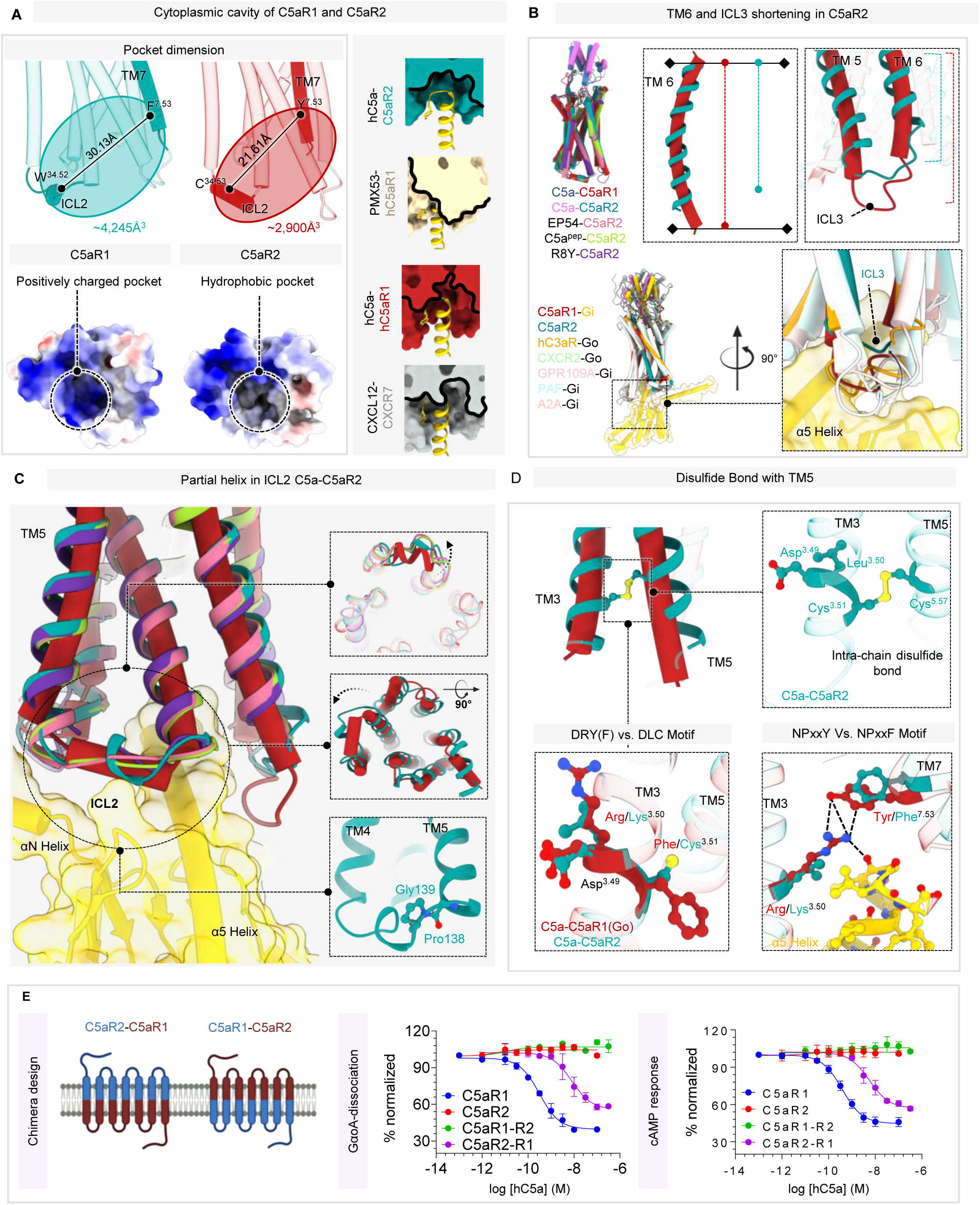
Structural insights into lack of G-protein coupling in C5aR2. **(A)** Surface electrostatic representation of C5aR1 and C5aR2 cytoplasmic cavity showing a positively-charged and hydrophobic cavity in C5aR1 and C5aR2, respectively. Right-most panel showing the comparison of cytoplasmic cavity of C5aR2 with respect to inactive-hC5aR1 (PMX53/Avacopan-hC5aR1; PDB: 6C1R), active-hC5aR1 (hC5a-hC5aR1-Go; PDB:8IA2), and the atypical chemokine receptor CXCR7 (CXCL12-CXCR7-FabCID24 ; PDB: 7SK5). The Gαo of hC5a-hC5aR1-Go (PDB 8IA2) structure has been docked into the cytoplasmic cavity of each receptor. The receptors are shown as surface slices and the α5 helix in ribbon representation. **(B)** Structural superposition of hC5a-bound hC5aR1 (PDB 8IA2) with that of C5aR2 bound to hC5a, EP54, C5a^pep^ and R8Y. Structural snapshots indicating TM5, TM6 and ICL3 shortening in C5aR2 (indicated by dashed arrow), to right. Lower-left panel indicates comparison of GPCR-Gα_o/i_ structures, viz., hC5a-hC5aR1-Go (PDB: 8IA2), hC5a-C5aR2, CXCL8-CXCR2-Go (PDB: 8XX6), Niacin-GPR109A-Gi (PDB: 8IY9), PAF-PAFR-Gi (PDB: 8XYD) and Epinephrine-α2AAR-Gi (PDB: 9CBL) superposed on the Gα_o_ of hC5a-C5aR1-Go (PDB 8IA2) structure showing ICL3 apposed against Gα making extensive contacts. Zoomed view (lower-right panel) of the loop and cavity shows the short ICL3 of C5aR2 is unable to dock in this cavity. **(C)** Conformational dynamics of ICL2 in C5aR2. Structural superposition of hC5a-bound hC5aR1 (PDB 8IA2) with that of C5aR2 bound to hC5a, EP54, C5a^pep^ and R8Y, docked on the Go of hC5a-hC5aR1-Go (PDB 8IA2) indicates that the ICL2 helix is either not formed (in EP54, C5a^pep^ or R8Y-bound C5aR2) or partially formed (in hC5a-C5aR2), in contrast to hC5aR1, where it is completely formed. This leads to loss of interactions it makes in a cavity formed by the α5 and αN helices of Gα. Inset showing the ICL2 outward swings in C5aR2 structures. Lower inset shows Pro138 and Gly139 residues in ICL2 preventing the formation of the ICL2 helix. **(D)** Unique NPxxY and DRY motif in C5aR2. Formation of the C5aR2 intra-helical disulfide bond (indicated in yellow-color) between Cys^3.51^ of the TM3 DLC-motif and Cys^5.57^. Lower-panel showing comparison of DRY(F) (left) and NPxxY (right) motif in hC5aR1 and C5aR2. TM7 NPxxY motif Tyr^7.53^ residue OH-group forms a polar contact with Arg^3.50^ of the DRY motif in hC5aR1, which holds Arg^3.50^ in position to interact with the α5-helix of Go. C5aR2, which instead has an Leu^3.50^ and Phe^7.53^, cannot form polar contacts with each other or with the α5-helix of Go. **(E)** Structure-guided designing of C5aR1 and C5aR2 chimeras, and dose-response curve of G-protein activation as measured by GαoA dissociation and cAMP response. Data represents mean ± SEM, n=3, independent experiments.

While ICL3 and helix 8 were unresolved in the C5aR2 complexes with peptide agonists, they are well resolved in the C5a-C5aR2 complex (**Figure S8**). Structural comparison with the C5a-C5aR1 complex revealed that TM6 in C5aR2 is shortened by five residues, TM5 is shortened by three residues, and the ICL3 region in the C5a-C5aR2 structure is also shortened **(Figure 7B)**. Previous studies suggest that TM5 is structurally important for maintaining the integrity of the G-protein-binding interface. Shortening of TM5 or altering its structure can disrupt the proper positioning of ICL3 and the cytoplasmic cavity, impairing G-protein-coupling^40,57,58^. Furthermore, residues of ICL3 and helix-8 have been found critical for the engagement of GPCRs with the α5 helix of G-proteins^59–67^ (**Figure 7B and S14B**). Dynamic behaviour and potentially occlusive conformational states of ICL3 and helix 8 in C5aR2 are likely to further contribute to hindering the formation of a proper cytoplasmic pocket required for the docking of α5 helix in C5aR2.

The residues corresponding to ICL2 of GPCRs typically adopt a short α-helical turn and contribute to the GPCR-G-protein interface by interacting with a hydrophobic groove formed by α5, αN, β1 and β3 strands of the Gα subunit enhancing binding stability^66,68–70^. The ICL2 residues in the R8Y-, EP54-, and C5a^pep^-bound C5aR2 structures adopt a linearized loop conformation and undergo a linear shift away from the receptor-G-protein interface and also the core of the receptor (**Figure 7C**). Although, the ICL2 residues form a half-helical turn in the C5a-C5aR2 structure, their orientation still remains consistent with the peptide agonist-bound C5aR2, positioning them away from both, the receptor core and the G-proteins (**Figure 7C**). In addition, the presence of a small residue at the end of TM3 (Gly138^3.56^) and proline as the first residue of ICL2 (Pro139^ICL2^) prevent the formation of a kink in ICL2, which in turn limits the ability of ICL2 to adopt a helical conformation, further restraining the engagement with G-proteins^71^ (**Figure 7C**). Therefore, the lack of a kink and the loss of a helical conformation in ICL2 likely alter the spatial positioning of the corresponding residues, thereby weakening the hydrophobic contacts with the α5 helix and polar interactions with the αN helix in G-proteins.

Finally, two of the conserved GPCR motifs namely the DRY motif in TM3 and the NPxxY motif in TM7 display an altered sequence in C5aR2, i.e., DLC and NPxxF, respectively. Interestingly, Cys133^3.53^ (TM3) in the DLC motif forms a disulfide bond with Cys219^5.57^ in TM5, which in turn rigidifies and restrains the movement of TM3 and TM5 in the receptor upon activation (**Figure 7D**). Moreover, in C5a-C5aR1, Arg134^3.50^ in the DRY motif interacts with Tyr300^7.53^ of the NPxxY motif in the active state, which simultaneously engages through an ionic bond with the Cys351 of α5 helix in the G-proteins. In contrast, the Phe300^7.53^ of the NPxxF motif in C5aR2 cannot interact with Leu132^3.50^ of the DLC motif due to the shorter side chain of leucine, and which in turn results in a more flexible cytoplasmic pocket that is inefficient for effective G-protein-coupling (**Figure 7D**). Taken together, the hydrophobicity of the binding cavity, shortening of TM5 and TM6, and the unique DLC motif further contribute to the inability of C5aR2 to couple with, and activate, G-proteins **(Figure 7E-F)**.

Intrigued by the remarkable differences at the cytoplasmic interface of C5aR1 and C5aR2, we hypothesised that substituting the cytoplasmic half of C5aR2 with that of C5aR1 might impart G-protein coupling ability. Based on this, we designed chimeric C5aR1 and C5aR2 constructs in which cytoplasmic halves were reciprocally swapped and subsequently, measured G-protein activation (**Figure 7E**). As anticipated, the C5aR2-C5aR1 chimera, harboring cytoplasmic interface of C5aR1, displayed robust G-protein dissociation and cAMP responses, with only slight shifts in potency and efficacy. This represents first demonstration of a functional gain of G-protein coupling in any ACR. Conversely, the C5aR1 construct bearing the cytoplasmic interface of C5aR2 completely lost G-protein activation, mirroring the behaviour of native C5aR2. Together, these results indicate that the molecular determinants underlying the intrinsic signaling bias of C5aR2 are predominantly localized within its cytoplasmic interface. Collectively, these structural constraints spanning the hydrophobic cavity, shortened TM5 and TM6, and the unique DLC motif, locks C5aR2 into a G-protein incompetent state (**Figure 8A**).

**Figure 8:**
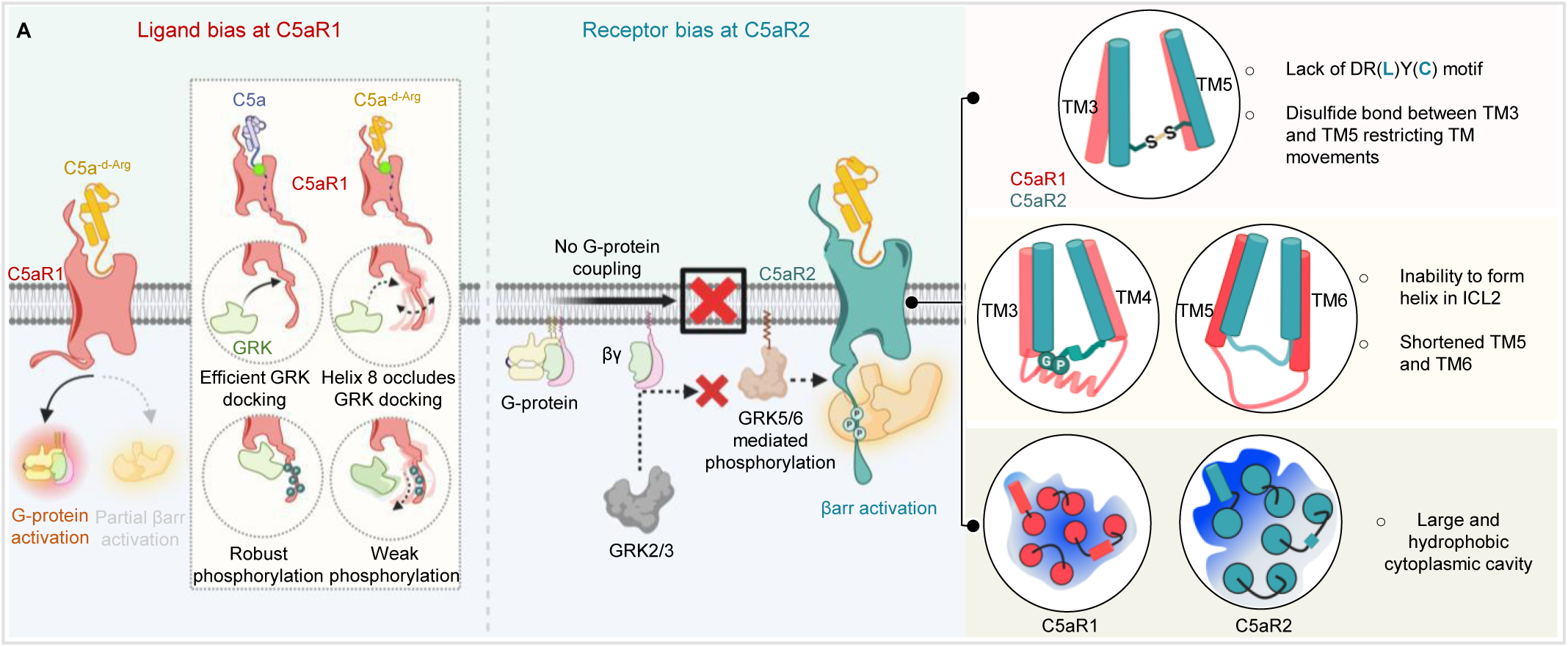
Molecular mechanism of ligand bias and receptor bias at complement anaphylatoxin receptors, C5aR1 and C5aR2. **(A)** Schematic summarizing the structural and functional constraints leading to C5a^-d-Arg^-encoded G-protein bias and intrinsic βarr bias at C5aR1 and C5aR2, respectively. Schematic was prepared in <u>BioRender</u>.

## Discussion

The generation of C3a^-d-Arg^ and C5a^-d-Arg^ is typically conceived as a regulatory mechanism to dampen the excessive inflammatory response, which is believed to arise from a significantly reduced affinity to their corresponding receptors^31–33,43,44,72^. However, our findings presented here, suggest a functional specialization instead, with the three anaphylatoxin receptors perceiving the impact differently. While C3a^-d-Arg^ exhibits near-complete loss of inducing G-protein and βarr-coupling to C3aR, C5a^-d-Arg^ maintains nearly identical activation of G-protein-coupling at C5aR1, but significantly attenuated βarr response. On the other hand, C5a^-d-Arg^ is able to activate βarr-coupling at C5aR2 at levels indistinguishable from C5a. Considering that C5aR2 is weaker than C5aR1 in terms of overall βarr recruitment and completely lacks G-protein-coupling, our data seem to suggest that C5a^-d-Arg^ is encoded to minimize βarr response through C5aR1 and C5aR2 instead of abrogating transducer-coupling entirely. Thus, it is tempting to speculate that G-protein-mediated responses are linked to desirable downstream outcomes and hence maintained via C5aR1, while sustained βarr-mediated responses may be deleterious and hence, minimized via both C5aR1 and C5aR2. In fact, our neutrophil mobilization data in mice where C5a^-d-Arg^ is significantly weaker than C5a, supports such a possibility since excessive C5a-mediated neutrophil mobilization is linked with tissue and organ damage^73–75^. It is also possible that attenuated βarr recruitment at C5aR1 is designed to sustain G-protein signaling via C5aR1, as receptor desensitization through βarrs in response to C5a results in rapid blunting of downstream responses. Probing these intriguing possibilities in future studies should help illuminate the correlation of the naturally-encoded biased-signaling with functional outcomes in physiological and pathophysiological context.

As discussed previously, we have recently observed a striking specialization at the level of species-specific pharmacology of the C3aR and C5aR1^42^. Along the same lines, the C5aR2-selective agonist identified here, also exhibits a strong preference for the human receptor compared to mouse C5aR2. Our previous study has successfully demonstrated that structure-guided receptor mutagenesis can help reverse the species-specific pharmacology, for example, for C5a^pep^ at C5aR1^11^. Therefore, it would be interesting to employ a similar approach to design gain-of-function variants of R8Y that work on mouse C5aR2, and such ligands may help dissect the functional separation of the two receptors in mouse model. In a broader sense, these emerging indications that the human and mouse receptors may have dramatically different pharmacology for natural and synthetic ligands, call for a more careful integration of the concept of species-specific pharmacology in drug discovery campaigns to reduce the disconnect between *in vitro* data and pre-clinical studies.

It is also worth noting that the cryo-EM structures of CXCR7 (ACKR3), which is also a βarr-biased 7TMR, have also been reported recently^71,76^, and the structural interpretation in terms of conformational dynamics of the cytoplasmic interface leading to inefficient G-protein interaction aligns with that of C5aR2 reported here although there are some key differences as well. For example, unlike C5aR2, CXCR7 harbors conserved DRY and NPXXY motifs but still fails to activate G-proteins, and therefore, it is likely that additional mechanisms specific to CXCR7 may also exist rendering it incapable of coupling to G-proteins. Finally, the Duffy antigen receptor for chemokines (DARC), also known as ACKR1, lacks any measurable coupling to either G-proteins or βarrs, which is attributed primarily to shortened TM5 and 6, and a kink at the cytoplasmic portion of TM3^40^, which is significantly different from that observed in C5aR2.

There are still several interesting questions that remain to be answered in the context of ACRs. For example, do the ACRs converge to a common signaling pathway mediated via βarrs? For prototypical GPCRs, agonist-induced activation of ERK1/2 phosphorylation has been used as a quintessential and convergent readout of signaling^77,78^, however, the ACR activation does not appear to effectively elicit this pathway. While future studies focused on deciphering the signaling networks at these receptors will illuminate the downstream signaling aspect further, it is worth noting that there are receptor-specific differences, even at the level of βarr-coupling profile of these ACRs. For example, ACKR2-4 appears to show an agonist-dependent βarr-recruitment while GPR182 (ACKR5) appears to have a constitutive βarr recruitment that is even proposed to decrease in response to selected ligands^12,79,80^. The molecular mechanisms underlying these differences need further exploration at the level of receptor conformation and phosphorylation in cellular context. We also note that structural snapshots provide a static image of the molecules, and the receptor-transducer coupling is likely to be regulated in a more dynamic fashion. Therefore, additional orthogonal approaches to probe the dynamic activation and conformational changes of ACRs may shed additional light in their lack of interaction with G-proteins. Taken together, these observations suggest possible specialization at the level of receptor-specific mechanisms leading to their diverse functional outcomes despite an overall converging thematic connection.

In conclusion, our study elucidates the molecular basis of naturally-encoded ligand-bias and receptor-bias at the complement anaphylatoxin receptors, identifies the first-in-class, C5aR2-selective peptide agonist, and guides the design of signaling-biased C5a variants. Our findings have direct and broad implications for understanding the framework of biased agonism at 7TMRs with direct implications for better therapeutic design.

## Declaration of interest

Authors declare no competing interests.

## Acknowledgements

Research on complement anaphylatoxin receptors in A.K.S.’s laboratory is currently supported by the Senior Fellowship of the DBT Wellcome Trust India Alliance (IA/S/20/1/504916), the Indian Council of Medical research (EMDR/SG/14/2024-01-02127), and the Department of Science and Technology (DST/TTI/TC/AMR/COE/2023/5). Part of the cryo-EM data was collected at the National cryo-EM Facility at IIT Kanpur established with the support from ANRF/SERB (IPA/2020/000405). A.K.S. is the Sonu Agrawal Memorial Chair Professor. This research was also supported by the JSPS KAKENHI grant numbers 21H05037 (O.N.) and 23KJ0491 (F.K.S.), the Platform Project for Supporting Drug Discovery and Life Science Research (Basis for Supporting Innovative Drug Discovery and Life Science Research [BINDS]) from the Japan Agency for Medical Research and Development (AMED) grant numbers JP22ama121012 and JP22ama121002 (O.N.). A.I. was funded by KAKENHI JP21H04791 and JP24K21281 from the Japan Society for the Promotion of Science (JSPS); JP22ama121038 and JP22zf0127007 from the Japan Agency for Medical Research and Development (AMED); JPMJFR215T and JPMJMS2023 from the Japan Science and Technology Agency (JST); The Uehara Memorial Foundation. Research in T.M.W.’s laboratory is supported by the National Health and Medical Research Council (APP2009957) and R.J.C.’s research is supported by the National Health and Medical Research Council (APP2012661). HDX-MS work in K.Y.C.’s laboratory was supported by grants from the National Research Foundation of Korea funded by the Korean government (NRF-2021R1A2C3003518 and NRF-2019R1A5A2027340 to K.Y.C.). We also acknowledge Australian Red Cross Lifeblood and human donors for providing blood for our research. We also thank Kayo Sato, Shigeko Nakano and Ayumi Inoue in the Inoue lab for their assistance in the plasmid construction and the NanoBiT assay. We sincerely thank Dr. Charles Mackay and Caroline Ang for providing the 1D9 and 4C8 hybridoma clones. We thank Shachie Sinha for helping with cellular assays, and Ashna Reyaz, Calvin D’Souza, and Debdatta Mukherjee with protein purification.

## Authors’ contribution

DT and MKY expressed and purified the receptor and prepared the complexes with help from AD; DT expressed and purified 4C8 antibody and prepared the Fab with help from NR in the early stages; KS and FKS screened the samples for cryo-EM, collected and processed the data with help from KY, HSO, KH, and built the initial model under the supervision of ON; AD and SM carried out the functional assays in HEK-293 cells; XXL carried out the Ca^2+^ flux, pERK, and BRET assays, and experiments in primary HMDMs/BMDM/PMNs, under the supervision of TMW; CSC carried out the *in vivo* neutrophil mobilisation assay with help from JNF and TL under the supervision of JDL and TMW. JCD identified and characterized R8Y under the supervision of RC and TMW; KK and DA performed and analyzed the HDX experiments under the supervision of KYC; NB expressed and purified wild-type C5a and C5a^-d-Arg^; TMS performed the MD simulation studies under the supervision of JS; AI carried out the C5aR1-GRK interaction assay; RB and MKG also processed cryo-EM data of apo-C5aR2 and R8Y-C5aR2 collected at IITK, refined and built the final models for all the cryo-EM structures, carried out the analyses, and helped prepare the figures together with DT and NR; RC, KYC, RB, FKS, TMW, ON, and AKS supervised the overall project. All authors contributed to writing and editing the manuscript.

## Data availability

The cryo-EM maps and structures have been deposited in the EMDB and PDB with accession numbers PDB ID-9V3C, EMD-64752 (Trx-C5a-C5aR2), PDB ID-9WDI, EMD-65890 (mC5a^-d-Arg^-mC5aR2), PDB ID-9V35 , EMD-64749 (Fab4C8-Apo-C5aR2), PDB ID-9V3Y, EMD-64761 (Fab4C8-C5a^pep^-C5aR2), PDB ID-9V38, EMD-64751 (Fab4C8-EP54-C5aR2) and PDB ID-9V4D, EMD-64777 (Fab4C8-R8Y-C5aR2).

## Materials and methods

### General chemicals and reagents

Most of the general reagents were purchased from Sigma-Aldrich unless otherwise specified. Dulbecco’s Modified Eagle’s Medium (DMEM), Trypsin-EDTA, Fetal Bovine Serum (FBS), Phosphate-Buffered Saline (PBS), Hanks’ Balanced Salt Solution (HBSS), and Penicillin-Streptomycin solution were obtained from Thermo Fisher Scientific. HEK-293T cells (ATCC) were maintained in DMEM (Gibco, Cat. No: 12800-017) supplemented with 10% (v/v) FBS (Gibco, Cat. No: 10270-106) and 100 U/mL penicillin and 100 μg/mL streptomycin (Gibco, Cat. No: 15140122) at 37 °C under 5% CO₂. *Sf9* cells were cultured in protein-free media (Gibco, Cat. No: 10902-088) at 27 °C at 135 rpm. The cDNA coding regions of C3aR, C5aR1 and C5aR2 were cloned into the pcDNA3.1 vector with an HA signal sequence and an N-terminal FLAG-tag followed by a TEV site and the receptor sequence. For expression into *Sf9* system, the cDNA of C5aR2 was cloned into pVL1393 vector harbouring N-terminal FLAG-tag, followed by N-terminus of the M4 receptor (residues 2-23), synthesized by GenScript. The constructs used in the NanoBiT-based β-arrestin recruitment assay were cloned into the pCAGGS vector, with SmBiT fused to the receptor’s C-terminus and LgBiT fused to the N-terminus of βarr1/2, as previously described^81^. The G-protein subunit constructs used in the dissociation assays were generously provided by Asuka Inoue. SRF and SRE reporter gene plasmids were purchased from Promega (Cat. No: E150 for both SRE and SRF). C5aR1 and C5aR2 mutants were generated by using the Q5^®^ Site-Directed Mutagenesis Kit (NEB, Cat. No: E0554S). All DNA constructs were verified by sequencing at Macrogen. Peptides, EP54, C5a^pep^, EP67, EP141, P32 and P59 were synthesized by GenScript. Antibodies were purchased from Sigma-Aldrich (M2-HRP coupled anti-FLAG), GenScript (HRP-coupled anti-rabbit), or Cell Signaling Technology (ERK1/2), pIMAGO kit purchased from Sigma-Aldrich (Cat No. 18419), hC5aR1 phosphorylation specific antibodies pT324/pS327 and p332/pS334 purchased from 7TM antibodies (Cat No. 7TM0032A and 7TM0032B, respectively).

### Human cell line

HEK-293T cells were procured from ATCC and regularly monitored under bright-field microscope for proper morphology, however the examination of mycoplasma contamination was not performed. The cell line was maintained in DMEM supplemented with 10% FBS, 100 U/mL penicillin and 100 µg/mL streptomycin, at 37 °C in 5% humidified CO2 incubator. The cells were maintained at 70-80% confluency either in T175 flasks or 10 cm cell-culture treated round dishes and sub-cultured every alternate day.

### Chinese hamster ovary (CHO-cells)

Chinese hamster ovary cells stably expressing either C3aR (CHO-C3aR) or C5aR1 (CHO-C5aR1) were maintained in Ham’s F12 media supplemented with 10% FCS, 100 U/mL penicillin, 100 µg/mL streptomycin and 400 µg/mL G418 (Invivogen, San Diego, USA). The cell line was maintained in T175 flasks (37 °C, 5% CO2) and sub-cultured at 90% confluency using TrypLE Express (Thermo Fisher Scientific, Melbourne, Australia).

### Insect cell line

*Spodoptera frugiperdA (Sf9)* cells were obtained from Expression systems. The cells were maintained in glass conical flasks at a density of 0.9 million cells per mL in protein-free insect cell medium with regular splitting at every alternate day. These cells were grown in a shaker incubator at 27 °C with a constant agitation at 135 rpm.

### Bacterial cell culture

*Escherichia coli* strain DH5alpha were used for plasmid DNA amplification and isolation, and they were cultured in Luria-Bertani (LB) broth at 37 °C with shaking at 160 rpm. For protein expression, BL21 (DE3), Rosetta (DE3), SHuffle strains of *Escherichia coli* were used, and they were cultured using Luria-Bertani (LB), Terrific Broth (TB), or 2XYT media under the indicated culture conditions (temperature and shaking) as described in the subsequent method sections.

### Primary Cell culture

Human monocyte-derived macrophages (HMDMs) were generated and cultured as previously described^82^, with experiments approved by The University of Queensland Human Research Ethics Committee. Briefly, human buffy coat blood from anonymous healthy donors was obtained through the Australian Red Cross Blood Service (Brisbane, Australia). Human CD14+ monocytes were isolated from blood using Lymphoprep density centrifugation (STEMCELL, Melbourne, Australia) followed by CD14+ MACS magnetic bead separation (Miltenyi Biotec, Sydney, Australia). The isolated monocytes were differentiated for 7 days in Iscove’s Modified Dulbecco’s Medium supplemented with 10% FBS, 100 IU/mL penicillin, 100 μg/mL streptomycin and 15 ng/mL recombinant human macrophage colony stimulating factor (BioLegend, San Diego, USA) on 10 mm square dishes (Bio-strategy, Brisbane, Australia). Non-adherent cells were removed by washing with DPBS, and the adherent differentiated HMDMs were harvested by gentle scraping.

Mouse bone marrow-derived macrophages (BMDMs) were obtained and cultured as previously described^83,84^. Briefly, mice were sacrificed by cervical dislocation. The tibia was removed and sterilised. Upon removal of both epiphyses, bone marrow cells were harvested by flushing the central cavity with complete RPMI-1640 medium using a 10 mL syringe attached to a 25-gauge needle. Cells were then cultured in complete RPMI-1640 medium (containing 10% FBS, 100 IU/mL penicillin, 100 μg/mL streptomycin) supplemented with 100 ng/mL recombinant mouse macrophage colony stimulating factor on 10 mm square dishes (Thermo Fisher Scientific, Melbourne, Australia). Mature adherent macrophages for assays were harvested on day 6 by gentle scraping.

### NanoBiT-based GoA dissociation assay

Ligand-induced G-protein activation was measured using a previously described NanoBiT-based G-protein dissociation assay^85^. Briefly, HEK-293T cells were transiently transfected with G-protein subunits harbouring LgBiT-tagged Gα subunit (1 µg), SmBiT-tagged Gγ2 (C68S mutation) subunit (4 µg) and untagged Gβ1 subunit (pcDNA3.1) (4 µg), along with the N-terminal FLAG-tagged human (h) and mouse (m) C5aR1 receptor. After 14-16 h of transfection, the cells were trypsinized and harvested using Trypsin-EDTA and resuspended in NanoBiT buffer (5 mM HEPES, pH 7.4, 1x HBSS, 0.01% BSA, and 10 µM coelenterazine (Gold Bio, Cat. No: CZ5). 100 μL of resuspended cells were then seeded into a 96-well plate at a density of 0.1 million cells per well and incubated at 37 °C for 90 min, followed by an additional 30 min incubation at room temperature. Basal luminescence was recorded for three cycles using a Fluostar Omega plate reader. Subsequently, the cells were stimulated with varying doses of ligand and decrease in luminescence was recorded as a functional readout of G-protein activation for 10 cycles. For analysis, response recorded at 10 min was basal corrected and % decrease in luminescence as a function of ligand concentration was plotted by normalising luminescence values with respect to the lowest ligand dose taken as 100%. The resulting data was plotted using GraphPad Prism 10.3.1 software, undertaking non-linear regression curve-fitting.

### GloSensor assay to measure cAMP response

To measure ligand-induced change in intracellular cAMP levels, GloSensor assay was employed as previously described^86^. Briefly, HEK-293T cells were transiently transfected with 3.5 μg of either, hC5aR1 or mC5aR1 (cloned in the pcDNA vector having N-terminus FLAG-tag), along with 3.5 μg of the F22 plasmid (Promega, Cat. No: E2301). After 14–16 h of transfection, cells were trypsinized and resuspended in an assay buffer comprising 20 mM HEPES, pH 7.4, 1x HBSS, and 0.5 mg/mL D-luciferin (GoldBio, Cat. No: LUCNA-1G). Following this, 100 µL of transfected cells were seeded into a 96-well plate at a density of 0.2 million cells per well. The cells were incubated at 37 °C for 90 min, followed by an additional 30 min incubation at room temperature. Basal luminescence was measured for three cycles. Subsequently, 5 μM forskolin was added to each well, and forskolin induced increase in luminescence was recorded till saturation, i.e., for 8 cycles. Finally, the ligand was added at the indicated doses, and ligand induced decrease in luminescence was recorded for 15 cycles. For data analysis, ligand-mediated decrease in cAMP response was normalised with forskolin-induced luminescence and the percentage decrease in cAMP (% normalisation) was calculated by normalising the values of each dose with the smallest ligand dose taken as 100%. Curve fitting was done by non-linear regression curve-fitting, using GraphPad Prism

#### 10.3.1 software

### NanoBiT-based β-arrestin1/2 recruitment

To measure agonist induced β-arrestin1/2 recruitment downstream to (h/m) C5aR1 and (h/m) C5aR2, NanoBiT-based enzyme complementation assay was employed, as described previously^85^. Briefly, HEK-293T cells were transiently transfected with either 3.5 µg hC5aR1/ 3.5 µg mC5aR1/ 5 µg hC5aR2/ 0.1 µg mC5aR2, harbouring SmBiT fragment at carboxyl terminus and β-arrestin1/2, 3.5 µg (for h/mC5aR1) / 2 µg (for h/mC5aR2), harbouring LgBiT fragment at N-terminus using polyethylenimine (PEI) at 1:3 (DNA: PEI) ratio. Post 16-18 h, cells were harvested using Trypsin-EDTA and resuspended in NanoBiT assay buffer containing 1x HBSS, 5 mM HEPES, pH 7.4, 0.01% BSA and 10 µM coelenterazine. 0.1 million cells were seeded in each well of a 96 well plate and incubated at 37 °C in a 5% humidified CO2 incubator for 90 min. Subsequently, the plates were incubated at room temperature for an additional 30 min, following this the basal luminescence was recorded for 3 cycles using a plate reader (LUMIStar Omega, BMG LABTECH). The cells were then stimulated with varying ligand doses prepared in NanoBiT drug buffer (1x HBSS and 5 mM HEPES, pH 7.4) and increase or decrease in luminescence was recorded for 12 cycles. The average luminescence for 6 cycles (5-10 cycles) was normalized with respect to luminescence recorded for lowest ligand dose taken as 1. Fold response was plotted as a function of logarithmic dose of ligand using GraphPad Prism 10.3.1 software. Bias factor was calculated by using https://biasedcalculator.shinyapps.io/calc/.

### Receptor surface expression

The surface expression of the receptors was quantified using whole cell-surface ELISA assay, as described previously^87^. Briefly, the transiently transfected cells were seeded in poly-D-lysine coated 24-well plate at a density of 0.2 million cells per well and incubated at 37 °C in 5% humidified CO2 incubator. Post 24 h, cells were washed with 1x Tris-Buffer Saline (TBS) and fixed with 4% paraformaldehyde (PFA) (w/v prepared in 1x TBS) for 20 min followed by washing with 1X TBS, three times, to completely remove the traces of PFA. Subsequently, the cells were incubated with 1% BSA (prepared in 1x TBS) for 1 h followed by incubation in anti-FLAG M2-HRP antibody (at 1:10,000 dilution; prepared in 1% BSA) (Sigma, Cat. No. A8592) for another 1 h. After 1 h, the cells were washed three times with 1% BSA to remove excess antibody and ELISA was developed using TMB-ELISA substrate (Thermo Scientific, Cat. No: 34028). For the same, the cells were incubated with 200 µL TMB-ELISA substrate until light blue colour appeared. The reaction was quenched by adding 100 µL of above coloured solution into 100 µL 1M H2SO4 and the resultant yellow colour intensity was measured by reading the absorbance at 450 nm using a multi-mode plate reader (PerkinElmer Victor^TM^ X4). The cell count was quantified using Janus-green stain. For this, the TMB substrate was removed by washing the cells with 1x TBS and incubated with 0.2% (w/v) Janus green B stain (Sigma, Cat. No: 201677) stain for 10 min. The excess stain was removed by washing the cells with MilliQ water. The retained dark-blue colour was dissolved by adding 800 µL of 0.5 N HCl and absorbance was recorded at 595 nm. To quantify the receptor surface expression, the ratio of absorbance at 450 nm and 595 nm was taken and normalized with respect to the mock (pcDNA) transfected cells and plotted using GraphPad Prism 10.3.1 software.

### Intracellular calcium mobilisation assays

Ligand-induced intracellular calcium mobilisation was assessed using Fluo-4 NW Calcium Assay kit following the manufacturer’s instructions (Thermo Fisher Scientific, Melbourne, Australia), as described previously^11^. Briefly, HMDMs (70,000 per well) or BMDMs (90,000 per well) were seeded in black clear-bottom 96-well tissue culture plates overnight. Cells were firstly stained with the Fluo-4 dye in assay buffer (1x HBSS, 20 mM HEPES) for 50 min (37 °C, 5% CO2). C5a/C5a^-d-Arg^ dilutions were prepared in assay buffer containing 0.1% BSA. On a Flexstation 3 platform, the fluorescence (Ex/Em: 494/516 nm) was continually monitored for a total of 100 s, with ligand addition performed at 16 s. Data were recorded as the magnitude of signal deviation from the baseline.

### Measuring ERK and RhoA signaling

For measuring ERK signaling downstream to stimulation of hC5aR1 and mC5aR1 with ligands, we undertook an SRE reporter assay^88^. HEK-293T cells were transfected with 3.5 μg of N-terminally FLAG-tagged hC5aR1/mC5aR1 and 3.5 μg of an SRE-based luciferase reporter plasmid pGL4.33 (Promega, Cat. no: E1340). 14-16 h post-transfection, cells were washed with 1x PBS, trypsinized and seeded into a 96-well plate at a density of 1 x 10^6^ cells per well in the presence of complete media. Cells were allowed to settle for 8 h, followed by starvation in serum-deprived DMEM (without FBS), for 12 h. Subsequently, the cells were stimulated with the indicated dose of the ligands (prepared in serum-free DMEM) and the plates were incubated at 37 °C for 3 h. Prior to reading, the serum-free media was carefully replaced with the assay buffer containing 20 mM HEPES pH 7.4 and 1x HBSS supplemented with 0.5 mg/mL D-luciferin. Luminescence was recorded immediately. Signal observed was normalized with respect to the luminescence observed at lowest concentration of each ligand, taken as 1. Data was plotted and analyzed using GraphPad Prism 10.3.1 software.

For measuring RhoA signaling, HEK-293T cells were transfected with 3.5 μg of N-terminally FLAG-tagged receptor and 3.5 μg of an SRF-based luciferase reporter plasmid pGL4.34 (Promega, Cat. no: E135A) followed by the same steps as that in SRE reporter assay.

The ligand-induced ERK1/2 phosphorylation in HMDM and BMDM was assessed using the AlphaLISA *Surefire Ultra* p-ERK1/2 (Thr202/Tyr204) kit (Revvity, Melbourne, Australia) as previously described^89^. Briefly, HMDMs (50,000/well) or BMDMs (90,000/well) were seeded in tissue culture-treated 96-well plate for 24 h and serum-starved overnight. Human and mouse (h/m) C5a/C5a^-d-Arg^ dilutions were prepared in serum-free medium containing 0.1% BSA and added to the cells (10 min for HMDMs; 5 min for BMDMs). After stimulation, cells were immediately lysed using AlphaLISA lysis buffer on a microplate shaker (450 rpm, 10 min). For the detection of phospho-ERK1/2 content, cell lysate (5 μL/well) was transferred to a 384-well ProxiPlate (Revvity) and added to the donor and acceptor reaction mix (2.5 μL/well, respectively) with 2 h incubation at room temperature in the dark. The plate was read on Tecan Spark 20M following standard AlphaLISA settings.

For measuring agonist-induced ERK1/2 phosphorylation in C5aR1-expressing HEK-293 stable cell line, a previously described Western blotting–based protocol was employed^90^. Briefly, hC5aR1 expressing stable cell lines were seeded into a 6-well plate at a density of 1 million cells per well. The cells were serum-starved for 12 h followed by stimulation with 1 μM concentration of hC5a and hC5a^-d-Arg^, at selected time points. After the stimulation, the medium was aspirated, and the cells were lysed in 100 μL of 2x SDS dye per well. The cells were heated at 95 °C for 15 min, followed by centrifugation at 15,000 rpm for 15 min. 10 μL of lysate was loaded per well and separated on SDS-PAGE, followed by Western blotting. The blots were blocked in 5% BSA (in 1x TBST) for 1 h and incubated overnight with rabbit-raised phospho-ERK (Cat no. 9101/CST) primary antibody at 1:5000 dilution. Following this, the blots were washed thrice with 1x TBST for 10 min each and incubated with anti-rabbit HRP-coupled secondary antibody (1:10,000, Cat no. A00098/GenScript) for 1 h. The blots were washed again with 1x TBST three times and developed with Promega ECL solution (Cat. No: W1015) using ChemiDoc (Bio-Rad). The blots were stripped with a low pH stripping buffer and then re-probed for total ERK using rabbit-raised p44/42 MAPK (Erk1/2) Antibody (Cell Signalling Technology, Cat. No: 9102) primary antibody at 1:5000 dilution. Blots were again incubated with anti-rabbit HRP-coupled secondary antibody for 1 h after washing three times with 1x TBST buffer. ERK phosphorylation signal was obtained after quantification with ImageJ software and fold normalized values were plotted using GraphPad Prism 10.3.1 software.

For measuring ligand pERK1/2 signaling in response to BM2020-7 and BM2020-8 analogues, ligand-induced phospho-ERK 1/2 signaling was assessed using the Alpha LISA SureFire Ultra pERK 1/2 (Thr202/Tyr204) assay kit (PerkinElmer, Melbourne, Australia). Either CHO-C3aR or CHO-C5aR1 cells were seeded (50,000 cells per well) onto 96-well tissue culture-treated plates, incubated for 24 h and subsequently serum starved overnight. Ligand dilutions were prepared in serum-free medium. Cells were stimulated with respective ligands for 10 min and then immediately lysed using AlphaLISA lysis buffer. Cell lysate (5 µL per well) was added to a 384-well ProxiPlate (PerkinElmer, Melbourne, Australia) followed by the donor and acceptor reaction mixes (2.5 µL per well each). Following a 2 h incubation in the dark, the plate was read on a CLARIOstar Plus microplate reader following standard AlphaLISA settings. Experiments were conducted in triplicate and conducted on at least three different days. Data were analysed using GraphPad software (Prizm 10.1) and expressed as mean ± standard error of mean (SEM). For each repeat, data was normalised prior to being combined. Logarithmic concentration-response curves were plotted using combined data and analysed to calculate the potencies of each peptide.

### Measurement of IL8 release using ELISA

To compare the ability of the C5a species to induce IL8 release from HMDMs^82^, HMDMs were seeded in 96-well tissue culture plates (90,000 per well) for 24 h before treatment. Cells were stimulated with various concentrations of human plasma-derived or recombinant C5a/ C5a^-d-Arg^ for 24 h (37 °C, 5% CO2) before the supernatant was collected. The IL8 level in the supernatant was quantified using human IL8 enzyme-linked immunosorbent assay (ELISA) kit (BD OptEIA) as per manufacturer’s protocol.

### PMN chemotaxis assay using FluroBlok

C5a-induced chemotaxis of human polymorphonuclear leukocytes (PMNs) were assessed using Corning^®^ FluoroBlok™ HTS 96-well Multiwell Permeable Support System (Corning, New York, USA). PMNs were obtained from venous whole blood (20 mL) collected from healthy volunteers under informed consent. Samples were collected using venepuncture into BD K2EDTA Vacutainer® blood collection tubes (BD Biosciences, Macquarie Park, Australia) and processed within 5 h. For PMN isolation, the anticoagulated blood was firstly layered over a Lymphoprep (STEMCELL, Melbourne, Australia) density gradient and then centrifuged (800xg, 30 min, 22 °C), followed by residual erythrocytes removal using hypotonic lysis^91^. Isolated PMNs were counted and resuspended in HBSS, 20 mM HEPES, 0.5% BSA migration buffer (2 x 10^6^ per mL). Calcein AM (2 µM) was added to label the cells (30 min, 37 °C). The cells were then gently washed once with HBSS buffer and added to the insert (2 x 10^5^ per insert). To initiate cell migration, C5a/C5a^-d-Arg^ prepared in the migration buffer were added to the receiver wells. On a Tecan Spark 20M microplate reader (Tecan, Männedorf, Switzerland) (37 °C), ligand-induced cell migration was continuously monitored at 2 min intervals for 40 min by quantifying Calcein AM fluorescence from the receiver-side of the insert (Ex/Em = 485 nm/525 nm). The relative cell migration (fold-baseline) at 20 min post ligand addition was used for graphing.

### BRET assay measuring β-arrestin recruitment to C5aR1

The C5a-mediated β-arrestin recruitment to C5aR1 was measured using bioluminescence resonance energy transfer (BRET)-based assay using methods described elsewhere^83,89^.

Briefly, HEK-293 cells were transiently transfected with human C5aR1-Rluc8 and β-arrestin 1/2-venus constructs using XTG9 (Merck, Melbourne, Australia) for 24 h. Transfected cells were then seeded (100,000 per well) onto white 96-well plates (Corning, New York, USA) in phenol-red free DMEM containing 5% FBS overnight. For BRET assay, cells were first incubated with the substrate Enduren (30 µM, Promega, Sydney, Australia) for 2 h (37 °C, 5% CO2). On a Tecan Spark 20M microplate reader (Tecan, Männedorf, Switzerland) maintained at 37 °C, the BRET light emissions (460-485 and 520-545 nm) were continuously monitored for 90 min with C5a/C5a^-d-Arg^ added after 15 min. The ligand-induced BRET ratio was calculated by subtracting the emission ratio of Venus (520-545 nm)/Rluc8 (460-485 nm) of the vehicle-treated wells from that of the ligand-treated wells. The data at 40 min post ligand addition was used for derivation of concentration-response curves.

For measuring β-arrestin 2 recruitment in response to BM2020-7 and BM2020-8 analogous peptides, HEK-293 WT cells were transiently transfected with C5aR1-*Renilla* Luciferase 8 (Rluc8) and β-arrestin 2-Venus or C5aR2-Venus and β-arrestin 2-Rluc constructs using X-tremeGENE 9 DNA Transfection Reagent (Roche, Sydney, Australia). The following day, adherent cells were detached using TrypLE Express and seeded plates (100,000 cells per well) onto white 96-well tissue culture-treated plates (Corning, NY, USA) in phenol-red free DMEM supplemented with 5% FBS. On the next day, the cells were incubated with either Enduren (C5aR1, Promega, Sydney, Australia) or Endurazine (C5aR2, Promega) substrates diluted in assay media for 2 h (37 °C, 5% CO2). BRET emissions (460-485 nm and 520-545 nm) were measured using either a Tecan Spark 20M or PHERAstar FSX microplate reader (37 °C) for 19 reads, with respective ligands added after the first 4 reads. The ligand induced BRET ratio was calculated by subtracting the Venus/Rluc8 ratio of the negative control wells from that of the ligand treated wells.

### Neutrophil mobilisation in mouse

Eight-to ten-week-old C57BL/6J mice (n=4 per group; Ozgene, Australia) received a single intraperitoneal dose of the C5aR1 receptor antagonist PMX205 (3 mg/kg) using a concentration of 1.5 mg/mL in 5% (w/v) dextrose (Baxter, cat. no. AHB0063) or an equal volume of vehicle (5% (w/v) dextrose). Fifteen minutes later, recombinant mC5a or mC5a^-d-Arg^ (produced in-house) diluted in sterile 0.9% (w/v) sodium chloride for injection (Pfizer, Cat. no. 1016696) was administered intravenously via the lateral tail vein at 50 µg/kg. Peripheral blood (20 µL) was collected from the tail tip of mice at 0 min and at 60 min after C5a challenge, then mixed with 480 µL V-52D diluent (Mindray, Cat. no. 105-005962-00). Absolute neutrophil counts were measured using an automated haematology analyser (BC-5000; Mindray, Shenzhen, China). All animal procedures were approved by the University of Queensland Animal Ethics Committee and performed in accordance with ARRIVE guidelines.

### MD simulation

Receptor dynamics-driven important functional features of GPCRs^47,48^ have been simulated for the C5aR1 using the AceMD engine^92^. Receptors (PDB code: 8IA2, 8JZZ) were prepared in MOE software (www.chemcomp.com). Systems were generated in Charmm-GUI software^93^ with parameters from the CharMM36M forcefield^94^. Each mutated system was generated with Charmm-GUI^93^, using the structure of the C5a-bound human C5aR1 (PDB code: 8IA2) as a starting point. The complexes were solvated (TIP3P water) and neutralized using a 0.15 concentration of NaCl ions. All systems underwent 100 ns equilibration in conditions of constant pressure (NPT ensemble, pressure maintained with Berendsen barostat, 1.01325 bar pressure), using a timestep of 2 fs. During this stage restraints were applied to the protein and ligand backbone. This was followed by 3 separate NVT runs for each system, 1 µs each. For each of the simulations we used a temperature of 310 K, which was maintained using the Langevin thermostat, hydrogen bonds were restrained using the RATTLE algorithm. Non-bonded interactions were cut-off at a distance of 9 Å, with a smooth switching function applied at 7.5 Å. The simulation data have been uploaded to the GPCRmd repository^48^: https://gpcrmd.org/dynadb/publications/XXXX/.

### Hydrogen-Deuterium exchange-mass spectrometry (HDX-MS)

mC5aR1 and C5aR2 samples were prepared at 100 µM in 20 mM HEPES pH 7.4, 150 mM NaCl, and 0.01% L-MNG. The protein samples (3.4 µL) were mixed with 26.6 µL D2O buffer (20 mM HEPES pH 7.4, 150 mM NaCl, and 10% glycerol), incubated for 10, 100, 1000, and 10000 s at room temperature (23-25 °C), and quenched with 30 µL of ice-cold quench buffer (60 mM NaH2PO4 pH2.01, 20 mM TCEP, 2 M GuHCl). The quenched samples were then snap-frozen on dry ice and stored at -80 °C. For non-deuterated samples, 3.4 µL of protein samples were mixed with 26.6 µL of their respective H2O buffers and quenched with the 30 µL quench buffer.

The quenched samples were thawed and immediately digested by passing through an immobilized pepsin column (2.1 x 30 mM) (Life Technologies, Carlsbad, CA, USA) at a flow rate of 100 µL/min in 0.05% formic acid in H2O at 12 °C. The peptic fragments were collected on a C18 VanGuard trap column (1.7 µM x 30 mM) (Waters) for desalting with 0.05% formic acid in H2O. The peptic fragments were then separated by an Acuity UPLC C18 column (1.7 µm, 1.0 x 100 mM) (Waters) at a flow rate of 40 µL/min in mobile phase A (0.1% formic acid in H2O) with an acetonitrile gradient increase starting from 8% to 85% over 8.5 min with mobile phase B (0.1% formic acid in acetonitrile). To minimize the back-exchange, the buffers were adjusted to pH 2.5, and the analysis was performed at 0.5 °C. Mass spectral analyses were performed using a Xevo G2 quadrupole time-of-flight (Q-TOF) equipped with a standard ESI source in MSE mode (Waters) in positive ion mode. The capillary, cone, and extraction cone voltages were set to 3 kV, 40 V, and 4 V, respectively. The source and desolvation temperatures were set at 120 °C and 350 °C, respectively. Trap and transfer collision energies were set to 6 V; the trap gas flow was 0.3 mL/min. Before analysis, the mass spectrometer was calibrated by sodium iodide (2 µg/µL). [Glu1]-Fibrinopeptide B (200 fg/µL) in MeOH:water (50:50 (v/v) + 1% acetic acid) was utilized for lock-mass correction. The ions at mass-to-charge ratio (m/z) of 785.8427 were monitored at a scan time of 0.1 s with a mass window of ± 0.5 Da. The reference internal calibrant was introduced into the lock-mass sprayer at a flow rate of 20 µL/min, and all spectra were automatically corrected. Two independent interleaved acquisition functions were created: the first function, typically set at 4eV, collected low-energy or unfragmented data, and the second function collected high-energy or fragmented data typically obtained using a collision ramp from 30-55 eV. Mass spectra were acquired in the range of m/z 100-2000 for 10 min. The peptides from non-deuterated samples were identified with ProteinLynx Global Server 2.4 (Waters), and the level of deuterium uptake for each peptide was determined by measuring the centroid of the isotopic distribution with DynamX 3.0 (Waters). Back-exchange was not corrected because we observed protein aggregates when preparing the fully deuterated samples.

### NanoBiT-based GRK recruitment assay

Ligand-induced GRK recruitment to C5aR1 was measured by a NanoBiT-GRK assay^85^ with minor modifications. Specifically, HEK-293A cells (Thermo Fisher Scientific) were seeded in a 6-well culture plate at a density of 0.2 million cells per mL (2 mL per well in DMEM (Nissui) supplemented with 5% FBS, glutamine, penicillin, and streptomycin) one day before transfection. The transfection solution was prepared by mixing 5 µL (per well) of polyethyleneimine (PEI) Max solution (1 mg/mL; Polysciences), 200 µL of Opti-MEM (Thermo Fisher Scientific), and a plasmid mixture containing 500 ng of sHA-FLAG-C5aR1-LgBiT and 500 ng of GRK-SmBiT. After a day of incubation, the transfected cells were harvested with Dulbecco’s PBS containing 0.5 mM EDTA, centrifuged, and resuspended in 3 ml of HBSS containing 0.01% bovine serum albumin (BSA; fatty acid-free grade; SERVA) and 5 mM HEPES, pH 7.4 (assay buffer). The cell suspension was dispensed into a white 96-well plate at a volume of 80 µL per well, and 20 µL of 50 µM coelenterazine (Angene) diluted in the assay buffer was added. After a 2 h incubation at room temperature, baseline luminescence was measured using a SpectraMax L (Molecular Devices), and a titrated test ligand (20 µL; 6x of final concentrations) was manually added. The plate was immediately read at room temperature in kinetics mode for 15 min, with measurements taken every 20 s. Luminescence counts from 5–10 min after ligand addition were averaged and normalized to the initial counts. The fold-change values were further normalized to those of vehicle-treated samples and used to plot the GRK recruitment response. GRK recruitment signals were fitted to a four-parameter sigmoidal concentration-response curve using Prism 10 software (GraphPad Prism 10.3.1 software). For each replicate experiment, the parameter *Span (= Top – Bottom)* for individual ligands was normalized to acetylcholine, and the resulting *Emax* values were used as a measure of efficacy.

### Receptor phosphorylation measurement via pIMAGO assay

Agonist-induced phosphorylation downstream of mC5aR1 was measured by using pIMAGO phosphoprotein detection kit from Sigma (Cat. No. 18419), following the manufacturer’s protocol. Briefly, *Sf9* cells were co-infected with baculovirus expressing mC5aR1 and either GRK2 or GRK6 at a density of 1.8 million cells per mL. 72 h post-infection, cells were stimulated with either 1 µM mC5a or mC5a^-d-Arg^ for 30 min at 37 °C. Following stimulation, cells were harvested by centrifugation at 5,000 rpm for 10 min. The harvested pellets were processed for lysis, and cells were dounce-homogenized in lysis buffer containing 20 mM HEPES, pH 7.4, 150 mM NaCl, 1x PhosSTOP (Roche, Cat. No. 57084100), and 1x protease inhibitor cocktail (Roche, Cat. No. 04693116001). Lysates were solubilized in 1% (w/v) L-MNG (Cat. No. NG31025GM) at room temperature for 1 h and centrifuged at 15,000 rpm for 10 min. The cleared lysate was transferred to a separate tube containing pre-equilibrated M1-FLAG beads supplemented with 5 mM CaCl₂. Samples were incubated at room temperature for 90 min with gentle tumbling to allow bead binding. The beads were washed five times with low-salt buffer (20 mM HEPES, pH 7.4, 150 mM NaCl, 2 mM CaCl₂, and 0.01% L-MNG) alternated with high-salt buffer (20 mM HEPES, pH 7.4, 350 mM NaCl, 2 mM CaCl₂, and 0.01% L-MNG). Bound proteins were eluted using FLAG-elution buffer containing 20 mM HEPES, pH 7.4, 150 mM NaCl, 2 mM EDTA, 0.01% L-MNG, and 250 µg/mL FLAG peptide. Subsequently, protein loading dye was added to each sample, followed by the addition of 5x IAA solution to a final 1x concentration from the pIMAGO kit. The samples were incubated at room temperature for 15 min in the dark. After incubation, samples were subjected to SDS-PAGE, followed by western blotting. The PVDF membrane was blocked in 1x blocking buffer overnight, then incubated with the pIMAGO reagent (1:1000, prepared in 1x pIMAGO buffer) for 1 h. The membrane was washed three times with 1x wash buffer (prepared from 10x stock) and once with 1x TBST (for 5 min each). The PVDF membrane was then incubated with avidin-HRP (1:1000, prepared in 1x blocking buffer) for 1 h at room temperature, followed by three washes with 1x TBST (5 min each). The signal was detected using the Promega ECL solution on a ChemiDoc imaging system (Bio-Rad). The PVDF membrane was subsequently stripped and re-probed for total receptor levels using HRP-conjugated anti-FLAG M2 antibody (Sigma, 1:5000). Phosphorylation and receptor signals were quantified using ImageLab software (Bio-Rad). Fold-normalized values were plotted using GraphPad Prism 10.3.1 software.

### Detection of phosphorylation status of receptor via hC5aR1 specific antibodies

To detect ligand-induced phosphorylation downstream of hC5aR1, HEK293-T cells were transfected with 7 µg of hC5aR1 using the PEI transfection reagent. 48 h post-transfection, cells were starved for 6 h in serum-free DMEM. Following starvation, cells were stimulated with 1 µM of hC5a or hC5a^-d-Arg^ for 30 min at 37 °C. After stimulation, cells were scraped, collected, and harvested by centrifugation at 10,000 rpm for 10 min. The resulting pellet was dounce-homogenized in lysis buffer containing 20 mM HEPES, pH 7.4, 150 mM NaCl, 1x PhosSTOP (Roche, Cat. No. 57084100), and 1x protease inhibitor cocktail (Roche, Cat. No. 04693116001). Lysates were then solubilized in 1% (w/v) L-MNG at room temperature for 90 min and centrifuged at 15,000 rpm for 10 min. The cleared lysate was transferred to a separate tube containing pre-equilibrated M1-FLAG beads supplemented with 5 mM CaCl₂. Samples were tumbled at room temperature for 90 min to allow bead binding. The beads were then washed five times, alternating between low-salt buffer (20 mM HEPES, pH 7.4, 150 mM NaCl, 2 mM CaCl₂, and 0.01% L-MNG) and high-salt buffer (20 mM HEPES, pH 7.4, 350 mM NaCl, 2 mM CaCl₂, and 0.01% L-MNG). Bound proteins were eluted using FLAG-elution buffer containing 20 mM HEPES, pH 7.4, 150 mM NaCl, 2 mM EDTA, 0.01% L-MNG, and 250 µg/mL FLAG peptide. After elution, 5x reducing dye was added to the sample, and proteins were separated by SDS-PAGE for western blotting. The PVDF membrane was blocked with 5% BSA prepared in 1x TBST for 1 h at room temperature. The membrane was then incubated with primary antibodies (1:5000 dilution) specific for p324/327 and p332/334 for 1 h at room temperature. Following incubation, the membrane was washed three times with 1x TBST and incubated with an anti-rabbit HRP-conjugated secondary antibody for 1 h at room temperature. This was followed by three additional washes with 1x TBST. The signal was detected using the Promega ECL solution on a ChemiDoc imaging system (Bio-Rad). The membrane was subsequently stripped and re-probed for total receptor expression using an HRP-conjugated anti-FLAG M2 antibody (Sigma, 1:5000). The phosphorylation signal was quantified using ImageLab software (Bio-Rad) and normalized to the total receptor signal. Fold-normalized values were plotted using GraphPad Prism 10.3.1 software.

### Purification of h/mC5a, h/mC5a^-d-Arg^, Trx-hC5a and Trx-hC5a^-d-Arg^

C5a and C5a^-d-Arg^ from human and mouse were cloned into pET-32a(+) vector with an N-terminus 6x-His-Trx tag followed by TEV-site, and purified using a previously described protocol^50,82^, with slight modifications. Briefly, starter culture inoculated with freshly transformed *E. coli* SHuffle cells in 50 mL LB media supplemented with 100 μg/mL ampicillin was grown overnight at 30 °C. This was inoculated in 1 L LB media, similarly, supplemented with 100 μg/mL ampicillin, and the culture grown at 30 °C till O.D600 reached 0.8-1. Subsequently, the protein expression was induced with 1 mM IPTG and shifted to 16 °C for overnight induction. Post-harvesting, cells were incubated with 1 mg/mL lysozyme in 50 mM HEPES, pH 7.4, 300 mM NaCl, 30 mM Imidazole, 1 mM PMSF, and 2 mM benzamidine for 40 min at 4 °C followed by disruption with ultrasonication and removal of cell debris with high-speed centrifugation. Trx-C5a/C5a^-d-Arg^ in the lysate was captured on an Ni-IDA resin (Takara, Cat. No: 635662) using gravity flow columns. Resin containing bound protein was thoroughly washed with 50 mM HEPES, pH 7.4, 1 M NaCl, 30 mM Imidazole to remove non-specific proteins, and Trx-fused-C5a/C5a^-d-Arg^ was eluted with 50 mM HEPES, pH 7.4, 150 mM NaCl, 300 mM Imidazole. To remove imidazole from the eluted protein, it was dialysed overnight in 30 mM HEPES and 150 mM NaCl in 4 °C and stored in 10% glycerol at -80 °C for further use in structural determination. To prepare His-Trx-free C5a/C5a^-d-Arg^, Trx-His tag was cleaved by incubation for 16 h, at room temperature, with TEV-protease (1:20 w/w, TEV: fusion protein). Cleaved C5a/C5a^-d-Arg^, was further separated by cation-exchange chromatography and stored at -80 °C with 10% glycerol concentration. This was used for functional assays and complexing unless specified otherwise.

### Receptor purification from *Sf9* cells

Human and mouse C5aR2 was purified from *Sf9* cells following a previously described protocol^12^. Briefly, human and mouse C5aR2 expressing baculovirus were employed to set up infection in *Sf9* cells at a density of 2 x 10^6^ cells per mL and allowed to grow for 72 h at 27 °C followed by harvesting the cells by centrifugation at 5,000 rpm for 15 min. The harvested pellet was immediately flash frozen in liquid nitrogen and stored at -80 °C until further use.

For purification, the receptor-expressing pellets were thawed and sequentially homogenised in hypotonic buffer (20 mM HEPES, pH 7.4, 1 mM MgCl2, 2 mM KCl, 1 mM PMSF, 2 mM benzamidine) followed by hypertonic buffer (20 mM HEPES, pH 7.4, 1M NaCl, 1 mM MgCl2, 2 mM KCl, 1 mM PMSF, 2 mM benzamidine) and centrifuged at 20,000 rpm for 20 min at 4 °C to remove cytosolic contaminants. Membrane solubilisation was carried out by resuspending the pellet in solubilisation buffer (20 mM HEPES, pH 7.4, 450 mM NaCl, 1% L-MNG (Anatrace, Cat. no: NG310), 0.1% CHS (Sigma, Cat. no: C6512), 1 mM PMSF, 2 mM benzamidine) and incubating the lysate for 2 h at 4 °C in the presence of 2 mM iodoacetamide. Following solubilisation, the lysate was 3-fold diluted in dilution buffer (20 mM HEPES, pH 7.4, 1 mM PMSF, 2 mM benzamidine, 2.5 mM CaCl2) and subjected to high-speed centrifugation at 20,000 rpm for 20 min. The resulting supernatant was filtered through 0.45 µ bottle-top filters (Merck Millipore, Cat. No: HVLP04700) and loaded on to pre-equilibrated gravity flow columns containing M1 anti-Flag resin (prepared in-house). The unbound and non-specifically bound proteins were removed by three washes of low salt buffer (20 mM HEPES, pH 7.4, 150 mM NaCl, 2 mM CaCl2, 0.1% L-MNG, 0.01% CHS) alternated with two washes of high salt buffer (20 mM HEPES, pH 7.4, 2 mM CaCl2, 350 mM NaCl, 0.1% L-MNG). The receptor was eluted in elution buffer (20 mM HEPES, pH 7.4, 150 mM NaCl, 0.01% L-MNG, 2 mM EDTA, 250 µg/ml FLAG-peptide). Finally, free cysteine residues in the receptor were blocked with 2 mM iodoacetamide and excess iodoacetamide was quenched with 2 mM L-cysteine. The purified receptor was stored in 10% glycerol at -80 °C until further use.

For Apo-C5aR2, the receptor purification was carried out in the absence of any ligand, however for ligand-bound receptor (Trx-hC5a/Trx-hC5a^-d-Arg^-C5aR2, mC5a^-d-Arg^-mC5aR2, EP54-C5aR2, C5a^pep^-C5aR2, R8Y-C5aR2), either 100 nM Trx-hC5a/hC5a^-d-Arg^, mC5a^-d-Arg^ or 1 µM peptide ligands were added at each purification step while keeping 1 µM Trx-hC5a/hC5a^-d-Arg^ and mC5a^-d-Arg^ and 10 µM peptide ligands at elution.

### Complexing of Trx-hC5a/Trx-hC5a^-d-Arg^-bound C5aR2 and mC5a^-d-Arg^-mC5aR2

For the structure determination of Trx-hC5a/Trx-hC5a^-d-Arg^-bound C5aR2 and mC5a^-d-Arg^-bound mC5aR2, SEC of the purified ligand-bound receptor was performed with running buffer having 20 mM HEPES, pH 7.4, 150 mM NaCl, 0.01% L-MNG, 0.001% CHS, supplemented with 100 nM Trx-hC5a/Trx-hC5a^-d-Arg^/mC5a^-d-Arg^, as applicable. The Trx-hC5a/Trx-hC5a^-d-Arg^-hC5aR2 and mC5a^-d-Arg^-mC5aR2 were separated on Superose^TM^ 6 Increase 10/300 GL (Cytiva, Cat. No: 29091596) and Superdex^TM^ 200 Increase 10/300 GL (Cytiva, Cat. No: 28990944), respectively. The elution fractions corresponding to Trx-hC5a/hC5a^-d-Arg^-bound C5aR2 were pooled and concentrated to 6-12 mg/mL using a 100 MWCO concentrator (Cytiva, Cat. no: 28932319) for Cryo-EM grid preparation.

### Cryo-EM grid preparation and data collection

3 μl of the purified Trx-hC5a/hC5a^-d-Arg^-hC5aR2, mC5a^-d-Arg^-mC5aR2, and Fab4C8-C5aR2 complexes were applied onto glow discharged Quantifoil holey carbon grids (R1.2/1.3, Au, 300 mesh) at a concentration ranging from 5-20 mg/ml. The grids were blotted for 4 s at 4 °C and 100% humidity with a blot force of 10 using a Vitrobot Mark IV (Thermo Fischer Scientific) and immediately plunge frozen in liquid ethane (-181 °C).

Data collection of all samples was performed on a Titan Krios G3i/G4 (Thermo Fisher Scientific) operating at an accelerating voltage of 300 kV equipped with a Gatan K3 direct electron detector and BioQuantum K3 imaging filter. Movie stacks were acquired in counting mode at a pixel size of 0.83 Å/px and a dosage rate of approximately 7.7 - 8.6 e^-^/Å^2^/s using EPU software over a defocus range of -0.8 to -1.6 μm. Each movie was fractionated into 64 frames with a total dose of 62.1-64.3 e^-^/Å^2^ that was obtained throughout the 5.0-6.0 s exposure period. In total 4,139, 3,200, 3,350, 3,381, 3,576, 4,189 and 1,652 movie stacks were acquired for Trx-C5a-C5aR2, Trx-C5a^d-Arg^-C5aR2, mC5a^-d-Arg^-mC5aR2, Fab4C8-Apo-C5aR2, Fab4C8-C5a^pep^-C5aR2, Fab4C8-EP54-C5aR2 and Fab4C8-R8Y-C5aR2, respectively.

### Image processing and map construction

All datasets were processed following a similar pipeline in sub-programs of CryoSPARC^72^ v4.6, unless stated otherwise. Briefly, dose-fractionated movie stacks were aligned with Patch Motion Correction (multi), and contrast transfer function (CTF) parameters were estimated using Patch CTF (multi). Particles were auto-picked using the blob-picker module, extracted using a box-size of 320 px (Fourier cropped to 64 px) and cleaned using reference-free 2D classification to remove ice contamination and distorted particles. Selected particles were subjected to reference free *ab-initio* reconstruction into multiple classes followed by heterogenous refinement. The particles corresponding to the best class were re-extracted using a box size of 320 px (Fourier cropped to 256 px) and subjected to non-uniform (NU) refinement and local refinement providing reconstructions with resolutions ranging from 2.97 Å – 3.82 Å at Fourier Shell Correlation of 0.143. Local resolutions were estimated using the LocRes module in cryoSPARC, providing half-maps as input. Details and number of particles in each step in the processing pipelines of individual complexes are presented in Figure S4-S5.

### Model building and refinement

The coordinates of C5aR2 obtained from Alpha Fold^95^ (AF-Q9P296) were used to dock into the EM maps of Trx-C5a-C5aR2, Fab4C8-Apo-C5aR2, Fab4C8-C5a^pep^-C5aR2, Fab4C8-EP54-C5aR2 and Fab4C8-R8Y-C5aR2 using UCSF Chimera^96^. The resulting coordinates and maps were imported and subjected to “All-atom Refine” in COOT^97–99^, followed by iterative rounds of manual adjustment in COOT and real-space refinement in Phenix^100,101^. The final refined models displayed good geometry, and the refinement statistics for all models are presented in Table S2. Since the Fab4C8 was derived from a hybridoma clone and we did not have the sequence information for the Fab component, we were unable to build models for the corresponding Fab densities in the map.

Protein-protein contacts and volumes of pockets were calculated using the PDBsum server^102^. All figures were prepared using UCSF Chimera or UCSF ChimeraX^96,103^.

### BM2020-7 and BM2020-8 peptide synthesis

Peptide synthesis was performed manually using Fmoc (9-fluorenylmethyloxycarbonyl)- based solid phase peptide synthesis (SPPS) on 2-chlorotrityl chloride (2-CTC) and Rink Amide AM resins. A ratio of 4 eq. of amino acid, 4 eq. of HBTU and 8 eq. of DIPEA was used for each coupling. N-terminal acetylation was performed on-resin using 2 x 5 min treatments of 5% acetic anhydride (Sigma-Aldrich) and 3% DIPEA (Sigma-Aldrich) in DMF (25° C) Following synthesis, the peptide was cleaved from the resin and side chain protecting groups removed with a treatment of trifluoroacetic acid (TFA)/triisopropylsilane(TIPS)/water (95:2.5:2.5). Following cleavage, purification was performed using reverse phase high performance liquid chromatography (RP-HPLC) using an increasing gradient of 1% buffer B (90% acetonitrile,0.05% TFA) in buffer A (0.05% TFA) over an 80 min period (Phenomenex Jupiter 300 Å, 10 µm, 250 x 21.2 mM). Analysis was performed using electrospray mass spectrometry (ESI-MS) (AB SCIEX API 2000) to identify fractions containing mass/charge ratios that matched the desired product. Purity was determined using analytical RP-HPLC (Agilent, 300 Å, 5 µm, 150 x 2.1 mM) with all peptides purified until >95% purity. C5a was synthesised as previously described^91^.

### mAb4C8 production and Fab4C8 generation

C5aR2-specific monoclonal antibodies producing hybridoma clones (mAb4C8) were obtained as a generous gift from Monash University. The frozen clones were revived in Dulbecco’s Modified Eagle Medium (DMEM) supplemented with 10% FBS and cultured at 37 °C in a 5% humidified CO2 incubator. Once the cells reached optimal density, approximately, 40 million cells were seeded in CELLine bioreactor (DWK Life Sciences; Cat. No: WCL-1000) using Hybridoma SFM (serum free medium) (Gibco, Cat. No: 12045-076) to facilitate high-yield antibody production. The culture was maintained at 37 °C in a 5% CO₂ humidified incubator, with supernatant was harvested weekly for antibody collection. Fresh Hybridoma SFM was replenished at each harvest to sustain continuous cell proliferation and monoclonal antibody secretion. The supernatant from each harvest is flash frozen in liquid nitrogen and stored at -80 °C till further use.

For antibody purification, the harvested supernatant was thawed and subjected to centrifugation at 22,000 x g for 20 min to remove residual cellular debris and sequentially filtered through 0.45 µ pre-filters followed by 0.22 µ filters to ensure the removal of particulates. To facilitate mAb binding to protein A affinity resin (MabSelect^TM^, Cytiva Cat. No: 17519902), the supernatant was buffered with 2 M Na2HPO4, pH 8.0, prior to loading on pre-equilibrated gravity flow columns containing ProteinA beads. The loading was carried out at a constant flow rate of 0.5 mL/min to maximize antibody recovery, following this the column was washed with 1x PBS, pH 7.4 (self-prepared) to remove unbound contaminants. The presence of antibody in wash fractions was monitored by measuring absorbance at 280 nm (A280) for IgG using NanoDrop (Thermo Fisher Scientific) and carried out till the absorbance reached a minimal baseline value. Subsequently, the column was washed with 100 mM NaCl to displace loosely bound non-specific proteins followed by washing with 1x PBS. The bound mAb4C8 was eluted under low pH buffer condition using 100 mM NaH2PO4, pH 2.5, and eluate was immediately neutralised with 1M Na2HPO4, pH 8.0. The elution was carried out till A280 for IgG minimises to ≤0.01, indicating the complete antibody recovery. The purified mAb4C8 was dialysed against buffer containing 20 mM HEPES, pH 7.4, 150 mM NaCl for 16 h and flash frozen in liquid nitrogen and stored at -80 °C until further use.

To generate antigen-binding fragments (Fab), purified mAb4C8 was digested using papain. 10 mg mAb4C8 was concentrated to 20- to 25-fold using a Cytiva Vivaspin 100 kDa MWCO centrifugal device (Cat. No: 28932363). Papain digestion was performed using a crude papain extract derived from *Carica papaya* (Sigma, Cat. No: P375-25G). The extract was solubilised in papain digestion buffer (20 mM HEPES, pH 7.4, 150 mM NaCl, 10 mM EDTA, 50 mM L-cysteine) and vortexed until completely dissolved. Insoluble debris was removed by centrifugation at 14,000 rpm for 10 min and the resulting supernatant was used as the enzyme source. The digestion was set up by incubating 10 mg of mAb4C8 with papain at final enzyme concentration of 50 µg/mg of mAb at 37 °C for 3 h. The reaction was irreversibly quenched with 50 mM iodoacetamide.

To separate the Fab and Fc fragments, the reaction mixture was loaded onto a pre-equilibrated HiLoad^TM^ 16/600 Superdex™ 200 pg (Cytiva, Cat. No: 28989335) for SEC. The purified Fab fragment was eluted in a buffer containing 20 mM HEPES, pH 7.4 and 150 mM NaCl, pooled, flash-frozen in liquid nitrogen and stored in 10% glycerol at -80 °C for further experiments.

### Complexing of Fab4C8-bound Apo/C5a^pep^/EP54/R8Y-C5aR2

Purified C5aR2 was mixed with 2-fold molar excess SEC purified Fab4C8 with or without either C5a^pep^ or EP54 or R8Y at a final concentration of 10 µM to form Fab4C8-bound Apo and peptide-bound C5aR2 complexes, respectively. The reaction mix was allowed for complexing in constant tumbling conditions at room temperature. The complex was separated by injecting the reaction mix in a Superose™ 6 Increase 10/300GL column (Cytiva, Cat. No: 29091596), equilibrated with SEC buffer (20 mM HEPES, pH 7.4, 150 mM NaCl, 0.01% L-MNG, 0.001% CHS, 1 µM either EP54/C5a^pep^/R8Y) and the peak fractions were pooled together, supplemented with either EP54/C5a^pep^/R8Y at a final concentration of 10 µM and concentrated to 5-20 mg/mL concentration in a 100 kDa centrifugal device.

### Ib30 reactivity

Agonist induced Ib30 reactivity downstream to hC5aR2 and mC5aR2 was measured by following the same protocol as described for NanoBiT-based ꞵarr1/2 recruitment and as discussed previously^104,105^. HEK-293T cells were transiently transfected with 5 µg of Ib30 tagged with N-terminal LgBiT fragment, 2 µg of ꞵarr1 tagged with C-terminal SmBiT fragment and untagged 3 µg of either hC5aR2 or mC5aR2 cloned in pcDNA3.1.

### Quantification and statistical analysis

GraphPad Prism 10.3.1 software was used to plot and analyze all the functional data presented in this manuscript, and all the relevant details such as number of replicates, data normalization, mean ± SEM, and statistical analyses are mentioned in the corresponding figure legends.

## Notes

### Competing Interest Statement

The authors have declared no competing interest.

