## Supplemental Figures for "Molecular mechanism of naturally-encoded signaling-bias at the complement anaphylatoxin receptors"

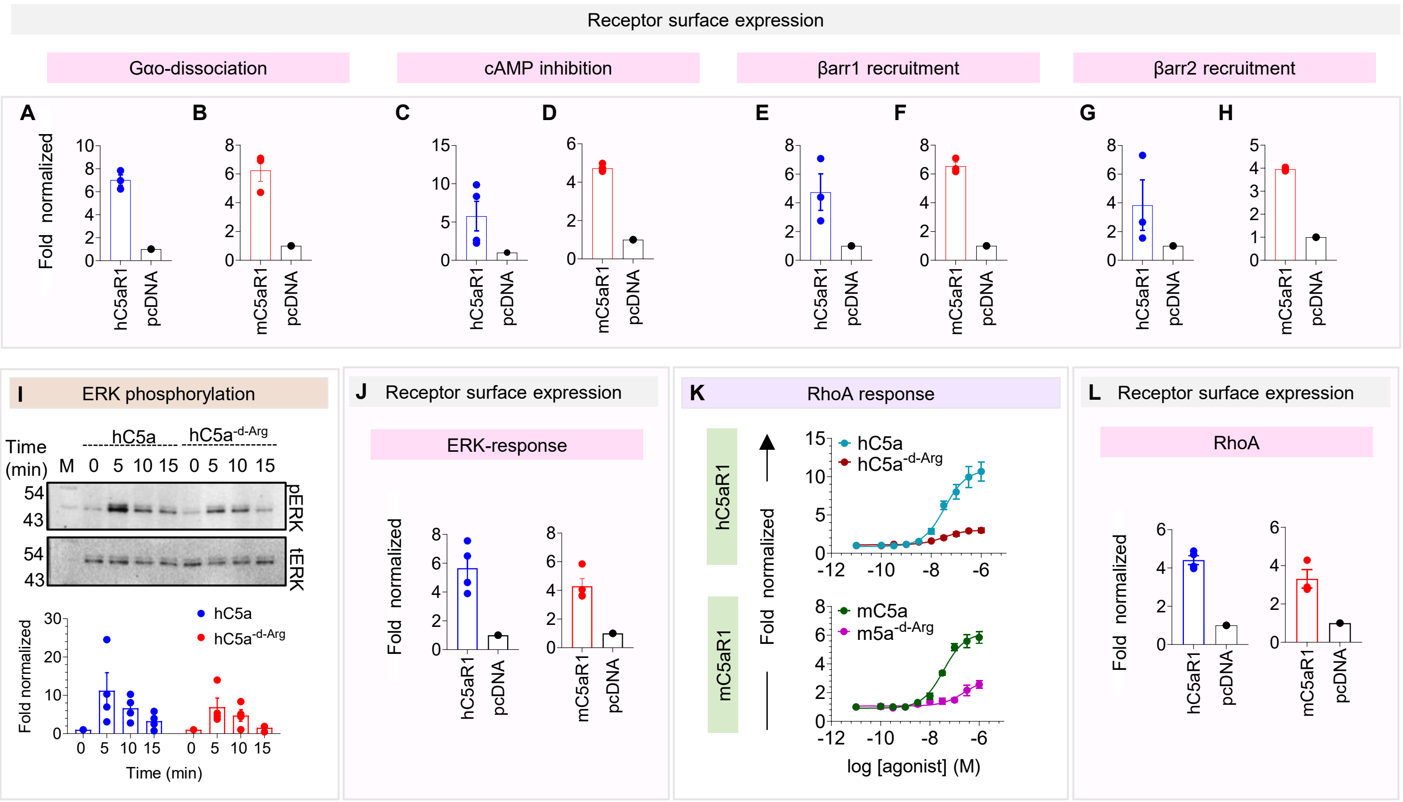


**Figure S1. Receptor surface expression, ERK1/2 phosphorylation and RhoA response assays, related to figure 1.**

**(A-H)** Receptor surface expression for different assays (corresponding to **Figure 1B-C**), measured by whole cell-based surface ELISA. Receptor fold-expression was calculated by normalising ELISA and Janus signals of receptor-transfected cells with respect to that of signals recorded for mock-transfected (pcDNA) cells. Data represents mean ± SEM, n=3-4 independent experiments.

**(I)** ERK1/2 phosphorylation using western blot to measure its activation downstream to hC5aR1 in response to hC5a and hC5a$^{d\text{-}Arg}$. Stimulation with hC5a and hC5a$^{d\text{-}Arg}$ leads to robust ERK1/2 phosphorylation in HEK293-T cells expressing hC5aR1. The upper panel shows representative blots, and the lower panel provides densitometry-based quantification. Data are presented as mean ± SEM (n = 3), normalized to the 0-min time point without ligand stimulation, taken as 1.

**(J)** Surface expression of indicated receptors measured using whole-cell based surface ELISA technique. Data is presented as mean ± SEM (n = 4), normalized as fold change over mock transfection (pcDNA) in various assays (corresponding to **Figure 1F -G**).

**(K)** Luminescence-based SRF-RE reporter assay used to measure RhoA response downstream of human and mouse C5aR1 in response to dose-dependent concentrations of the indicated ligands. Data are presented as mean ± SEM (n = 4), normalized with respect to the lowest ligand concentration, taken as 1.

**(L)** Surface expression of indicated receptors measured using whole-cell based surface ELISA technique. Data is presented as mean ± SEM (n = 4), normalized as fold change over mock transfection (pcDNA) in various assays (corresponding to **Figure Figure S1K**).

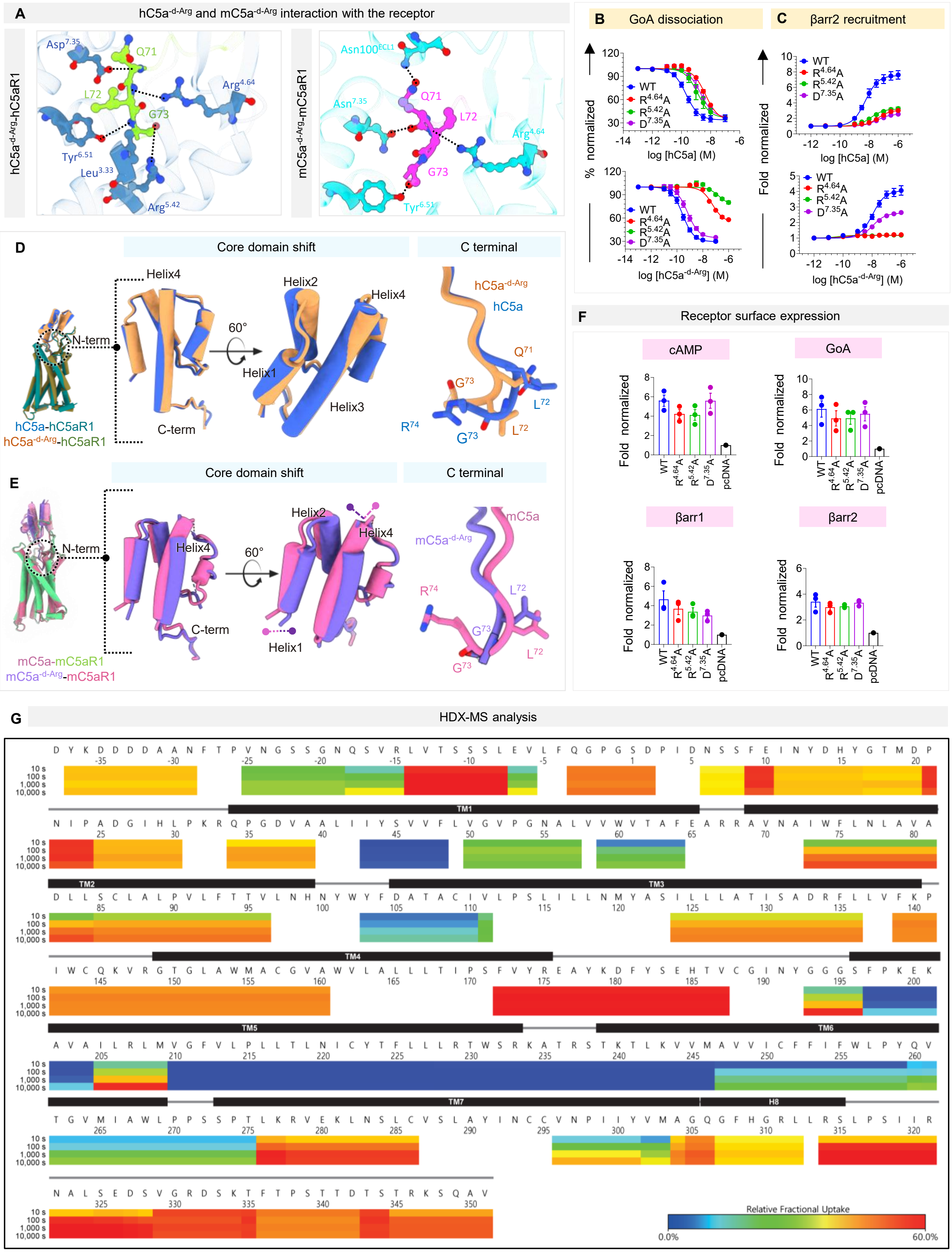

A
hC5a-d-Arg and mC5a-d-Arg interaction with the receptor
hC5a-d-Arg-hC5aR1
mC5a-d-Arg-mC5aR1
B
GoA dissociation
C
βarr2 recruitment
% normalized
Fold normalized
log [hC5a] (M)
log [hC5a-d-Arg] (M)
WT
R4.64A
R5.42A
D7.35A
D
Core domain shift
C terminal
Helix1
Helix2
Helix3
Helix4
N-term
C-term
60°
hC5a-hC5aR1
hC5a-d-Arg-hC5aR1
E
Core domain shift
C terminal
mC5a-mC5aR1
mC5a-d-Arg-mC5aR1
F
Receptor surface expression
cAMP
GoA
βarr1
βarr2
Fold normalized
WT
R4.64A
R5.42A
D7.35A
pcDNA
G
HDX-MS analysis
10 s
100 s
1,000 s
10,000 s
TM1
TM2
TM3
TM4
TM5
TM6
TM7
H8
Relative Fractional Uptake
0.0%
60.0%

**Figure S2. Structural and functional insights into C5a$^{-d-Arg}$ mediated G-protein bias, related to figure 2.**

**(A)** Interaction of G73 residue within the orthosteric binding pocket of hC5a$^{-d-Arg}$-hC5aR1 and mC5a$^{-d-Arg}$-mC5aR1.

**(B-C)** GoA-dissociation **(B)** and βarr2 recruitment **(C)** downstream to hC5aR1 mutants in response to hC5a and hC5a$^{-d-Arg}$ orthosteric pocket residues, related to **Figure 2B-C**.

**(D-E)** Superimposition of hC5a/hC5a$^{-d-Arg}$-bound hC5aR1 and mC5a/mC5a$^{-d-Arg}$-mC5aR1 reveals a core domain shift in the ligands, while both the ligands maintain a similar hook like orientation.

**(F)** Receptor surface expression for different assays (corresponding to **Figure 2B-C and Figure S2B**), measured by whole cell-based surface ELISA. Receptor fold-expression was calculated by normalising ELISA and Janus signals of receptor-transfected cells with respect to that of signals recorded for mock-transfected (pcDNA) cells. Data represents mean ± SEM, n=3-4 independent experiments.

**(G)** Relative fractional uptake of deuterium performed for Apo-mC5aR1 (without ligand) shown in heat map format. The plot indicating overall HDX levels are higher in the N-term, C-term, loop regions than TMs. The HDX levels at ICL3 are very low while the HDX levels of TM7 are relatively high considering it is a transmembrane region. TMs and loop regions are indicated according to the snake map of GPCRdb (GPCRdb.org).

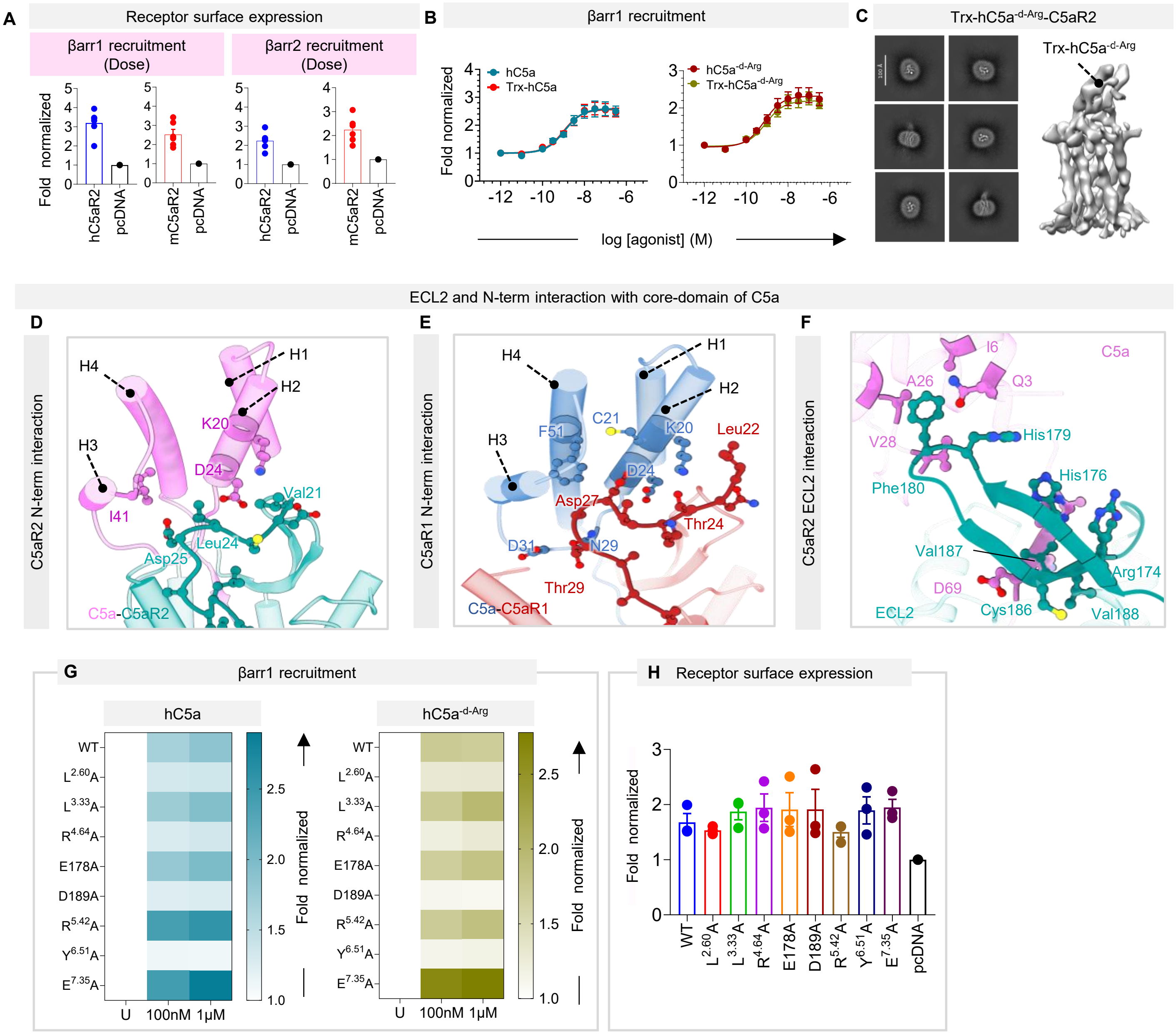
A
Receptor surface expression
βarr1 recruitment (Dose)
βarr2 recruitment (Dose)
Fold normalized
hC5aR2
pcDNA
mC5aR2
B
βarr1 recruitment
hC5a
Trx-hC5a
hC5a-d-Arg
Trx-hC5a-d-Arg
log [agonist] (M)
C
Trx-hC5a-d-Arg-C5aR2
Trx-hC5a-d-Arg
ECL2 and N-term interaction with core-domain of C5a
D
C5aR2 N-term interaction
H1
H2
H3
H4
K20
D24
I41
Val21
Leu24
Asp25
C5a-C5aR2
E
C5aR1 N-term interaction
F51
C21
K20
D24
Leu22
Asp27
Thr24
D31
N29
Thr29
C5a-C5aR1
F
C5aR2 ECL2 interaction
I6
C5a
A26
Q3
V28
His179
Phe180
His176
Val187
Arg174
D69
Cys186
Val188
ECL2
G
βarr1 recruitment
hC5a
hC5a-d-Arg
WT
L2.60A
L3.33A
R4.64A
E178A
D189A
R5.42A
Y6.51A
E7.35A
U
100nM
1μM
Fold normalized
H
Receptor surface expression
Fold normalized
pcDNA

**Figure S3. Structural insights and functional validation of C5a and C5aR2 binding, related to figure 3.**

**(A)** Receptor surface expression for βarr1/2 recruitment (corresponding to **Figure 3B**), measured by whole cell-based surface ELISA. Receptor fold-expression was calculated by normalising ELISA and Janus signals of receptor-transfected cells with respect to that of signals recorded for mock-transfected (pcDNA) cells. Data represents mean ± SEM, n=6 independent experiments.

**(B)** Dose-response curve for βarr1 recruitment downstream to C5aR2 in response to hC5a/hC5a$^{-d-Arg}$and Trx-hC5a/hC5a$^{-d-Arg}$, measured by NanoBiT-based assay. Data represents mean ± SEM, n=3 for independent experiments.

**(C)** 2-D class averages and 3-D reconstruction of Trx-hC5a$^{-d-Arg}$ bound to C5aR2.

**(D-E)** Structural snapshots displaying non-bonded contacts and polar interactions between several residues from the N-terminus of C5aR2 (**D**) and C5aR1 (**E**) with the core helices of C5a (related to **Figure 3E**).

**(F)** Structural snapshot showing ECL2 of C5aR2 engaged in polar and non-bonded contacts with C5a in C5a-bound C5aR2 structure (related to **Figure 3E**).

**(G)** Heat map showing βarr1 recruitment downstream to C5aR2 mutants rationally designed on the basis of C5a and C5aR2 interaction analysis. The recruitment was performed by undertaking NanoBiT-based assay. Data represents mean values of three independent experiments.

**(H)** Receptor surface expression for βarr1 recruitment downstream to C5aR2 mutants (corresponding to **Figure S8G**), measured by whole cell-based surface ELISA. Receptor fold-expression was calculated by normalising ELISA and Janus signals of receptor-transfected cells with respect to that of signals recorded for mock-transfected (pcDNA) cells. Data represents mean ± SEM, n=3 independent experiments.

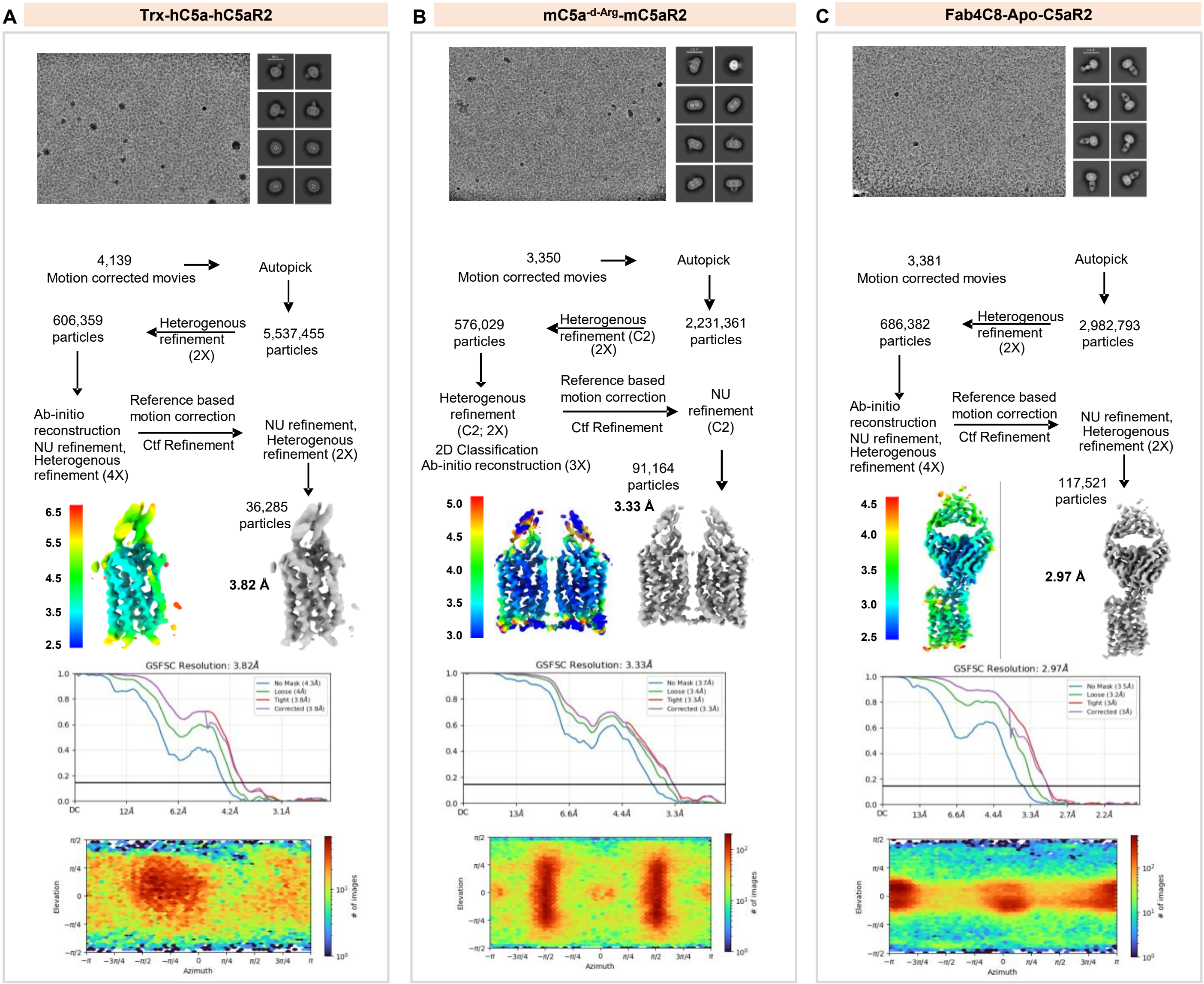


**Figure S4: Workflow for cryo-EM data processing of C5aR2 complexes, related to figure 3, 4 and 5.**

**(A-B)** Representative cryo-EM micrograph, selected 2D class averages representing different orientations, schematic representation of cryo-EM data processing workflow, local resolution map of the 3D reconstruction, gold standard fourier shell correlation curve (GSFSC) at 0.143 threshold, and angular distribution of the particles against the final reconstruction of **(A)** Trx-hC5a-hC5aR2, **(B)** mC5a$^{-d-Arg}$-mC5aR2 complexes, and (**C**) Fab4C8-Apo-hC5aR2.

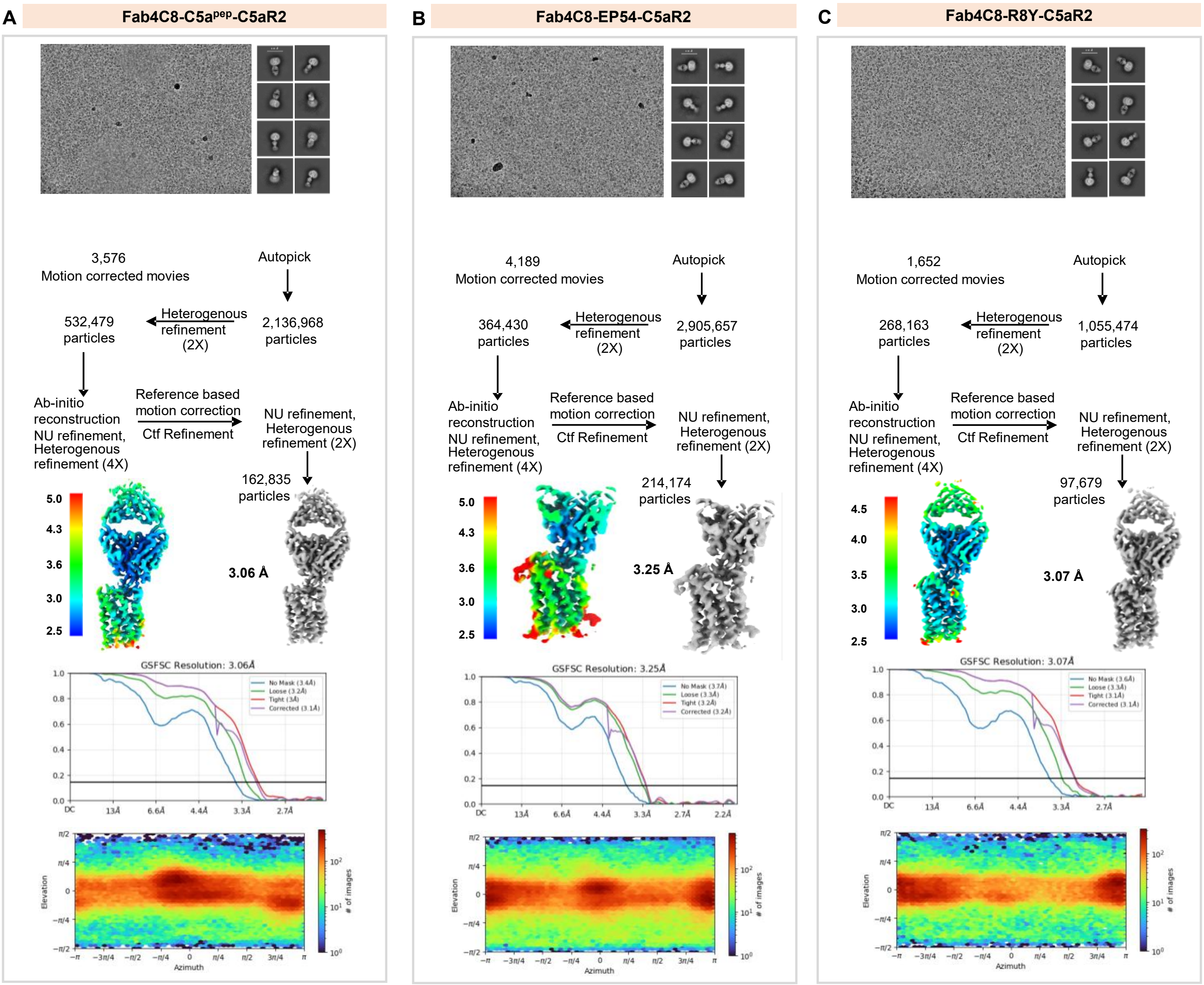


**Figure S5: Workflow for cryo-EM data processing of C5aR2 complexes, related to figure 5.**

**(A-C)** Representative cryo-EM micrograph, selected 2D class averages representing different orientations, schematic representation of cryo-EM data processing workflow, local resolution map of the 3D reconstruction, gold standard fourier shell correlation curve (GSFSC) at 0.143 threshold, and angular distribution of the particles against the final reconstruction of (**A**) Fab4C8-C5a$^{pep}$-hC5aR2, (**B**) Fab4C8-EP54-C5aR2 and (**C**) Fab4C8-R8Y-C5aR2complexes.

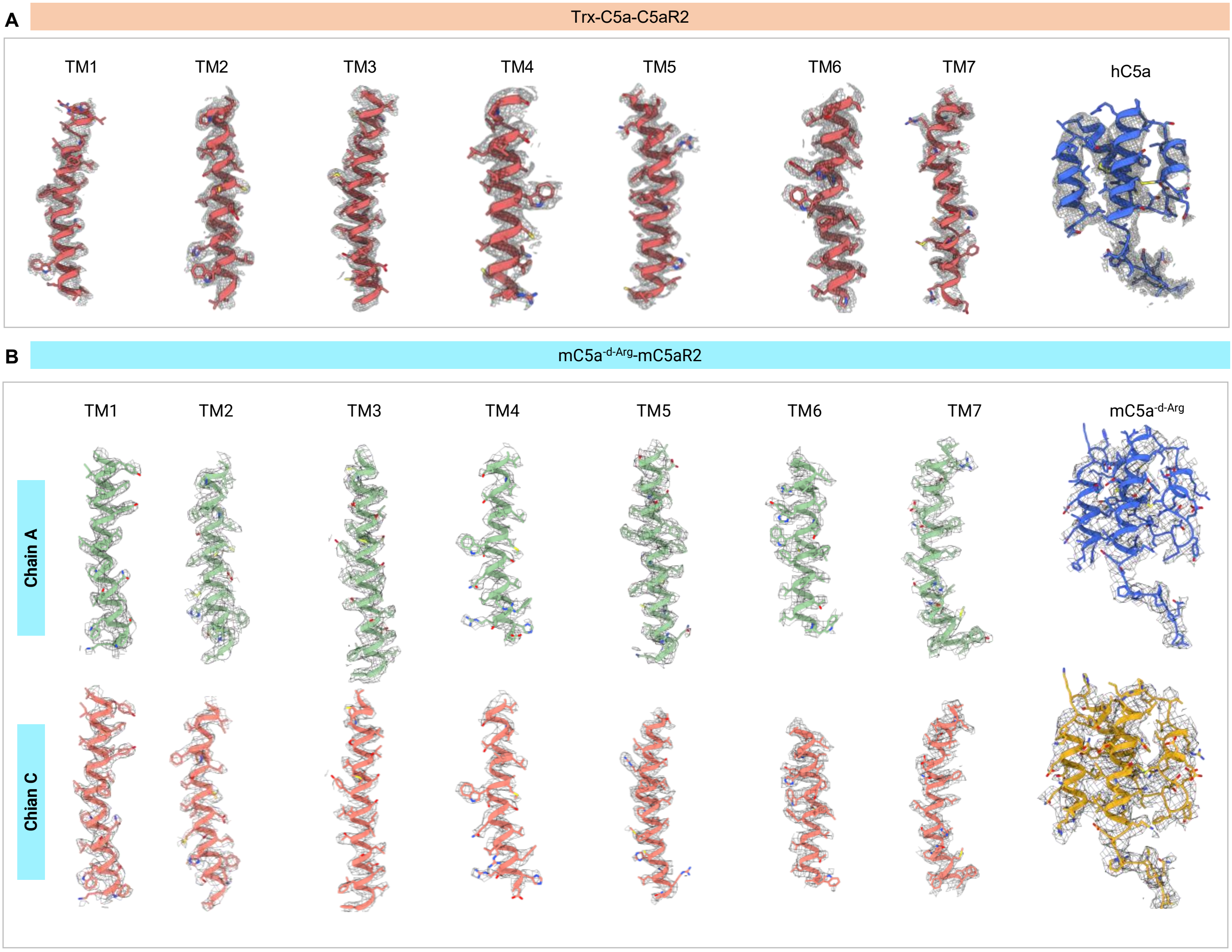


**Figure S6. Representative electron density maps, related to figure 3 and 4.**

**(A-B)**. EM densities of TM1 to TM7 and ligands of Trx-C5a-C5aR2 (**A**) and mC5a$^{-d\text{-}Arg}$-mC5aR2 (**B**) complexes, respectively.

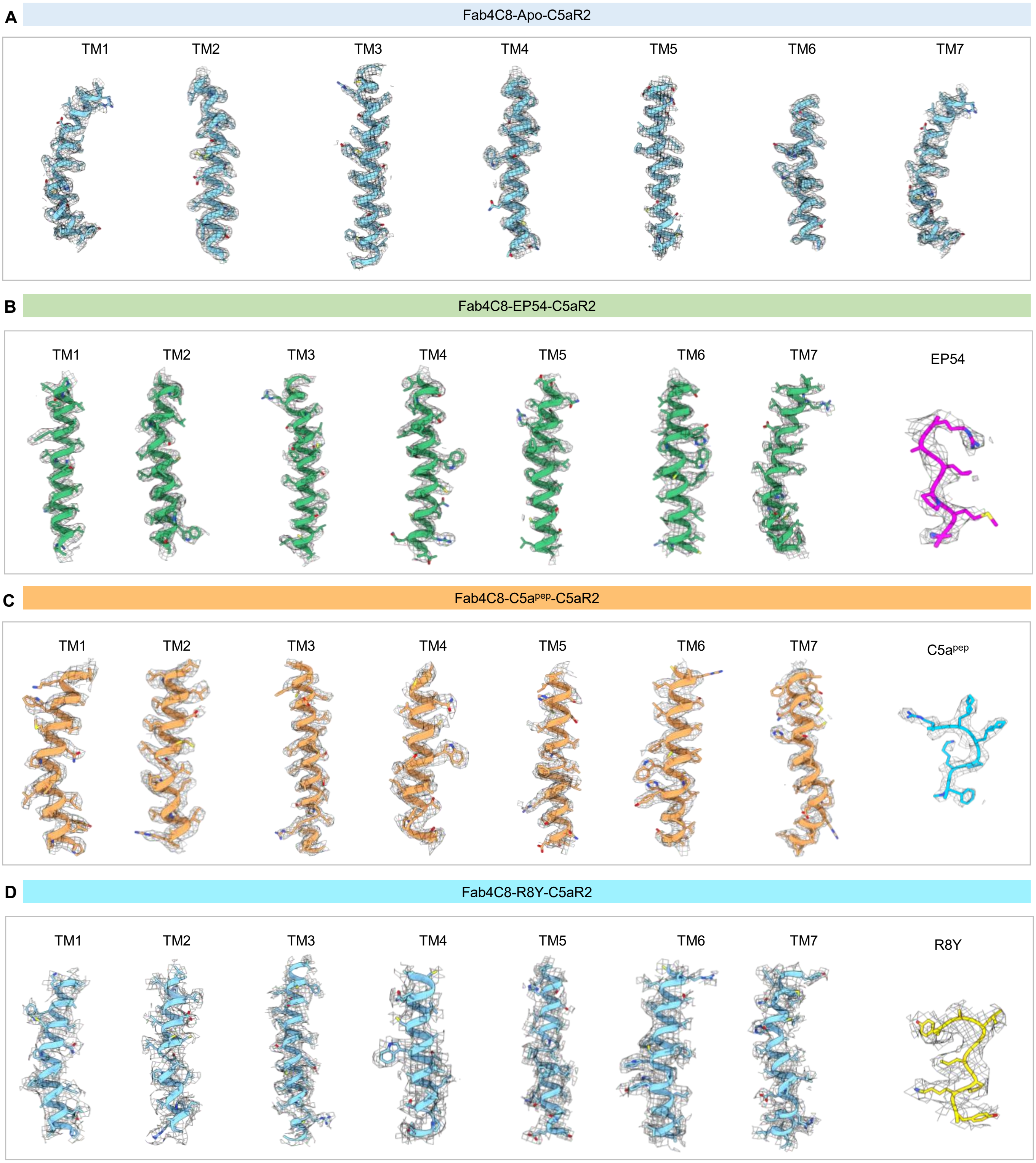


**Figure S7. Representative electron density maps, related to figure 5 .**

**(A-D)**. EM densities of TM1 to TM7 and ligands of Fab4C8-Apo-C5aR2, Fab4C8-EP54-C5aR2, Fab4C8-C5a$^{pep}$-C5aR2, and Fab4C8-R8Y-C5aR2 complexes, respectively.

| | Trx-hC5a-C5aR2 | | mC5a$^{-d-Arg}$-mC5aR2 | | Apo-C5aR2 | | EP54-C5aR2 | | C5a$^{pep}$-C5aR2 | | R8Y-C5aR2 | |
|---|---|---|---|---|---|---|---|---|---|---|---|---|
| **Component** | **Total residues** | **Resolved residues** | **Total residues** | **Resolved residues** | **Total residues** | **Resolved residues** | **Total residues** | **Resolved residues** | **Total residues** | **Resolved residues** | **Total residues** | **Resolved residues** |
| **Ligand** | T1-R74 | T1-R74 | N1-G76 | H3-G76 | - | - | Y1-R10 | P5-R10 | MEA1-DAR6 | MEA1-DAR6 | Y1-Y8 | Y1-Y8 |
| **Receptor** | Met1 - Val337 | Val21 - Ala306 | Met1-Val344 | Asp38-His314 | Met1-Val337 | Val36-Ala301 | Met1-Val337 | Ala37-Cys225<br>Cys231-Phe294 | Met1-Val337 | Ala37-Cys225<br>Arg230-Phe294 | Met1-Val337 | Val36-Ala140<br>Arg147-Cys225<br>Arg230-Phe294 |

**Figure S8. Residues resolved in Trx-hC5a-C5aR2, mC5a$^{-d-Arg}$-mC5aR2, Apo-C5aR2, EP54-C5aR2, C5a$^{pep}$-C5aR2 and R8Y-C5aR2.**

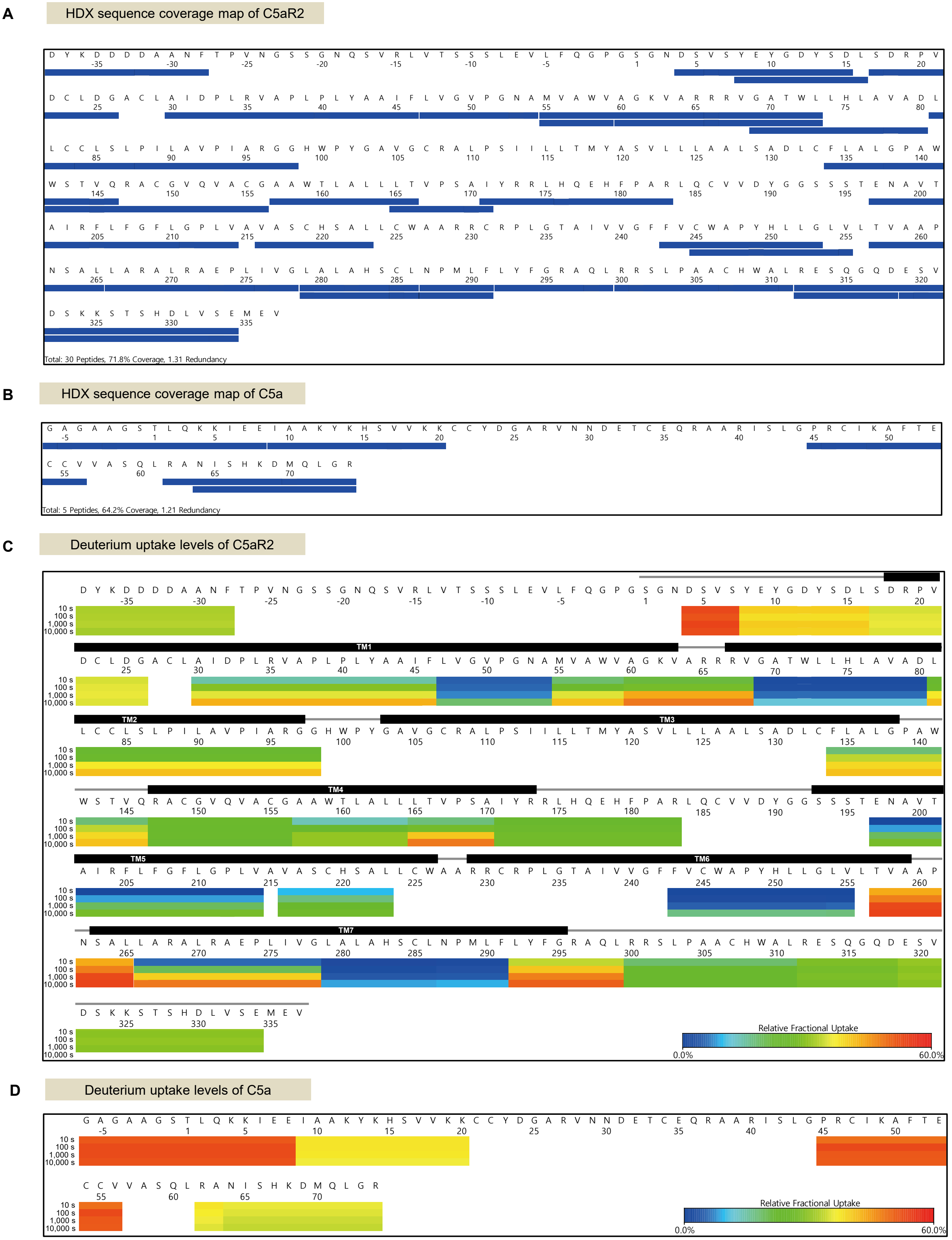


**Figure S9. HDX-MS analysis of C5a-C5aR2, related to Figure 3J**

**(A-B)** Sequence coverage map of C5aR2 **(A)** and C5a **(B)** that has undergone HDX-exchange. The blue bars indicate analyzed peptic peptides. The letters and numbers indicate the amino-acid sequence and their corresponding numbers, respectively. (Sequence number in negative indicates the tags and additional sequences present in the construct to facilitate receptor expression and purification).

**(C-D)** The deuterium uptake levels of Apo-C5aR2 **(C)** and C5a-alone **(D)** are visualized as heat maps. TMs and loop regions are indicated according to the snake map of GPCRdb (GPCRdb.org).

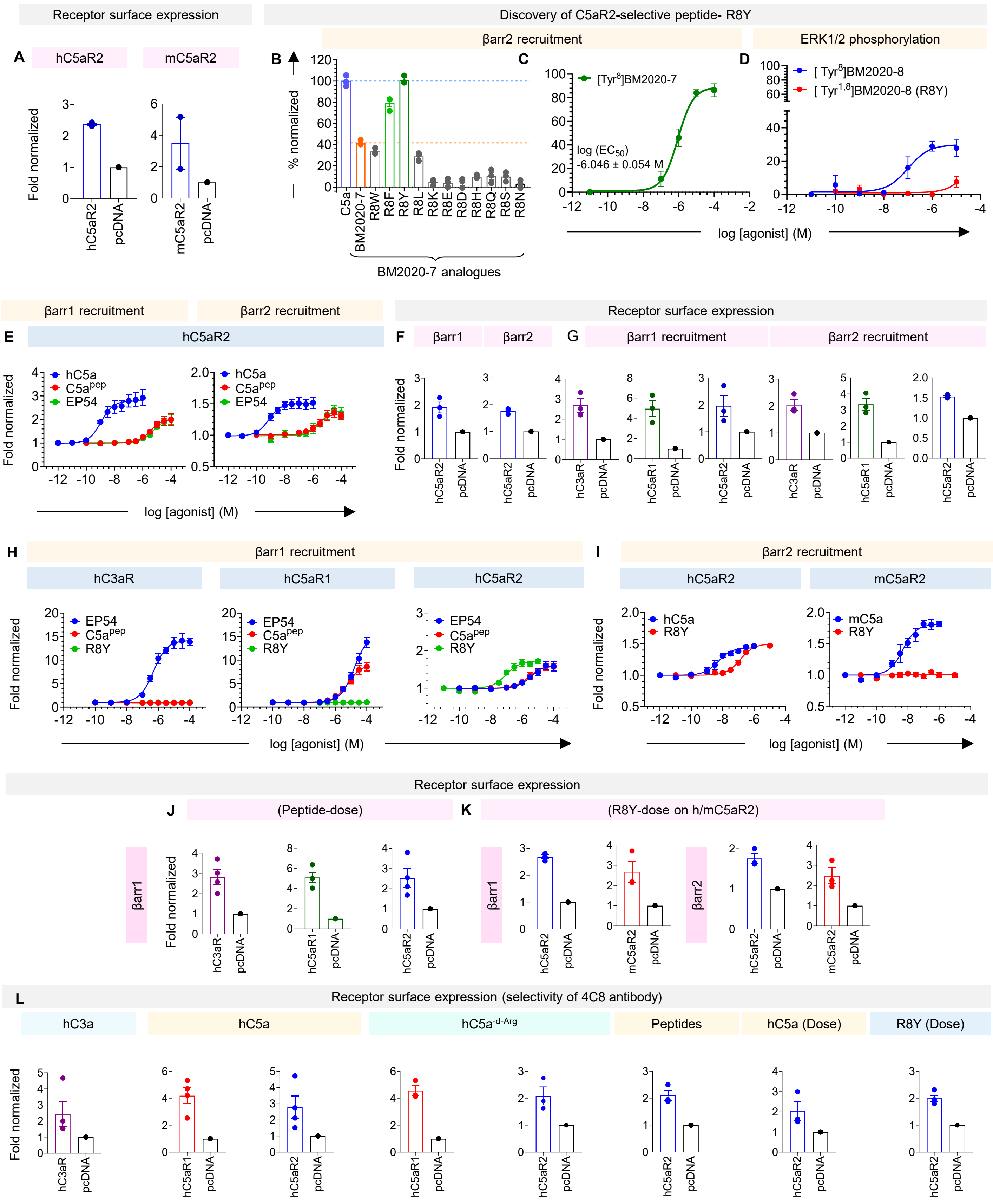
Receptor surface expression
Discovery of C5aR2-selective peptide- R8Y
A
hC5aR2
mC5aR2
Fold normalized
hC5aR2
pcDNA
mC5aR2
pcDNA
B
βarr2 recruitment
ERK1/2 phosphorylation
% normalized
C5a
BM2020-7
R8W
R8F
R8Y
R8L
R8K
R8E
R8D
R8H
R8Q
R8S
R8N
BM2020-7 analogues
C
[Tyr8]BM2020-7
log (EC50)
-6.046 ± 0.054 M
D
[ Tyr8]BM2020-8
[ Tyr1,8]BM2020-8 (R8Y)
log [agonist] (M)
βarr1 recruitment
βarr2 recruitment
Receptor surface expression
E
hC5aR2
hC5a
C5apep
EP54
F
βarr1
βarr2
G
βarr1 recruitment
βarr2 recruitment
hC3aR
hC5aR1
H
βarr1 recruitment
hC3aR
hC5aR1
hC5aR2
EP54
C5apep
R8Y
I
βarr2 recruitment
hC5aR2
mC5aR2
hC5a
mC5a
R8Y
Receptor surface expression
J
(Peptide-dose)
K
(R8Y-dose on h/mC5aR2)
βarr1
βarr2
mC5aR2
L
Receptor surface expression (selectivity of 4C8 antibody)
hC3a
hC5a
hC5a-d-Arg
Peptides
hC5a (Dose)
R8Y (Dose)

**Figure S10. Development of C5aR2-selective peptide agonist and its functional characterisation, related to figure 5.**

**(A)** Receptor surface expression for βarr1 recruitment (corresponding to **Figure 5B**), measured by whole cell-based surface ELISA. Receptor fold-expression was calculated by normalising ELISA and Janus signals of receptor-transfected cells with respect to that of signals recorded for mock-transfected (pcDNA) cells. Data represents mean ± SEM, n=2 independent experiments.

**(B-D)** HEK-293 cells were transiently transfected using C5aR2-Venus and β-arrestin 2-Rluc8 BRET pairs for 24 hours and seeded (100,000/well) overnight. Filtered light emissions between 460-490 nm (Rluc8) and 520-550 nm (Venus) were continually monitored for 90 min with ligand or vehicle added at the 0 min time point. **(B)** Ligand-induced BRET ratios (normalized to 100 nM C5a) of BM2020-7 position 8 analogues (100 µM). **(C)** Concentration-response curve for [$Tyr^{8}$]-BM2020-7. **(D)** Selectivity testing of [$Tyr^{8}$]BM2020-8 and [$Tyr^{1,8}$]BM2020-8 via C5aR1-mediated ERK signalling in stably expressing C5aR1-CHO cells. All data represent the mean ± SEM of triplicate measurements from 3-5 independent experiments.

**(E)** Dose-response curve for βarr1/2 recruitment downstream to hC5aR2 to investigate pharmacology of EP54 and $C5a^{pep}$ with respect to hC5a.

**(F-G)** Receptor surface expression for βarr1/2 recruitment downstream to indicated receptors (**F**- corresponding to **Figure S11E, G-**corresponding to **Figure 5D**). Surface expression was measured by whole cell-based surface ELISA. Receptor fold-expression was calculated by normalising ELISA and Janus signals of receptor-transfected cells with respect to that of signals recorded for mock-transfected (pcDNA) cells. Data represents mean ± SEM, n>3 independent experiments.

**(H-I)** Dose-response curve for βarr1/2 recruitment downstream to indicated receptors to display the selectivity profile of different peptide agonists. NanoBiT-based assay was employed in HEK293-T cells transiently transfected with either hC3aR or hC5aR1 or hC5aR2. Heat map showing mean ± SEM values for three-four independent experiments, normalized with respect to unstimulated condition taken as 1.

**(J-L)** Receptor surface expression for various assays undertaken in **Figure 5** and **Figure S11** (**J**- corresponding to **Figure 5D**, **I**- corresponding to **Figure S11E**, **J**- corresponding to **Figure S11F, K**-corresponding to **Figure 5E**- right panel and **Figure S11G, L**- corresponding to **Figure 5F**). Surface expression was measured by whole cell-based surface ELISA. Receptor fold-expression was calculated by normalising ELISA and Janus signals of receptor-transfected cells with respect to that of signals recorded for mock-transfected (pcDNA) cells. Data represents mean ± SEM, n>3 independent experiments.

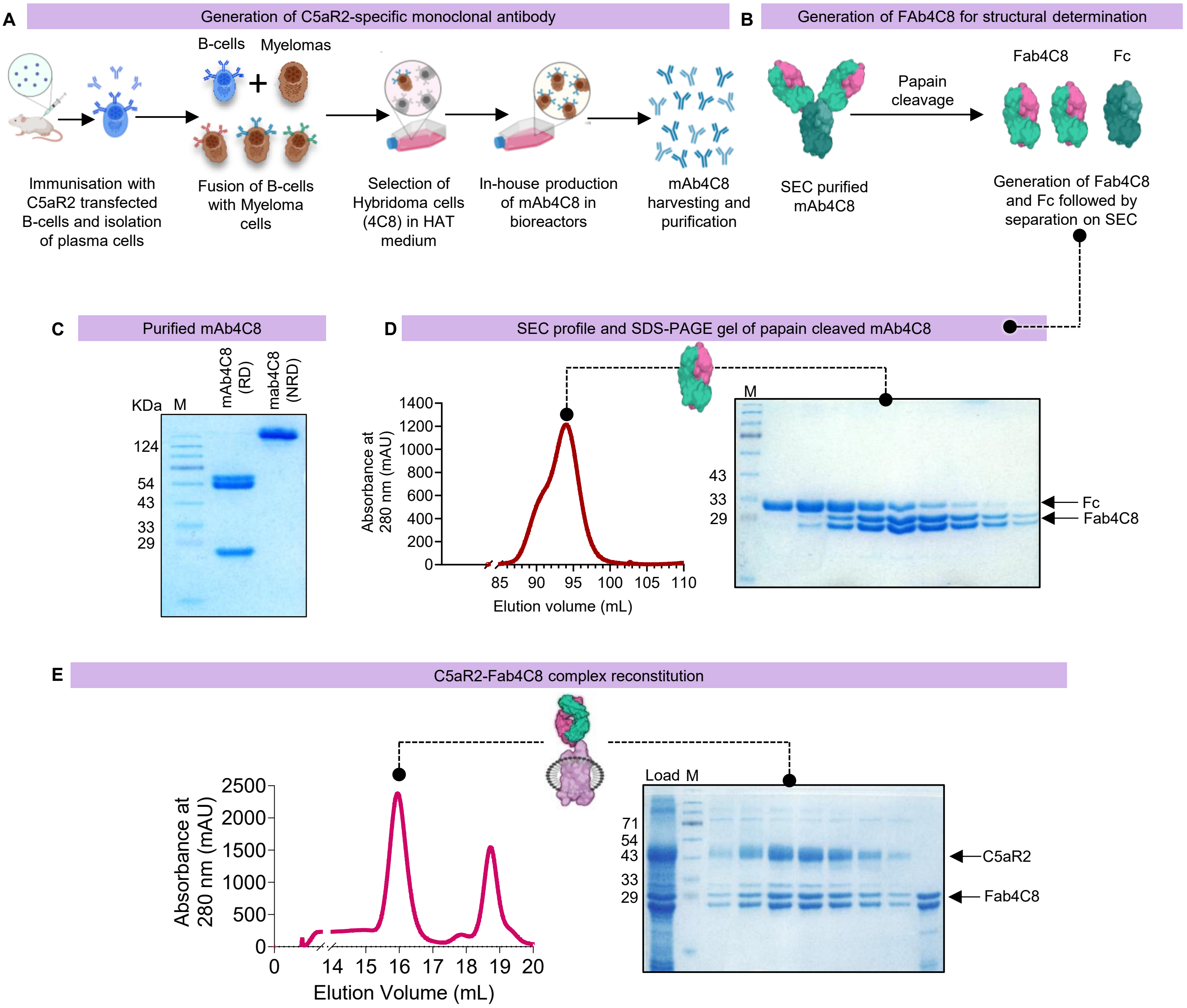


**Figure S11. Synthesis and production of C5aR2-specific blocking monoclonal antibody 4C8 (mAb4C8), related to figure 5F.**

**(A)** Schematic representing the generation and purification of C5aR2-specific antibody. (Schematic for mAb4C8 generation was prepared on the basis of protocol described previously).

**(B)** Schematic representing Fab4C8 generation via papain cleavage of mAb4C8 (produced in-house). All the schematics were prepared in BioRender.

**(C)** SDS-PAGE gel showing heavy and light chains of mAb4C8 under non-reducing (NRD) and reducing (RD) condition.

**(D)** Size-exclusion profile (SEC) and its corresponding SDS-PAGE gel image of Fab4C8. Digested mAb4C8 samples were run on HiLoad column for SEC to separate Fab4C8 and Fc fragments.

**(E)** SEC profile and its corresponding SDS-PAGE gel image of Fab4C8-R8Y-C5aR2 complex.

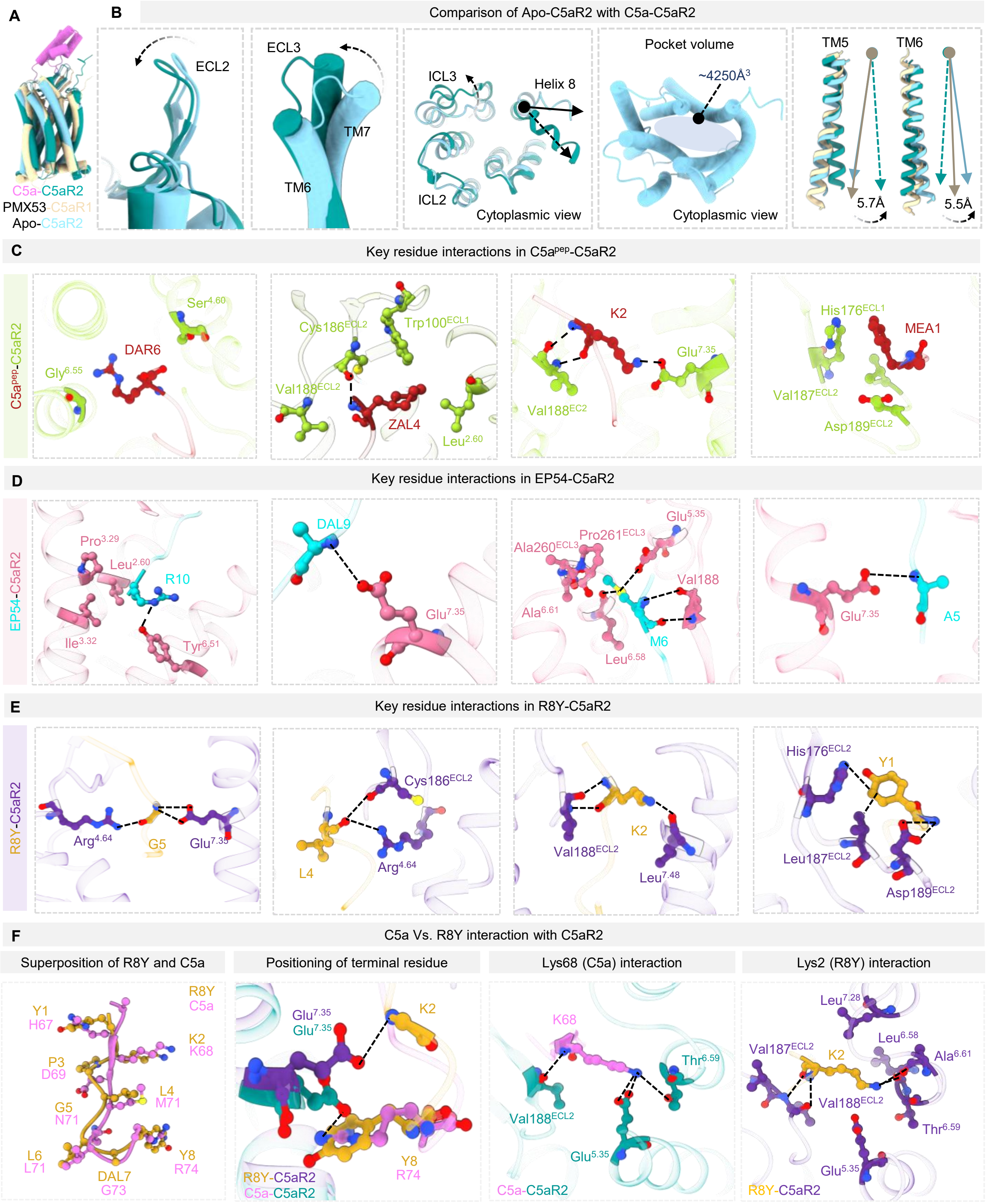
A
B
Comparison of Apo-C5aR2 with C5a-C5aR2
C5a-C5aR2
PMX53-C5aR1
Apo-C5aR2
ECL2
ECL3
TM7
TM6
ICL3
Helix 8
ICL2
Cytoplasmic view
Pocket volume
~4250Å$^3$
Cytoplasmic view
TM5
TM6
5.7Å
5.5Å
C
Key residue interactions in C5a$^{pep}$-C5aR2
C5a$^{pep}$-C5aR2
Ser$^{4.60}$
DAR6
Gly$^{6.55}$
Cys186$^{ECL2}$
Trp100$^{ECL1}$
Val188$^{ECL2}$
ZAL4
Leu$^{2.60}$
K2
Glu$^{7.35}$
Val188$^{EC2}$
His176$^{ECL1}$
MEA1
Val187$^{ECL2}$
Asp189$^{ECL2}$
D
Key residue interactions in EP54-C5aR2
EP54-C5aR2
Pro$^{3.29}$
Leu$^{2.60}$
R10
Ile$^{3.32}$
Tyr$^{6.51}$
DAL9
Glu$^{7.35}$
Glu$^{5.35}$
Pro261$^{ECL3}$
Ala260$^{ECL3}$
Val188
Ala$^{6.61}$
M6
Leu$^{6.58}$
Glu$^{7.35}$
A5
E
Key residue interactions in R8Y-C5aR2
R8Y-C5aR2
Arg$^{4.64}$
G5
Glu$^{7.35}$
Cys186$^{ECL2}$
L4
Arg$^{4.64}$
K2
Val188$^{ECL2}$
Leu$^{7.48}$
His176$^{ECL2}$
Y1
Leu187$^{ECL2}$
Asp189$^{ECL2}$
F
C5a Vs. R8Y interaction with C5aR2
Superposition of R8Y and C5a
Positioning of terminal residue
Lys68 (C5a) interaction
Lys2 (R8Y) interaction
R8Y
C5a
Y1
H67
K2
K68
P3
D69
L4
M71
G5
N71
L6
L71
DAL7
G73
Y8
R74
Glu$^{7.35}$
Glu$^{7.35}$
K2
Y8
R74
R8Y-C5aR2
C5a-C5aR2
K68
Thr$^{6.59}$
Val188$^{ECL2}$
Glu$^{5.35}$
C5a-C5aR2
Leu$^{7.28}$
Leu$^{6.58}$
Val187$^{ECL2}$
K2
Ala$^{6.61}$
Val188$^{ECL2}$
Thr$^{6.59}$
Glu$^{5.35}$
R8Y-C5aR2

**Figure S12. Structural snapshots of C5aR2 interacting with synthetic peptides, related to figure 6.**

**(A)** Superimposition of Apo-C5aR2, C5a-C5aR2 and PMX53-C5aR1 (6C1R).

**(B)** Comparison of Apo-C5aR2 with C5a-C5aR2 showing variability and flexibility in TM regions and loop regions of Apo-C5aR2.

**(C)** Structural snapshots of key residue interactions of $C5a^{pep}$-bound C5aR2.

**(D)** Structural snapshots of key residue interactions of EP54-bound C5aR2.

**(E)** Structural snapshots of key residue interactions of R8Y-bound C5aR2.

**(F)** Structural snapshots comparing overall binding pose and key residue interaction of C5a-bound C5aR2 and R8Y-bound C5aR2.

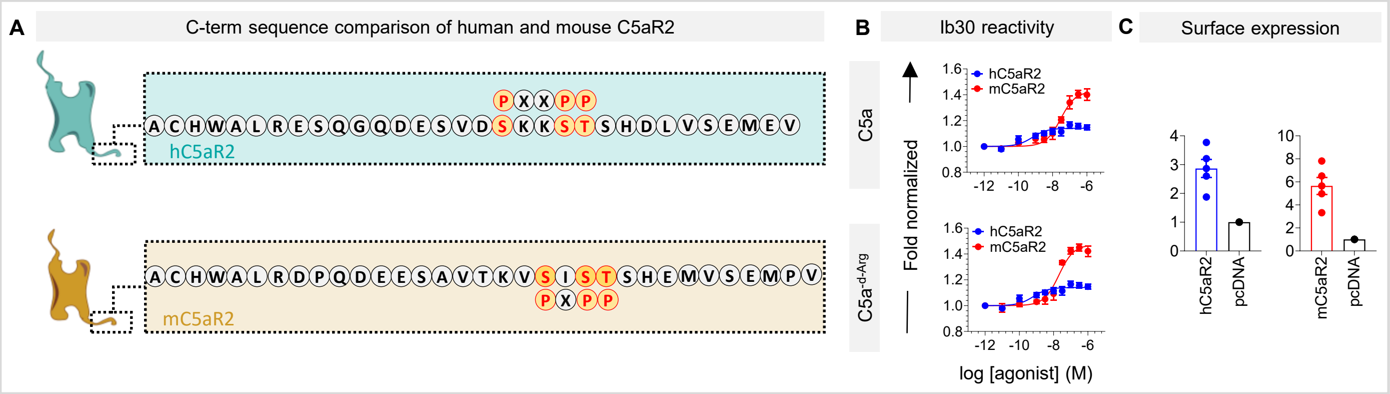


**Figure S13. Ib30 reactivity in human and mouse C5aR2.**

**(A)** Schematic representation of C-terminal of hC5aR2 and mC5aR2 showing former lacks P-X-P-P motif, whereas the latter possess the same. Schematic was prepared in BioRender.

**(B)** Ib30 reactivity downstream to human and mouse C5aR2 in response to human and mouse C5a, respectively. Data represents mean ± SEM, n=5 independent experiments.

**(C)** Receptor surface expression for Ib30 reactivity (related to **Figure S14B**). Surface expression was measured by whole cell-based surface ELISA. Receptor fold-expression was calculated by normalising ELISA and Janus signals of receptor-transfected cells with respect to that of signals recorded for mock-transfected (pcDNA) cells. Data represents mean ± SEM, n=5 independent experiments.

Go surface charge and receptor ICL3 and H8 engagement

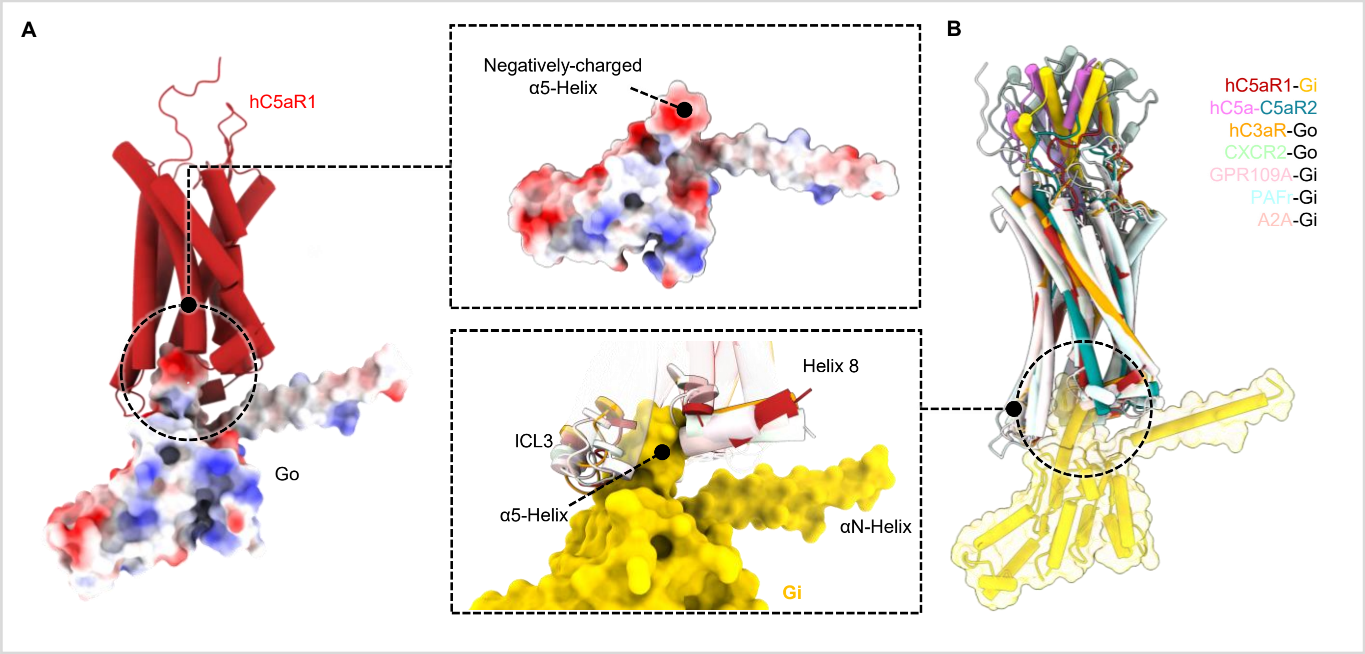


**Figure S14. GPCR and G-protein docking interfaces, related to figure 7.**

**(A)** Structural snapshot of hC5a-hC5aR1-Go (PDB 8IA2), showing C5aR1 in tube helix representation and Go as surface charges, demonstrating the positively charged α5-helix of Go, which docks into a compatible cavity of C5aR1 (C5a not shown).

**(B)** Superposition of GPCR-Go/i structures, viz., hC5a-hC5aR1-Go (PDB: 8IA2), hC5a-C5aR2 (this study), hC3a-hC3aR-Go (PDB: 8I9L), CXCL8-CXCR2-Go (PDB: 8XX6), Niacin-GPR109A-Gi (PDB: 8IY9), PAF-PAFR-Gi (PDB: 8XYD) and Epinephrine-α2AAR-Gi (PDB: 9CBL) superposed on the Go of hC5a-hC5aR1-Go (PDB: 8IA2) structure showing extensive interface of ICL3 and H8 with Gα, signifying the importance of these receptor structural motifs.
